# A chemogenetic platform for controlling plasma membrane signaling and synthetic signal oscillation

**DOI:** 10.1101/2021.03.16.435568

**Authors:** Yuka Hatano, Sachio Suzuki, Akinobu Nakamura, Tatsuyuki Yoshii, Kyoko Atsuta-Tsunoda, Kazuhiro Aoki, Shinya Tsukiji

## Abstract

Chemogenetic methods that enable the rapid translocation of specific signaling proteins in living cells using small molecules are powerful tools for manipulating and interrogating intracellular signaling networks. However, existing techniques rely on chemically induced dimerization of two protein components and have certain limitations, such as a lack of reversibility, bioorthogonality, and usability. Here, by expanding our self-localizing ligand-induced protein translocation (SLIPT) approach, we have developed a versatile chemogenetic system for plasma membrane (PM)-targeted protein translocation. In this system, a novel engineered *Escherichia coli* dihydrofolate reductase in which a hexalysine (K6) sequence is inserted in a loop region (^iK6^DHFR) is used as a universal protein tag for PM-targeted SLIPT. Proteins of interest that are fused to the ^iK6^DHFR tag can be specifically recruited from the cytoplasm to the PM within minutes by addition of a myristoyl-d-Cys-tethered trimethoprim ligand (m^D^cTMP). We demonstrated the broad applicability and robustness of this engineered protein–synthetic ligand pair as a tool for the conditional activation of various types of signaling molecules, including protein and lipid kinases, small GTPases, heterotrimeric G proteins, and second messengers. In combination with a competitor ligand and a culture-medium flow chamber, we further demonstrated the application of the system for chemically manipulating protein localization in a reversible and repeatable manner to generate synthetic signal oscillations in living cells. The present bioorthogonal ^iK6^DHFR/m^D^cTMP-based SLIPT system affords rapid, reversible, and repeatable control of the PM recruitment of target proteins, offering a versatile and easy-to-use chemogenetic platform for chemical and synthetic biology applications.

## INTRODUCTION

Cellular functions are regulated by signaling networks involving proteins, lipids, and other second messenger molecules, and cells precisely coordinate biochemical signaling events in space and time in a fast and reversible manner.^[1,2]^ Recently, single-cell imaging experiments have also revealed that several signaling proteins, such as the extracellular signal-regulated kinase (ERK), show oscillatory activation dynamics and the frequency of the activation pulse determines the cellular responses.^[3,4]^ The ability to manipulate the activity of signaling molecules and pathways in a rapid, reversible, and repeatable manner would therefore be useful for understanding the relationships between signaling input dynamics and cellular outputs, and ultimately for engineering synthetic cellular behaviors.

Methods that enable the rapid localization control or translocation of specific proteins in living cells offer a powerful means for modulating signaling activities. Consequently, several chemogenetic^[5–7]^ and optogenetic^[8,9]^ approaches have been developed as tools for controlling protein translocation. Chemogenetic protein translocation systems are particularly attractive because small-molecule control is easy to perform *in vitro*, *ex vivo*, and *in vivo* to confer rapid temporal modulation. In addition, unlike most optogenetic approaches, chemogenetic methods can be used together with fluorescent reporters of various colors, enabling experiments that combine user-defined perturbation with simultaneous visualization of the subcellular dynamics of multiple signaling activities in the same living cell (“experimental multiplexing”).^[10]^

The most widely used method for chemogenetic protein translocation control relies on a chemically induced dimerization (CID) system using the small-molecule rapamycin, which induces the heterodimerization of the FK506-binding protein (FKBP) and the FKBP-rapamycin-binding (FRB) domain.^[11–13]^ The rapamycin CID system has been proven to be a versatile tool to control the translocation of a wide range of signaling proteins with fast kinetics. However, the rapamycin CID is essentially irreversible.^[14]^ Rapamycin also binds and interferes with the functions of endogenous FKBP and the mammalian target of rapamycin (mTOR), leading to undesired biological effects. To address these issues, several new CID systems have been rationally constructed using two ligand-binding protein tags, including those based on *Escherichia coli* dihydrofolate reductase (eDHFR) and the FKBP F36V mutant,^[15,16]^ SNAP-tag and (wild-type) FKBP,^[17]^ and eDHFR and HaloTag^[16,18–20]^ pairs. In these systems, chimeric molecules consisting of two small-molecule ligands for each protein tag are used as chemical dimerizers. These systems allow reversible protein translocation by the combined use of the chemical dimerizer and a free competitor ligand. However, the protein tag-based CID systems require the careful adjustment of the chemical dimerizer concentrations because excess chemical dimerizer will bind to both the protein components, competitively interfering with the protein heterodimerization. In addition, the chemical dimerizer for the SNAP-tag/FKBP CID system lacks biorthogonality because it also binds to endogenous FKBP.^[17]^ Moreover, as a common limitation shared by all CID tools, dimerization-dependent methods require the coexpression of two proteins with appropriate expression levels and stoichiometry to control the target protein, which is challenging in practice. Therefore, the establishment of chemogenetic protein translocation methods that overcome these limitations is highly desirable.

Here we present a versatile, single protein component, chemogenetic protein translocation system that can be used for manipulating diverse signaling processes at the plasma membrane (PM) based on our self-localizing ligand-induced protein translocation (SLIPT) strategy.^[21–24]^ In this system, a novel engineered eDHFR in which a hexalysine (K6) sequence is inserted in a loop region is used as a universal protein tag for PM-targeted SLIPT. Proteins of interest that are fused to the loop-engineered eDHFR can be rapidly and specifically recruited to the PM upon addition of myristoyl-d-Cys-tethered trimethoprim (m^D^cTMP), a previously developed self-localizing ligand (SL). We show the broad applicability and robustness of this bioorthogonal, loop-engineered eDHFR/m^D^cTMP-based SLIPT system for the conditional activation of various types of signaling molecules, including protein and lipid kinases, small GTPases, heterotrimeric G proteins, and second messengers, such as Ca^2+^ and cAMP. We further demonstrated that the combined use of the present tool and a culture-medium flow system can enable the chemical control of protein localization in a reversible and repeatable manner to produce synthetic signal oscillations in living cells, expanding the repertoire of chemogenetic tools for manipulating intracellular signaling dynamics.

## RESULTS AND DISCUSSION

### Development of a loop-engineered eDHFR tag

The inner leaflet of the PM serves as a platform for intracellular signaling networks, and almost all signaling pathways that determine cell physiology, such as growth, differentiation, phagocytosis, and migration, are initiated and modulated at the PM. Therefore, chemogenetic methods capable of recruiting signaling proteins to the PM are particularly important. Using a bioorthogonal small-molecule trimethoprim (TMP) and eDHFR pair,^[25]^ we have previously developed a PM-targeted SLIPT system (**Figure S1a**).^[22,23]^ In this system, a TMP ligand is conjugated via a flexible linker to a designer myristoyl-d-Cys (myr^D^C) lipopeptide motif to form m^D^cTMP, which is used as an SL (**Figure 1a**).^[23]^ The myr^D^C motif undergoes *S*-palmitoylation of the Cys residue by palmitoyl acyltransferases in cells, which localizes the motif to the PM and the Golgi. In conjunction, an eDHFR variant containing a hexalysine (K6) sequence at the *N*-terminus (K6-eDHFR) is used as a protein tag for fusion to proteins of interest.^[22]^ Non-engineered (wild-type) eDHFR is recruited, not only to the PM, but also undesirably to the Golgi by m^D^cTMP (**Figure S1b** and **Figure S2a**). However, K6-eDHFR is translocated preferentially to the PM by m^D^cTMP because the PM localization of the m^D^cTMP/K6-eDHFR complex is enhanced by electrostatic interactions between the cationic K6 tag and the negatively charged phospholipid phosphatidylserine (PS) present at the inner PM.^[22]^ Consequently, by fusing a protein of interest to the *C*-terminus of the K6-eDHFR tag, the resulting protein can be recruited specifically to the PM upon addition of mDcTMP (**Figure S1c** and **Figure S2b**). This m^D^cTMP/K6-eDHFR-based SLIPT system has been used as a tool for conditional PM-specific protein translocation in live cultured cells and a nematode (*Caenorhabditis elegans*).^22,23^ However, when the K6-eDHFR tag was fused to the *C*-terminus of a protein, such as enhanced green fluorescent protein (EGFP) (EGFP-K6-eDHFR), the protein was translocated to the Golgi in addition to the PM (**Figure S1d** and **Figure S2c**), implying that the PM specificity of K6-eDHFR was insufficient. This result indicated that K6-eDHFR can be used as a PM-specific tag only when it is fused to the *N*-terminus of protein targets, limiting the application of the system.

**Figure 1.**
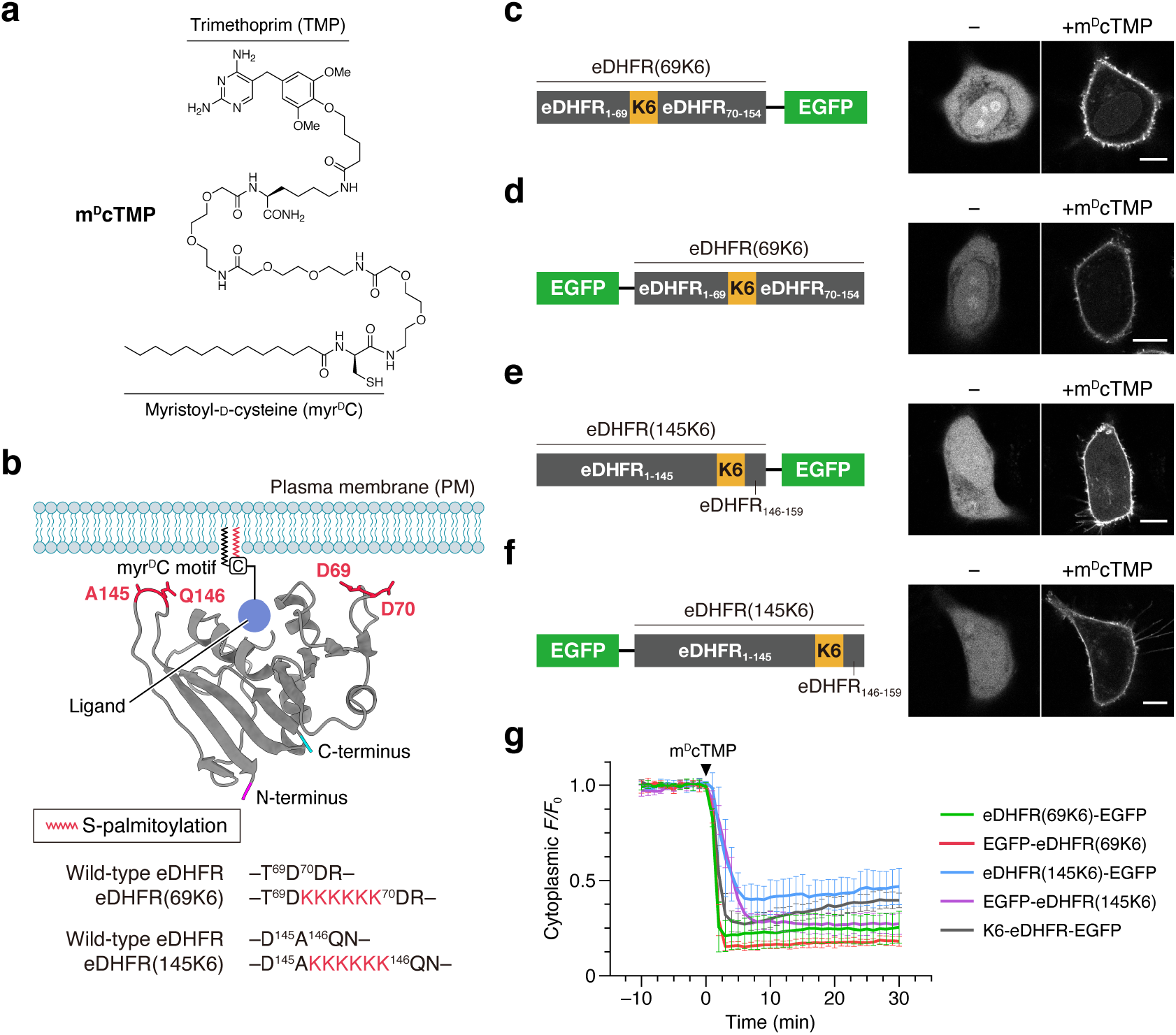
Development of a universal protein tag for PM-specific SLIPT. (**a**) Chemical structure of m^D^cTMP. (**b**) Design of loop-engineered eDHFR constructs based on the topology of PM-anchored eDHFR. In the schematic illustration, the crystal structure of eDHFR complexed with methotrexate (PDB: 1RG7^[26]^) was used and the cartoon model was created with UCSF ChimeraX^[47]^ (methotrexate is not shown). The structure is depicted in a manner such that eDHFR is anchored on the putative PM by m^D^cTMP. The *N*- and *C*-termini are shown in magenta and cyan, respectively. The K6 motif was inserted between D69 and D70 or between A145 and Q146 of wild-type eDHFR to generate eDHFR^69K6^ and eDHFR^145K6^, respectively. (**c**–**f**) m^D^cTMP-induced translocation of eDHFR^69K6^-EGFP (**c**), EGFP-eDHFR^69K6^ (**d**), eDHFR^145K6^-EGFP (**e**), and EGFP-eDHFR^145K6^ (**f**). Confocal fluorescence images of HeLa cells expressing the indicated construct were taken before (left) and 30 min after the addition of m^D^cTMP (10 μM) (right). Scale bars, 10 μm. For the time-lapse movie of EGFP-eDHFR^69K6^ translocation, see **Movie S1**. (**g**) Time course of PM translocation (quantification of data shown in **c**–**f** and **Figure S2**). The normalized fluorescence intensity in the cytoplasm was plotted as a function of time. Data are presented as the mean ± SD (n = 5 cells).

In this work, we aimed to engineer a universal SLIPT tag that can be fused to either the *N*-terminus or *C*-terminus of proteins or even between two proteins or protein domains, while retaining the PM-targeting specificity. To this end, we reconsidered the molecular orientation of the m^D^cTMP/eDHFR complex when it is anchored to the inner surface of the PM. As shown in **Figure 1b**, in the crystal structure of eDHFR (PDB: 1RG7^[26]^), the *N*- and *C*-termini of eDHFR are located at the opposite sides of the ligand-binding pocket. In the previous K6-eDHFR tag, the K6 motif was linked to the *N*-terminus of eDHFR via a 16 amino acid linker such that the K6 sequence could access the PM.^[22]^ Therefore, it is reasonable to consider that when a protein is fused to the *N*-terminus of K6-eDHFR, the access of the K6 tag to the PM will be sterically hindered. This may explain the impaired PM-specificity of the EGFP-K6-eDHFR translocation observed previously. To address this issue, we sought to introduce the K6 sequence to a different site of the eDHFR. On the basis of the model shown in **Figure 1b**, we focused on two loop regions, Ser63–Val72 and Ser135–Ser150. These loops are positioned near the ligand-binding pocket and closely face the inner surface of the PM when the m^D^cTMP/eDHFR complex is in the PM-anchored orientation. We therefore decided to insert the K6 motif into the loop regions of eDHFR, and constructed two variants, eDHFR^69K6^ and eDHFR^145K6^, which contained the K6 motif between Asp69 and Asp70, and Ala145 and Gln146, respectively (**Figure 1b**).

Prior to performing SLIPT assays in cells, we measured the binding affinity of the eDHFR variants to TMP *in vitro*. The dissociation constants (*K*_d_) of wild-type eDHFR, K6-eDHFR, eDHFR^69K6^, and eDHFR^145K6^ to fluorescein-conjugated TMP (TMP-FL) were determined by fluorescence polarization measurements to be 18, 24, 106, and 108 nM, respectively (**Figure S3**). Although the binding affinity to TMP was decreased approximately 6-fold by the insertion of the K6 motif into the loop regions, the loop-engineered eDHFR^69K6^ and eDHFR^145K6^ still maintained sub-micromolar affinity to TMP.

We tested the ability of the loop-engineered eDHFR variants to act as protein tags for SLIPT. We first fused eDHFR^69K6^ to the *N*-terminus of EGFP (eDHFR^69K6^-EGFP) and expressed the construct in human epithelial HeLa cells. Before ligand treatment, eDHFR^69K6^-EGFP was distributed over the entire cell. The addition of m^D^cTMP induced PM-specific translocation of eDHFR^69K6^-EGFP with almost no observable Golgi localization (**Figure 1c** and **Figure S2d**). Furthermore, when we fused eDHFR^69K6^ to the *C*-terminus of EGFP (EGFP-eDHFR^69K6^), this protein was also recruited specifically to the PM in response to m^D^cTMP (**Figure 1d, Figure S2e**, and **Movie S1**). Similarly, the fusion of eDHFR^145K6^ to the *N*- or *C*-terminus of EGFP achieved PM-specific translocation by m^D^cTMP (**Figure 1e,f** and **Figure S2f,g**). These results demonstrated that both loop-engineered eDHFR^69K6^ and eDHFR^145K6^ functioned as PM-specific SLIPT tags that can be fused to either the *N*- or *C*-termini of proteins. The times required for half-maximal PM recruitment (*t*_1/2_) of eDHFR^69K6^-EGFP and EGFP-eDHFR^69K6^ (at 10 μM m^D^cTMP) were 1.4 and 1.5 min, respectively, which were almost comparable with that of K6-DHFR-EGFP (*t*_1/2_ = 1.7 min) (**Figure 1g**). In contrast, the m^D^cTMP-induced PM recruitment times for eDHFR^145K6^-EGFP and EGFP-eDHFR^145K6^ (*t*_1/2_ = 2.5 and 3.2 min, respectively) were slightly slower compared with the eDHFR^69K6^ counterparts. On the basis of these results, we selected eDHFR^69K6^ as a universal protein tag for m_D_cTMP-induced PM-specific protein translocation, which is hereafter denoted as ^iK6^DHFR (internally K6-tagged eDHFR). The ^iK6^DHFR tag also induced PM-specific recruitment of the EGFP fusion protein in NIH3T3 and Cos-7 cells (**Figure S4**), demonstrating the general applicability of the m^D^cTMP/^iK6^DHFR SLIPT system to various cell lines.

### Mechanistic characterization of the m^D^cTMP/eDHFR^iK6^ SLIPT system

Next, we investigated which membrane lipids contributed to the PM-targeting specificity of the ^iK6^DHFR tag. The inner PM is highly anionic compared with other organelle membranes because of the presence of the negatively charged phospholipids, such as PS,^[27,28]^ and phosphatidylinositol 4-phosphate (PI4P) and phosphatidylinositol-4,5-bisphosphate [PI(4,5)P_2_]^[28–30]^. On the basis of our previous work,^[22]^ we performed depletion experiments of these anionic lipid species. After recruitment of EGFP-^iK6^DHFR to the PM by m^D^cTMP, we reduced the PS concentration in the inner PM by the addition of ionomycin.^[27,28]^ As a result, EGFP-^iK6^DHFR dissociated from the PM and was relocalized to the endomembrane (**Figure S5a,b**). In contrast, when we depleted PM PI4P and PI(4,5)P_2_ by the PM recruitment of Pseudojanin, a chimeric protein of Sac1 (PI4P phosphatase) and INPP5E [PI(4,5)P_2_ phosphatase], using a rapamycin CID tool,^[29]^ no noticeable dissociation of the PM-localized EGFP-^iK6^DHFR was observed (**Figure S5c–e**). These results suggested that the interaction of the K6 motif in ^iK6^DHFR with the PM PS plays a dominant role in controlling the PM-specific localization of the m^D^cTMP/^iK6^DHFR system.

In nature, many PM-localized proteins, such as MARCS and KRas4B, possess a single lipid modification (myristoylation or prenylation) and a polybasic domain for PM targeting.^[27–30]^ This fact raises the possibility that the m^D^cTMP/^iK6^DHFR complex, which contains a myristoyl lipid anchor and the K6 motif, might be able to localize to the PM without the *S*-palmitoylation of mDcTMP. We thus evaluated whether the *S*-palmitoylation of the D^-^Cys moiety of mDcTMP was required for the m^D^cTMP-induced PM localization of ^iK6^DHFR. When cells expressing EGFP-^iK6^DHFR were treated with myrTMP lacking the palmitoylatable Cys moiety,^[31]^ EGFP-^iK6^DHFR was still recruited to the PM but was also localized undesirably to the endomembrane region (**Figure S6a,b**). This result indicated that the d-Cys moiety of m^D^cTMP is not necessarily essential for PM localization but is critical for achieving the efficient and PM-specific translocation of ^iK6^DHFR-fusion proteins.

### Chemogenetic activation of the Raf/ERK signaling pathway

We applied the m^D^cTMP/^iK6^DHFR SLIPT system to manipulate intracellular signaling. We first focused on cRaf, which is a protein kinase that regulates diverse cell functions, including growth, survival, and differentiation.^[32]^ When cRaf is recruited to the PM, it is activated by autophosphorylation, which triggers the activation of downstream MEK and ERK. To construct a synthetic cRaf protein, the localization of which could be controlled by mDcTMP, full-length cRaf was fused to the *C*-terminus of EGFP-^iK6^DHFR to generate EGFP-^iK6^DHFR-cRaf (**Figure 2a**). We then expressed this protein in HeLa cells. To monitor the activity of endogenous ERK, we coexpressed a kinase translocation reporter (KTR) for ERK, which was fused to monomeric Kusabira Orange (mKO) (ERK-KTR-mKO).^[22]^ In the KTR system, the endogenous ERK activity can be quantified by the ratio of the cytoplasmic to the nuclear fluorescence intensity (C/N ratio) of the reporter protein.^[33,34]^ Prior to the addition of mDcTMP, EGFP-^iK6^DHFR-cRaf showed a cytoplasmic distribution (**Figure 2b**). When cells were treated with m^D^cTMP, EGFP-^iK6^DHFR-cRaf translocated to the PM within minutes (*t*_1/2_ = 1.2 min) (**Figure 2c** and **Movie S2**). Concomitant with this cRaf translocation, an increase in the C/N ratio (indicative of nuclear export of ERK-KTR-mKO) was observed, demonstrating efficient activation of endogenous ERK (**Figure 2b**). The C/N ratio increase reached a plateau approximately 20 min after mDcTMP addition (**Figure 2c**). Such ERK activation was not observed when we performed the same experiment in the presence of PD184352, a MEK inhibitor (**Figure S7a**), or using EGFP-^iK6^DHFR lacking the cRaf protein (**Figure S7b**). These results demonstrated that the m^D^cTMP/^iK6^DHFR SLIPT system can be used for synthetic activation of the Raf/ERK signaling pathway. Furthermore, the above experiment showed that the loop-engineered ^iK6^DHFR functions as a PM-specific SLIPT tag even when it is inserted between two protein domains (in this case, EGFP and cRaf).

**Figure 2.**
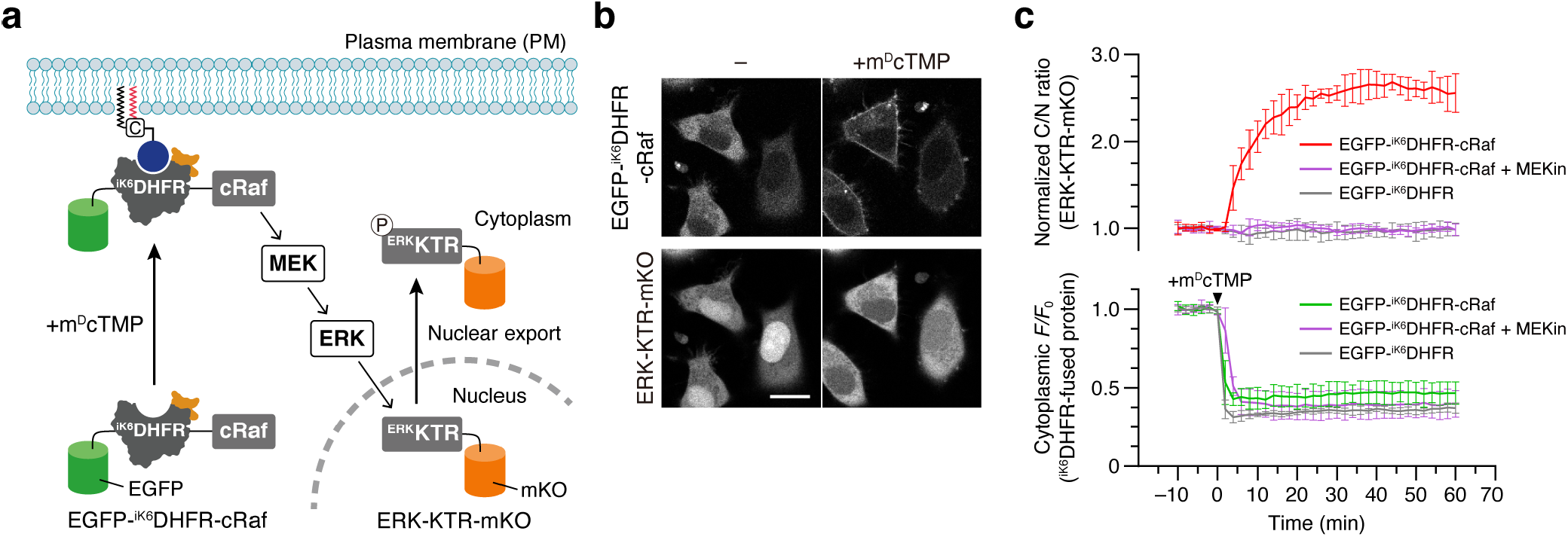
Chemogenetic activation of the Raf/ERK pathway. (**a**) Schematic illustration of the experimental setup. (**b**) Confocal fluorescence images of HeLa cells coexpressing EGFP-^iK6^DHFR-cRaf and ERK-KTR-mKO were taken before (left) and 60 min after the addition of m^D^cTMP (10 μM) (right). Scale bar, 20 μm. For the time-lapse movie, see **Movie S2**. (**c**) Time course of EGFP-^iK6^DHFR-cRaf translocation and ERK activation. To evaluate EGFP-^iK6^DHFR-cRaf translocation (bottom), the normalized fluorescence intensity of EGFP-^iK6^DHFR-cRaf in the cytoplasm was plotted as a function of time. To evaluate ERK activity (top), the normalized ratios of the cytoplasmic fluorescence intensity to the nuclear fluorescence intensity (C/N ratios) of ERK-KTR-mKO were plotted as a function of time. Data from control experiments using EGFP-^iK6^DHFR-cRaf in the presence of the MEK inhibitor PD184352 (**Figure S7a**) or EGFP-^iK6^DHFR (lacking the cRaf protein) (**Figure S7b**) were plotted on the same graph. Data are presented as the mean ± SD (n = 5 cells).

### Chemogenetic control of heterotrimeric G proteins and second messenger signaling

Heterotrimetric G proteins are an important class of signaling molecules regulated downstream of G protein-coupled receptors (GPCRs). In particular, there are several Gα isoforms, including Gαq, Gαs, Gα_12/13_, and Gαi, and more than one of these isoforms are often activated simultaneously upon GPCR stimulation.^[35,36]^ Thus, it has been difficult to study the contribution of each isoform to cellular functions. In a pioneering work, Putyrski and Schultz have developed conditional Gα activation systems using the rapamycin CID technique.^[37]^ However, chemogenetic methods applicable to control heterotrimeric G proteins are still limited. We therefore tested the utility of the present SLIPT system for manipulating G protein signaling. First, we targeted Gαq. The Gαq protein is an activator of phospholipase Cβ (PLCβ), and activation of Gαq triggers Ca^2+^ release from the endoplasmic reticulum (ER) to the cytoplasm, and subsequent Ca^2+^ oscillations.^[38,39]^ To control Gαq localization and activity using mDcTMP, a palmitoylation-deficient constitutively active form of Gαq was fused at the *C*-terminus to mNeonGreen-tagged ^iK6^DHFR (mNG-^iK6^DHFR-Gαq) and expressed in HeLa cells (**Figure 3a**). To evaluate the intracellular Ca^2+^ concentration, we coexpressed mNG-^iK6^DHFR-Gαq and R-GECO, a genetically encoded red fluorescent Ca^2+^ indicator.^[40]^ When cells coexpressing mNG-^iK6^DHFR-Gαq and R-GECO were treated with m^D^cTMP, mNG-^iK6^DHFR-Gαq was rapidly translocated to the PM, which induced a series of Ca^2+^ spikes in the cells (**Figure 3b,c** and **Movie S3**). In contrast, PM recruitment of a PLCβ-binding-deficient Gαq mutant did not induce Ca^2+^ oscillations (**Figure S8**). Therefore, the applicability of the m^D^cTMP/^iK6^DHFR SLIPT system to control Gαq protein and Ca^2+^ signaling was demonstrated.

**Figure 3.**
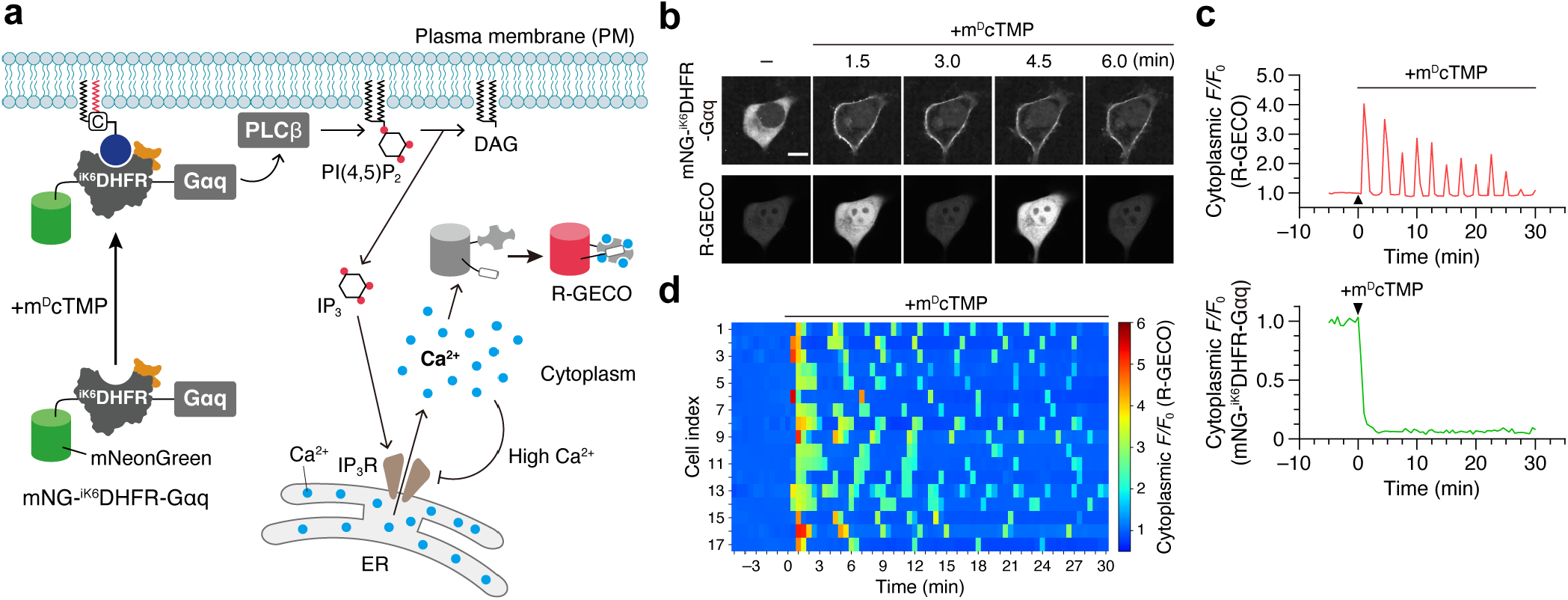
Chemogenetic activation of Gαq signaling and Ca^2+^ oscillations. (**a**) Schematic illustration of the experimental setup. (**b**) Representative time-lapse confocal fluorescence images of a HeLa cell coexpressing mNG-^iK6^DHFR-Gαq and R-GECO. Images were taken before and after the addition of m^D^cTMP (5 μM). Scale bar, 10 μm. For the time-lapse movie, see **Movie S3**. (**c**) The time course of mNG-^iK6^DHFR-Gαq translocation and Ca^2+^ spikes observed in the cell is shown in panel **b**. The normalized fluorescence intensities of mNG-^iK6^DHFR-Gαq (bottom) and R-GECO (top) in the cytoplasm were plotted as a function of time. (**d**) Heatmaps depicting m^D^cTMP-induced Ca^2+^ oscillations for 17 randomly selected cells coexpressing mNG-^iK6^DHFR-Gαq and R-GECO.

Furthermore, it was also possible to apply the system to control various signaling proteins that are naturally regulated by receptor tyrosine kinases and GPCRs, including Gαs (cAMP production) (**Figure S9** and **Movie S4**), RasGEF (Ras activation) (**Figure S10** and **Movie S5**), PI3K (phosphatidylinositol 3,4,5-trisphosphate [PI(3,4,5)P3] production) (**Figure S11** and **Movie S6**), and Tiam1 (Rac activation and lamellipodium induction) (**Figure S12** and **Movie S7**), verifying the broad applicability and robustness of the system as a chemogenetic platform for manipulating intracellular signaling processes at the PM.

### Chemical induction of synthetic signal oscillations

Reversibility is an important characteristic of cell signaling events. Recent single-cell analyses have revealed that several signaling proteins, including p53, ERK, and NF-kB, exhibited oscillatory dynamics to govern cell fate decisions.^[3,4,41–43]^ Therefore, the ability to artificially induce oscillatory dynamics of cell signaling in living cells would be highly valuable for understanding the mechanisms of the dynamical encoding by these signaling molecules. This ability requires methods that allow protein activity pulses to be manipulated in a reversible and repeatable manner. To date, such oscillatory control of protein activity has been achieved only by optogenetics,^[41,44–46]^ and chemogenetic approaches for this purpose have not yet been well established. Hence, we attempted to apply our m^D^cTMP/^iK6^DHFR SLIPT system to the oscillatory (multi-cycle) control of protein activity in living cells.

Our strategy was as follows (**Figure 4a**). First, an ^iK6^DHFR-tagged signaling protein is recruited to the PM by m^D^cTMP, inducing the first synthetic signal activation. The protein is then returned to the cytoplasm by treating the cells with excess free TMP (as a competitor ligand), terminating the first signal pulse. Subsequently, by a washing procedure, the free TMP is removed from the cells, whereas m^D^cTMP (and its palmitoylated form) remains in the cells because of its high affinity with the membranes. Consequently, the ^iK6^DHFR-tagged protein is relocalized to the PM. Therefore, by repeating the TMP addition and washing steps, following the initial m^D^cTMP addition, we should be able to reversibly and repeatedly induce the PM–cytoplasm shuttling of the protein with the desired temporal control.

**Figure 4.**
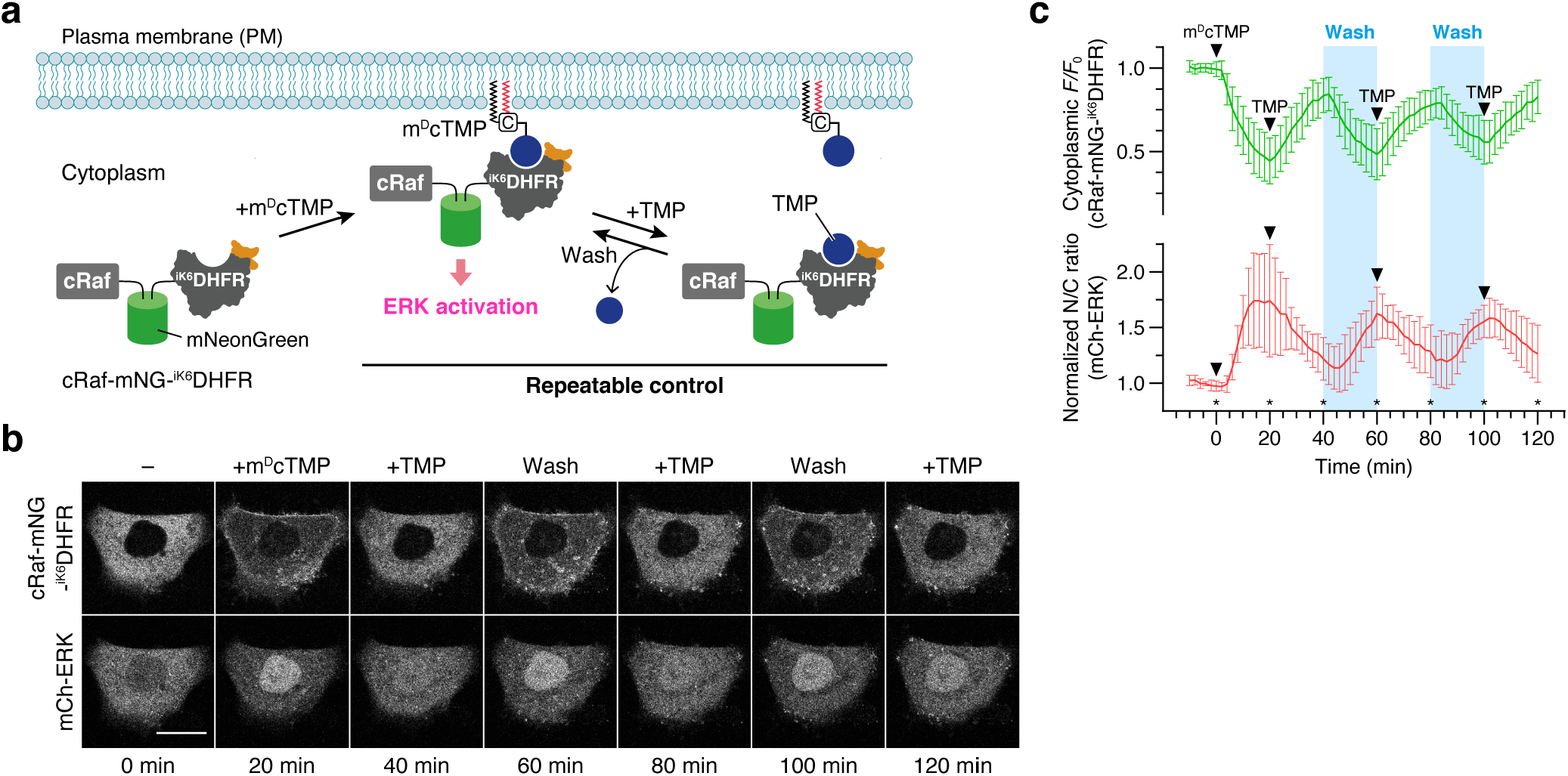
Chemical induction of synthetic ERK signal oscillation. (**a**) Schematic illustration of the experimental setup. The washing procedure was carried out using a culture-medium flow system as shown in **Figure S13a**. (**b**) Representative time-lapse confocal fluorescence images of a HeLa cell coexpressing cRaf-mNG-^iK6^DHFR and mCh-ERK. Images were taken at the time points indicated by the asterisks shown in panel **c**. m^D^cTMP and TMP were added at concentrations of 10 and 50 μM, respectively. During the washing step (the blue bar in panel **c**), fresh medium continuously flowed at a rate of 1 mL/min. Scale bar, 20 μm. For the time-lapse movie, see **Movie S8**. (**c**) Time course of cRaf-mNG-^iK6^DHFR translocation and ERK activity. To evaluate cRaf-mNG-^iK6^DHFR translocation (top), the normalized fluorescence intensity of cRaf-mNG-^iK6^DHFR in the cytoplasm was plotted as a function of time. To evaluate ERK activity (bottom), the normalized ratios of the nuclear fluorescence intensity to the cytoplasmic fluorescence intensity (N/C ratios) of mCh-ERK were plotted as a function of time. Data are presented as the mean ± SD (n = 12 cells).

To test this strategy, HeLa cells expressing ^iK6^DHFR-EGFP were plated on a culture dish equipped with a culture-medium flow system (**Figure S13a**). First, the cells were treated with mDcTMP (10 μM) to induce the PM translocation of ^iK6^DHFR-EGFP (**Figure S13b**). Following this treatment, the cells were incubated with excess TMP (50 μM), which caused ^iK6^DHFR-EGFP to be returned to the cytoplasm. After 20 min, the cells were washed with flesh medium using the flow system. As expected, we observed the relocalization of ^iK6^DHFR-EGFP to the PM (**Figure S13b**). When we repeated the cycle of TMP addition and washout, the localization of ^iK6^DHFR-EGFP oscillated between the cytoplasm and the PM (**Figure S13b,c**). This experiment demonstrated the proof of principle that the combined use of the m^D^cTMP/^iK6^DHFR SLIPT system, free TMP, and a medium flow system can enable the user-defined oscillatory control and single-cell imaging of reversible PM–cytoplasm protein translocation.

Finally, we investigated the chemical control of ERK signal oscillations using the system established above. For this purpose, we used HeLa cells coexpressing cRaf-mNG-^iK6^DHFR and mCherry-tagged ERK (mCh-ERK) (**Figure 4a**). The cells were first treated with mDcTMP, followed by repeated TMP addition and washing cycles. Using this procedure, ERK activity was reversibly switched on and off synchronously with the chemically controlled shuttling of cRaf-mNG-^iK6^DHFR between the PM and the cytoplasm (**Figure 4b,c** and **Movie S8**). Therefore, we successfully demonstrated the applicability of the present SLIPT system to the chemogenetic control of synthetic ERK signal oscillations.

## CONCLUSION

In this work, we have developed a universal protein tag for PM-specific SLIPT, ^iK6^DHFR, by inserting a K6 motif into a loop region of eDHFR. The ^iK6^DHFR tag can be fused to either the *N*-terminus or *C*-terminus of proteins, and even between two proteins or protein domains, without the loss of its PM specificity, offering a high degree of flexibility in the construction of ^iK6^DHFR fusion proteins. The ^iK6^DHFR-tagged proteins can be rapidly recruited to the PM on the order of minutes in the presence of mDcTMP. We demonstrated the applicability of the m^D^cTMP/^iK6^DHFR SLIPT system to control a wide range of signaling proteins with different sizes (ca. 24 to 73 kDa) and physiological functions, including cRaf, Gαq, Gαs, RasGEF, PI3K, and Tiam1, verifying the versatility and general utility of the system. From the viewpoint of protein engineering, the design of ^iK6^DHFR reported in this work is a successful example of creating a chemogenetic protein tag that interfaces with a specific cellular membrane, i.e., the PM, by loop engineering.

In combination with a competitor ligand (free TMP) and a culture-medium flow chamber, we further demonstrated the application of the m^D^cTMP/^iK6^DHFR SLIPT system to control cRaf localization and ERK signaling activity in a reversible and repeatable manner. To the best of our knowledge, this work is the first report of chemically generated synthetic oscillations in cell signaling. In the ERK signal oscillation system, the forward (signal ON) and reverse (signal OFF) maximum translocation rates of synthetic cRaf (cRaf-mNG-^iK6^DHFR) were 7.1 (*t*_1/2_) and 8.3 min (*t*_rev1/2_), respectively. We were able to chemically generate consecutive synthetic ERK activity pulses three times in 2 h with a periodicity of ca. 40 min. Based on previous work, stochastic ERK activation induced by noise and cell-to-cell propagation occurs with a frequency of approximately 10–20 pulses per day depending on the cell density, which corresponds to an averaged periodicity of ca. 1–2 h.^[41]^ Therefore, the SLIPT-based synthetic signal oscillation system will be useful to reproduce the frequency of ERK activity pulses observed in cultured cells for biological research. It should also be noted that the signal duration time and frequency of signal activation can further be modulated by the timing of the TMP addition and washing steps.

In conclusion, the bioorthogonal m^D^cTMP/^iK6^DHFR-based SLIPT system, which is easy to use and enables the rapid, reversible, and repeatable chemical manipulation of protein localization and signaling processes at the PM, will offer a powerful and versatile chemogenetic platform to interrogate dynamic signaling networks and engineer cell functions for mammalian synthetic biology applications.

## ACKNOWLEDGMENTS

We thank Dr. K. Kuwata (Nagoya University) for HRMS measurement of fluorescein-conjugated TMP. This work was supported by JSPS Grants-in-Aid for Scientific Research (KAKENHI) (Grant Nos. JP15H05949 “Resonance Bio”, JP18H02086, and JP18H04546 and JP20H04706 “Chemistry for Multimolecular Crowding Biosystems”), the Uehara Memorial Foundation, and the Takeda Science Foundation (to S.T.). This work was also supported by JST PRESTO (JPMJPR178B) and MEXT Leading Initiative for Excellent Young Researchers (to T.Y.). S.S. acknowledges scholarship support from the Hirota Scholarship Society and the SUNBOR Scholarship from the Suntory Foundation for Life Sciences. A.N. is a recipient of the JSPS Research fellowship for Young Scientists (JP19J01341).

