## Supplementary Information for "A chemogenetic platform for controlling plasma membrane signaling and synthetic signal oscillation"

### Supplementary Figures

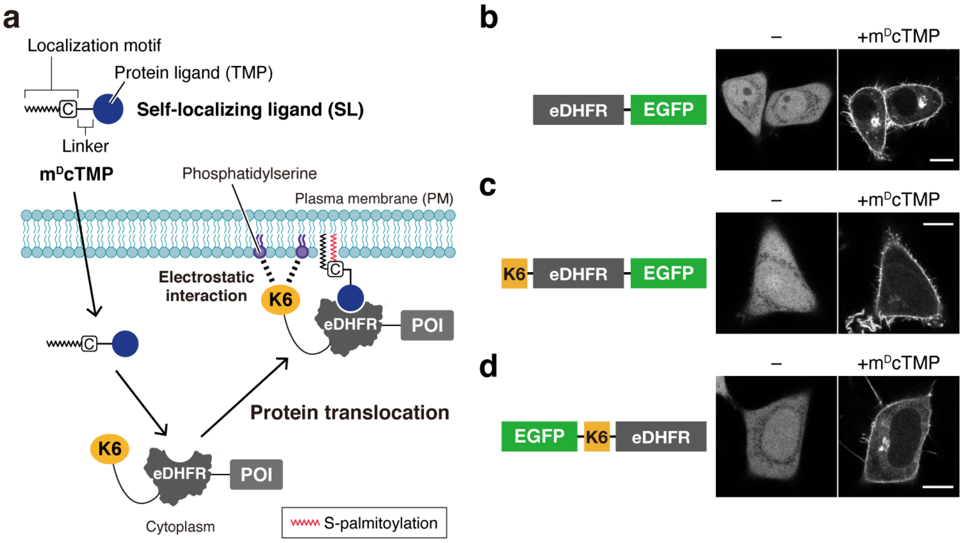


**Figure S1.** The previously reported PM-specific SLIPT system based on the K6-eDHFR tag and m$^{D}$cTMP. (**a**) Schematic illustration of the system. (**b**–**d**) m$^{D}$cTMP-induced translocation of eDHFR-EGFP (**b**), K6-eDHFR-EGFP (**c**), and EGFP-K6-eDHFR (**d**). Confocal fluorescence images of HeLa cells expressing the indicated constructs were taken before (left) and 30 min after the addition of m$^{D}$cTMP (10 µM) (right). Scale bars, 10 µm.

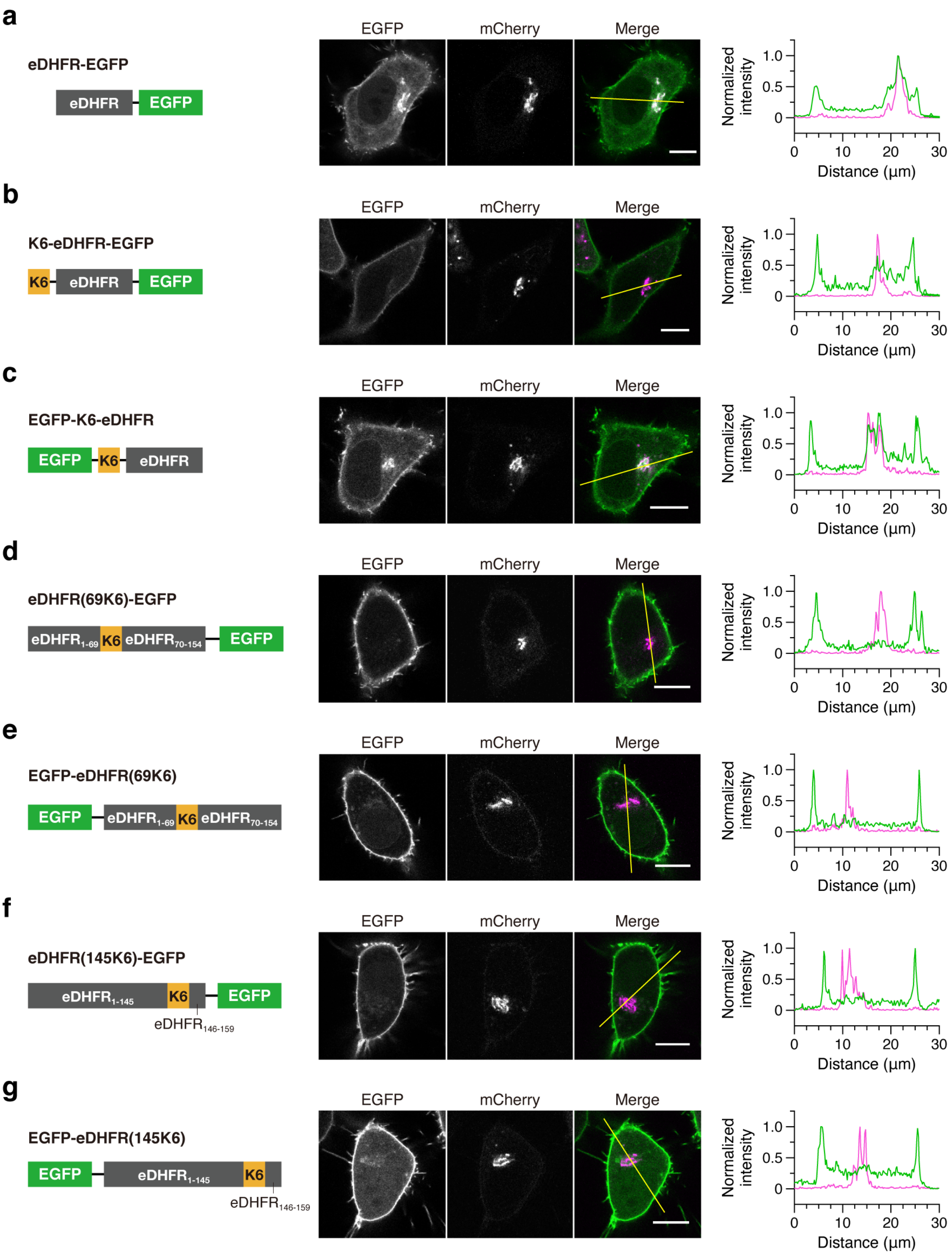


**Figure S2.** Colocalization analysis of the PM-targeting specificity of m$^{D}$cTMP-induced protein translocation. Confocal fluorescence images of HeLa cells coexpressing the indicated constructs and mCherry-Giantin (Golgi marker) were obtained 30 min after incubation with m$^{D}$cTMP (10 µM). The fluorescence intensity profiles of EGFP (green) and mCherry (magenta) channels across the yellow line are shown in the right panel. Scale bars, 10 µm.

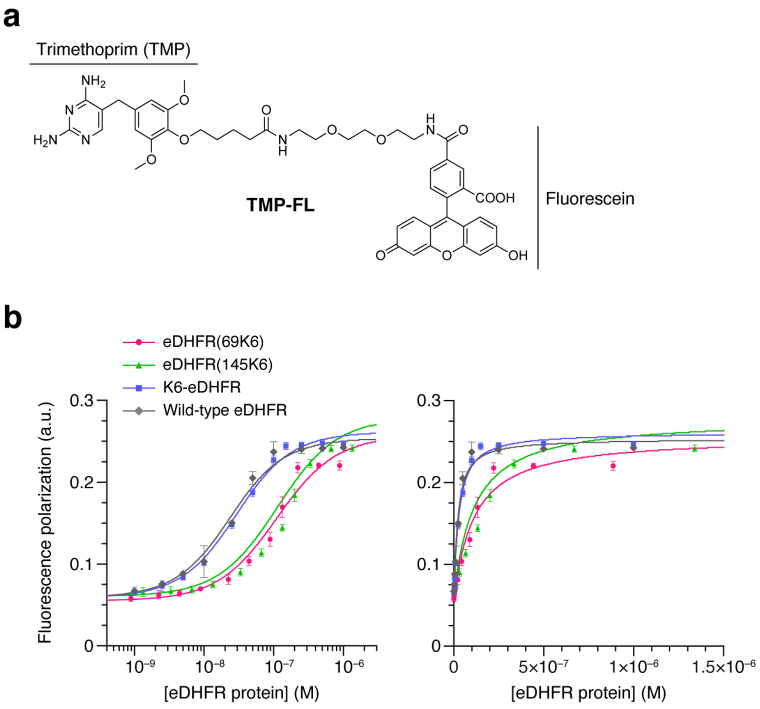


c

| Proteins | $K_d$ (nM) |
|---|---|
| eDHFR(69K6) | 106 ± 10 |
| eDHFR(145K6) | 108 ± 1 |
| K6-eDHFR | 24 ± 3 |
| Wild-type eDHFR | 18 ± 3 |

**Figure S3.** Binding affinity assays of wild-type and engineered eDHFR proteins. (**a**) Chemical structure of TMP-FL. (**b**) Fluorescence polarization experiments. TMP-FL (10 nM) was titrated with varying concentrations of wild-type or engineered eDHFR proteins in the presence of a 50-fold excess of NADPH. The fluorescence polarization values were plotted as a function of the protein concentration, and the binding affinities were determined by fitting the titration data to a 1:1 binding model as described in the "**Binding affinity assays**" section (p. S20). Data are presented as the mean ± SD (n = 3). (**c**) The dissociation constants ($K_d$) of wild-type and engineered eDHFR proteins with TMP-FL determined by fluorescence polarization assays.

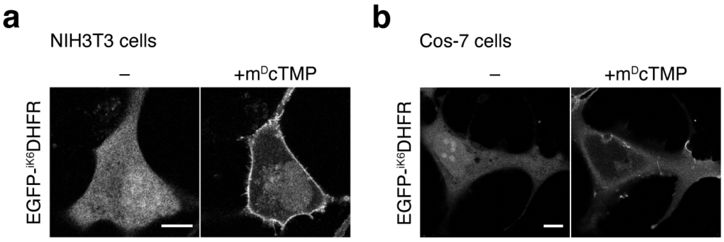


**Figure S4.** EGFP-$^{iK6}$DHFR translocation in other cell lines. Confocal fluorescence images of NIH3T3 cells (**a**) and Cos-7 cells (**b**) expressing EGFP-$^{iK6}$DHFR were taken before (left) and 30 min after the addition of m$^{D}$cTMP (10 μM) (right). Scale bars, 10 μm.

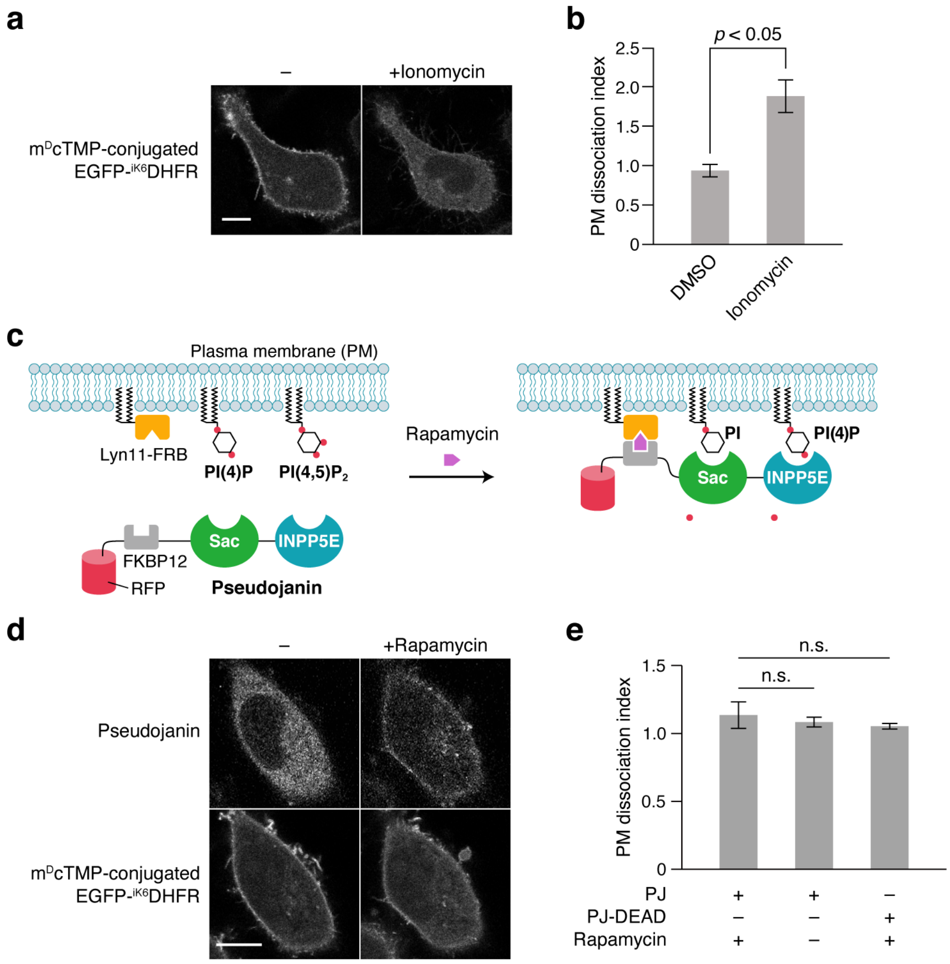


**Figure S5.** The loop-inserted K6 tag of $^{iK6}$DHFR interacted mainly with PS on the inner leaflet of the PM. (**a**) PS depletion experiment. HeLa cells expressing EGFP-$^{iK6}$DHFR were pre-treated with m$^{D}$cTMP (10 µM) for 30 min to recruit the protein to the PM. Confocal fluorescence images of the cells were then taken before (left) and 10 min after incubation with ionomycin (10 µM) (right). Scale bar, 10 µm. (**b**) Quantification of the PM dissociation of the m$^{D}$cTMP/EGFP-$^{iK6}$DHFR complex by PS depletion. The PM dissociation index is given by $(F_{cyt}/F_{PM})/(F_{cyt}/F_{PM})_0$, where $(F_{cyt}/F_{PM})$ and $(F_{cyt}/F_{PM})_0$ are the ratios of the cytoplasmic fluorescence intensity to the PM fluorescence intensity after and before lipid depletion, respectively. Data are presented as the mean ± SD (n = 5 cells). (**c**) Schematic illustration of the rapamycin CID-based PI4P/PI(4,5)$P_2$ depletion system.[S1] (**d**) PI4P/PI(4,5)$P_2$ depletion experiment. HeLa cells co-expressing EGFP-$^{iK6}$DHFR, Pseudojanin, and Lyn11-FRB were pre-treated with m$^{D}$cTMP (10 µM) for 30 min to recruit the protein to the PM. Confocal fluorescence images of the cells were then taken before (left) and 10 min after incubation with rapamycin (200 nM) (right). Scale bar, 10 µm. (**e**) Quantification of the PM dissociation of the m$^{D}$cTMP/EGFP-$^{iK6}$DHFR complex by PI4P/PI(4,5)$P_2$ depletion. The PM dissociation index was estimated as described in **b**. Inactivated Pseudojanin (PJ-DEAD)[S1] was used as a control. Data are presented as the mean ± SD (n = 5 cells).

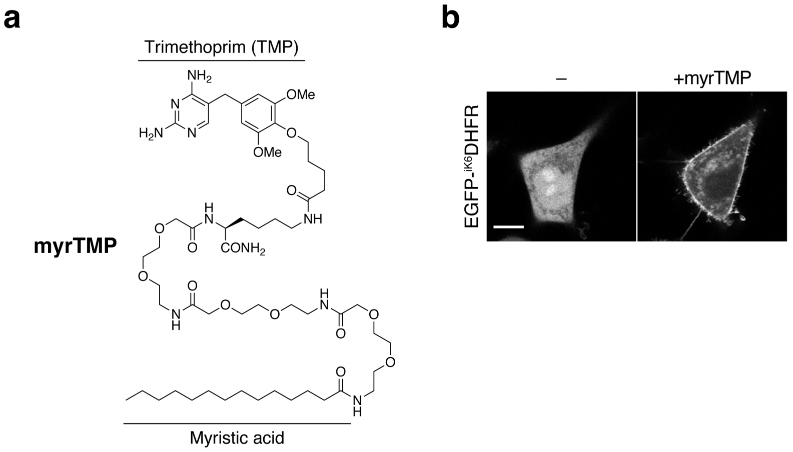


**Figure S6.** Investigation of the effect of m$^{D}$cTMP palmitoylation on the PM-targeted $^{iK6}$DHFR recruitment ability. (**a**) Chemical structure of myrTMP, an unpalmitoylatable m$^{D}$cTMP derivative lacking the $^{D}$Cys residue.[S2] (**b**) Confocal fluorescence images of HeLa cells expressing EGFP-$^{iK6}$DHFR were taken before (left) and 30 min after incubation with myrTMP (10 μM) (right). Scale bar, 10 μm.

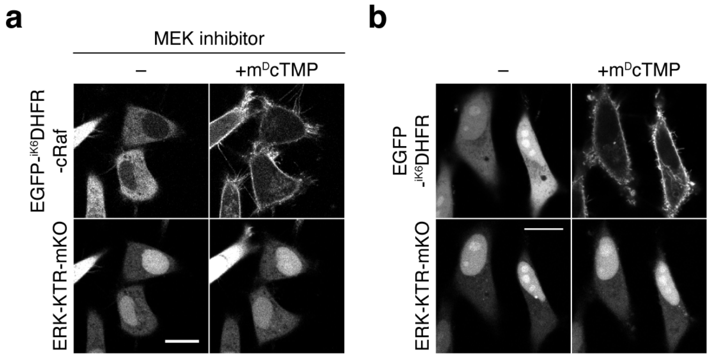


**Figure S7.** Control experiments for the chemogenetic ERK activation shown in **Figure 2**. (**a**) PM recruitment of EGFP-eDHFR$^{iK6}$-cRaf in the presence of the MEK inhibitor PD184352 (30 μM). (**b**) PM recruitment of EGFP-eDHFR$^{iK6}$ lacking cRaf. Confocal fluorescence images of HeLa cells coexpressing the indicated constructs were taken before (left) and 60 min after the addition of m$^D$cTMP (10 μM) (right). Scale bars, 20 μm.

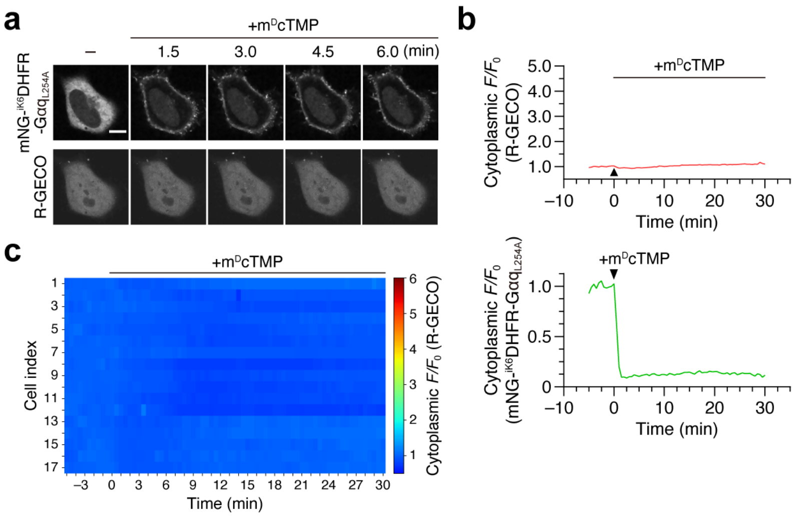


**Figure S8.** Control experiment for the chemogenetic activation of $Ca^{2+}$ oscillations shown in **Figure 3**. (**a**) Representative time-lapse confocal fluorescence images of a HeLa cell coexpressing mNG-$^{iK6}$DHFR-Gαq$_{L254A}$ (carrying a PLCβ-binding-deficient Gαq mutant) and R-GECO. Images were taken before and after the addition of m$^{D}$cTMP (5 μM). Scale bar, 10 μm. (**b**) Time course of the mNG-$^{iK6}$DHFR-Gαq$_{L254A}$ translocation and $Ca^{2+}$ spikes observed in the cell shown in panel **a**. The normalized fluorescence intensities of mNG-$^{iK6}$DHFR-Gαq$_{L254A}$ (bottom) and R-GECO (top) in the cytoplasm were plotted as a function of time. (**c**) Heatmaps depicting $Ca^{2+}$ oscillations for 17 randomly selected cells coexpressing mNG-$^{iK6}$DHFR-Gαq$_{L254A}$ and R-GECO.

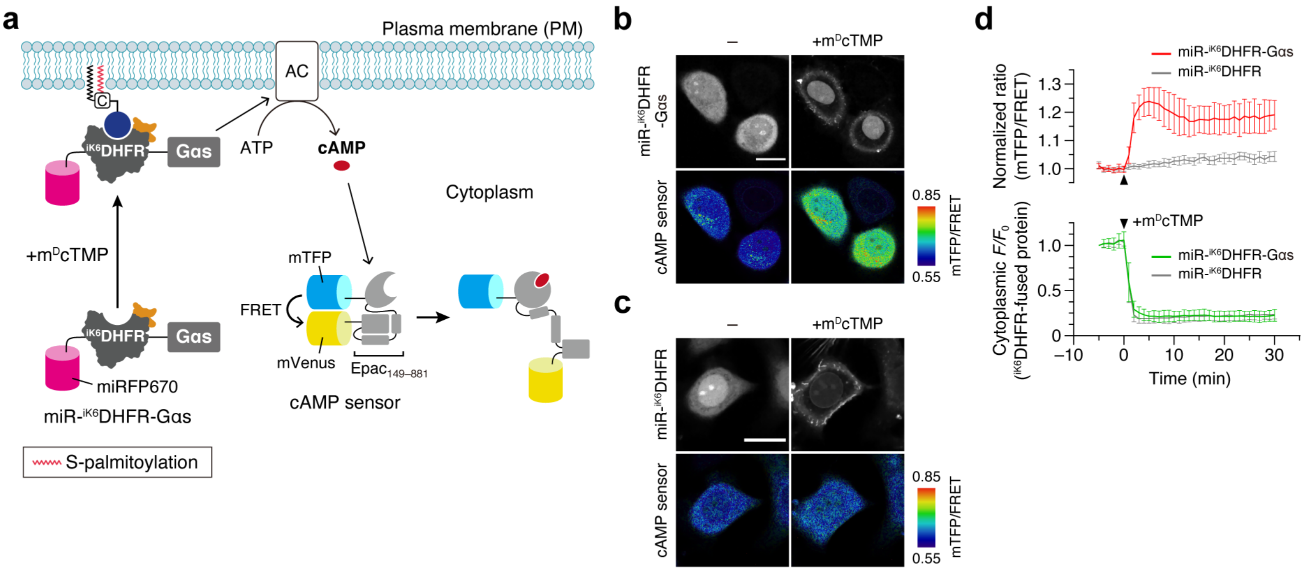


**Figure S9.** Chemogenetic control of Gαs signaling and cAMP production. (**a**) Schematic illustration of the experimental setup. A synthetic, m$^{\mathrm{D}}$cTMP-responsive Gαs protein was constructed by fusing a palmitoylation-deficient Gαs mutant (C3S)[S3] to the *C*-terminus of miRFP670-tagged $^{\mathrm{iK6}}$DHFR (miR-$^{\mathrm{iK6}}$DHFR-Gαs). This protein was used to activate endogenous adenylate cyclase (AC) upon addition of m$^{\mathrm{D}}$cTMP. The intracellular cAMP levels were monitored using a FRET sensor for cAMP.[S4,S5] (**b**) Confocal fluorescence images of HeLa cells coexpressing miR-$^{\mathrm{iK6}}$DHFR-Gαs and the cAMP sensor taken before (left) and 30 min after the addition of m$^{\mathrm{D}}$cTMP (10 μM) (right). For the time-lapse movie, see **Movie S4**. (**c**) PM recruitment of mNG-$^{\mathrm{iK6}}$DHFR lacking the Gαs protein (control experiment for **b**). For **b** and **c**, scale bars = 20 μm. (**d**) Time course of SLIPT and cAMP production. To evaluate SLIPT (bottom), the normalized fluorescence intensities of miR-$^{\mathrm{iK6}}$DHFR-Gαs (or mNG-eDHFR$^{\mathrm{iK6}}$) in the cytoplasm were plotted as a function of time. To evaluate the cAMP levels (top), the normalized ratios of the mTFP fluorescence intensity to the FRET (mVenus) fluorescence intensity (mTFP/FRET ratios) of the cAMP sensor were plotted as a function of time. Data are presented as the mean ± SD (n = 20 cells).

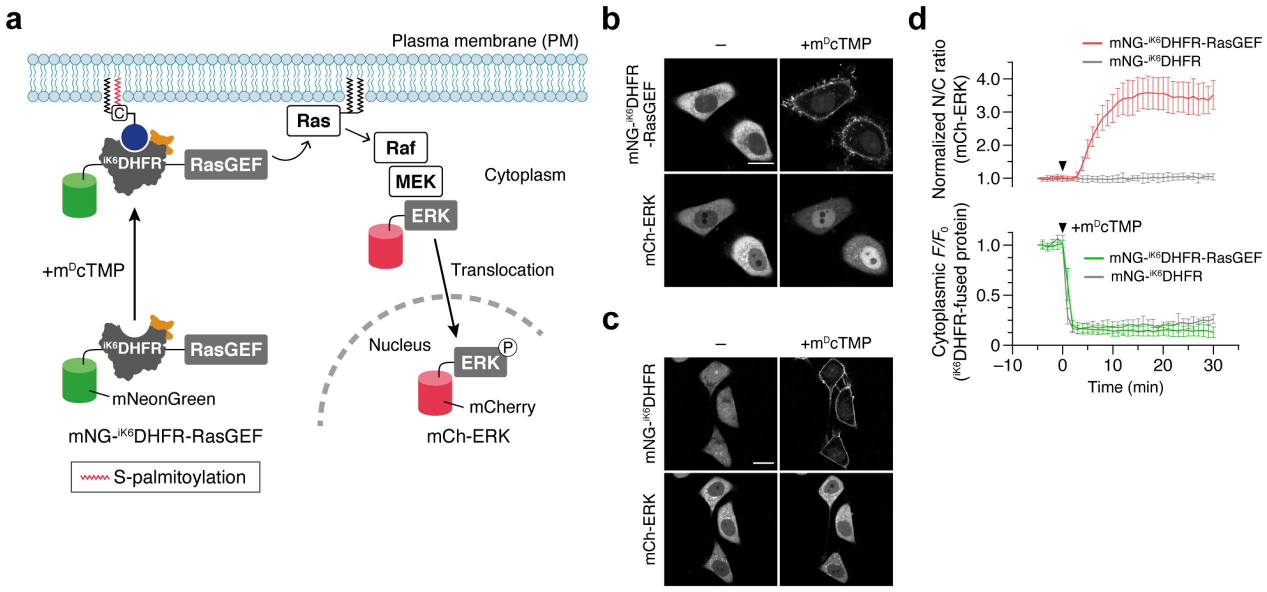


**Figure S10.** Chemogenetic activation of the Ras/ERK pathway. (**a**) Schematic illustration of the experimental setup. A synthetic, $m^{D}$cTMP-responsive guanine nucleotide exchange factor for Ras (RasGEF) was constructed by fusing the catalytic Cdc25 domain of RasGRF1[S6] to the *C*-terminus of mNG-$^{iK6}$DHFR (mNG-$^{iK6}$DHFR-RasGEF). This protein was used to activate endogenous Ras at the PM upon addition of $m^{D}$cTMP. The activity of the ERK pathway was monitored by mCherry-ERK (mCh-ERK). (**b**) Confocal fluorescence images of HeLa cells coexpressing mNG-$^{iK6}$DHFR-RasGEF and mCh-ERK were taken before (left) and 30 min after the addition of $m^{D}$cTMP (10 μM) (right). For the time-lapse movie, see **Movie S5**. (**c**) PM recruitment of mNG-$^{iK6}$DHFR lacking the RasGEF domain (control experiment for **b**). For **b** and **c**, scale bars = 20 μm. (**d**) Time course of SLIPT and ERK activation. To evaluate SLIPT (bottom), the normalized fluorescence intensities of mNG-$^{iK6}$DHFR-RasGEF (or mNG-$^{iK6}$DHFR) in the cytoplasm were plotted as a function of time. To evaluate ERK activity (top), the normalized ratios of the nuclear fluorescence intensity to the cytoplasmic fluorescence intensity (N/C ratios) of mCh-ERK were plotted as a function of time. Data are presented as the mean ± SD (n = 6 cells).

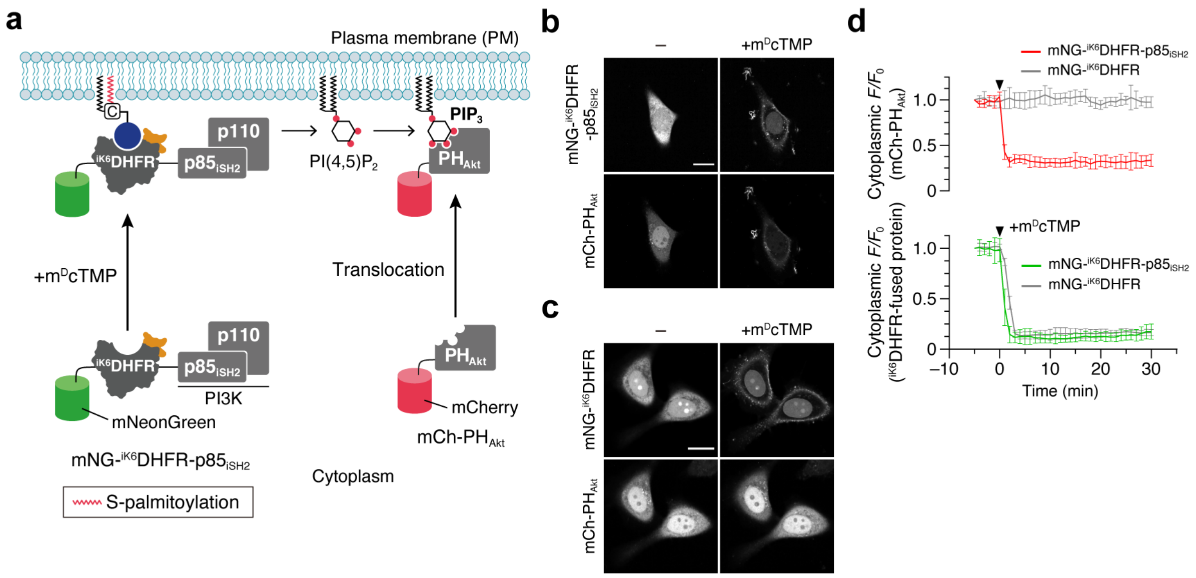


**Figure S11.** Chemogenetic control of PI(3,4,5)$P_3$ production. (**a**) Schematic illustration of the experimental setup. A synthetic, m$^D$cTMP-responsive phosphatidylinositol 3-kinase (PI3K) was constructed by fusing the inter-Src homology 2 (iSH2) domain of the p85 subunit of PI3K to the *C*-terminus of mNG-$^{iK6}$DHFR (mNG-$^{iK6}$DHFR- p85$_{iSH2}$). The iSH2 domain forms a complex with the endogenous p110 subunit of PI3K in cells.[S7,S8] mNG-$^{iK6}$DHFR- p85$_{iSH2}$ was used to produce PI(3,4,5)$P_3$ via phosphorylation of PI(4,5)$P_2$ at the PM upon addition of m$^D$cTMP. Production of PI(3,4,5)$P_3$ was monitored by the pleckstrin homology (PH) domain of Akt fused to mCherry (mCh-PH$_{Akt}$).[S9] (**b**) Confocal fluorescence images of HeLa cells coexpressing mNG-$^{iK6}$DHFR-p85$_{iSH2}$ and mCh-PH$_{Akt}$ were taken before (left) and 30 min after the addition of m$^D$cTMP (10 µM) (right). For the time-lapse movie, see **Movie S6**. (**c**) PM recruitment of mNG-$^{iK6}$DHFR lacking the iSH2 domain (control experiment for **b**). For **b** and **c**, scale bars = 20 µm. (**d**) Time course of SLIPT and PI(3,4,5)$P_3$ production. The normalized fluorescence intensities of mNG-$^{iK6}$DHFR- p85$_{iSH2}$ (or mNG-$^{iK6}$DHFR) (bottom) and mCh-PH$_{Akt}$ (top) in the cytoplasm were plotted as a function of time. Data are presented as the mean ± SD (n = 8 cells).

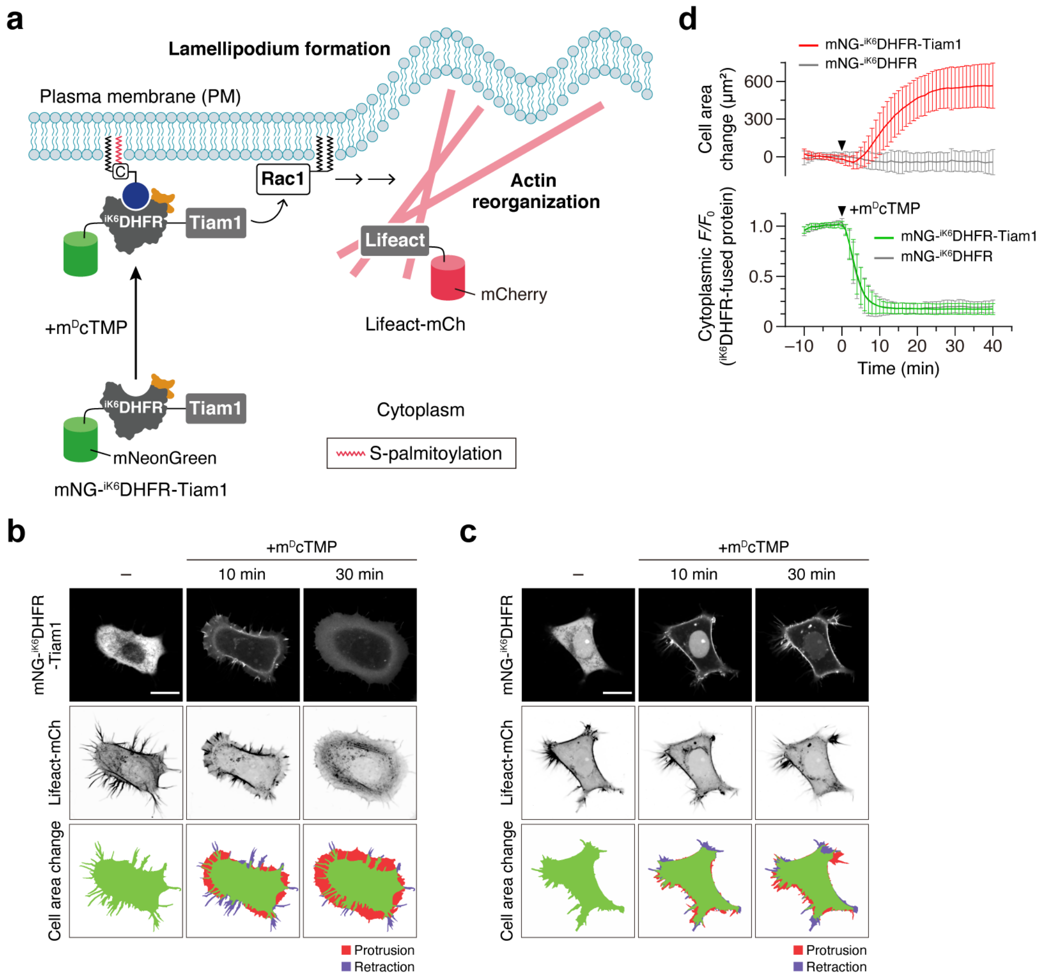


**Figure S12.** Chemogenetic control of Rac activation and lamellipodia formation. (**a**) Schematic illustration of the experimental setup. A synthetic, m$^{D}$cTMP-responsive guanine nucleotide exchange factor for Rac was constructed by fusing the DH-PH domain of Tiam1 to the *C*-terminus of mNG-$^{iK6}$DHFR (mNG-$^{iK6}$DHFR-Tiam1).[S10] This protein was used to activate endogenous Rac upon addition of m$^{D}$cTMP. The cell morphology was monitored by the F-actin marker Lifeact fused to mCherry (Lifeact-mCh).[S11] (**b**) Representative time-lapse confocal fluorescence images of a HeLa cell coexpressing mNG-$^{iK6}$DHFR-Tiam1 and Lifeact-mCh. Images were taken before (left) and after the addition of m$^{D}$cTMP (5 μM) (right). For the time-lapse movie, see **Movie S7**. (**c**) PM recruitment of mNG-$^{iK6}$DHFR lacking the Tiam1 domain (control experiment for **b**). For **b** and **c**, scale bars = 20 μm. (**d**) Time course of SLIPT and cell area change. To evaluate SLIPT (bottom), the normalized fluorescence intensities of mNG-$^{iK6}$DHFR-Tiam1 (or mNG-$^{iK6}$DHFR) in the cytoplasm were plotted as a function of time. Cell areas were calculated from the Lifeact-mCh images, and cell area changes were plotted as a function of time. Data are presented as the mean ± SD (n = 40 cells for mNG-$^{iK6}$DHFR-Tiam1 and 24 cells for mNG-$^{iK6}$DHFR).

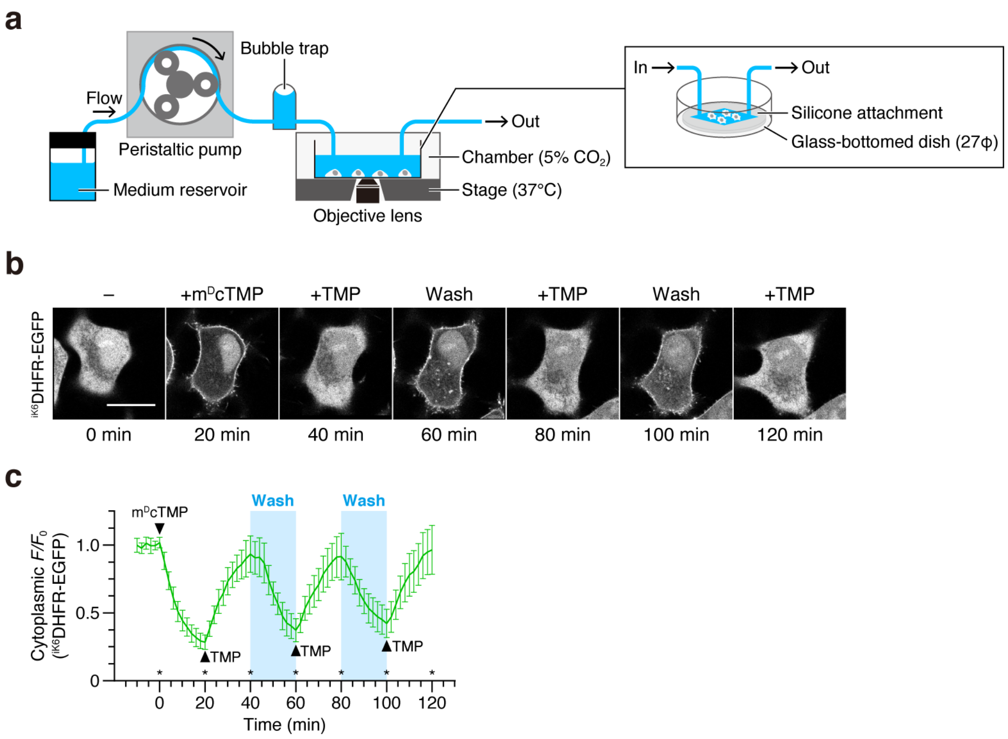


**Figure S13.** Chemogenetic control of protein translocation oscillations. (**a**) Schematic illustration of the culture-medium flow system. (**b**) Representative time-lapse confocal fluorescence images of a HeLa cell expressing $^{iK6}$DHFR-EGFP. Images were taken at the time points indicated by the asterisks shown in panel **c**. m$^{D}$cTMP and TMP were added at concentrations of 10 µM and 50 µM, respectively. During the washing step (the blue bar in panel **c**), fresh medium continuously flowed at a rate of 1 mL/min. Scale bar, 20 µm. (**c**) Time course of repeated $^{iK6}$DHFR-EGFP translocation. The normalized fluorescence intensity of $^{iK6}$DHFR-EGFP in the cytoplasm was plotted as a function of time. Data are presented as the mean ± SD (n = 12 cells).

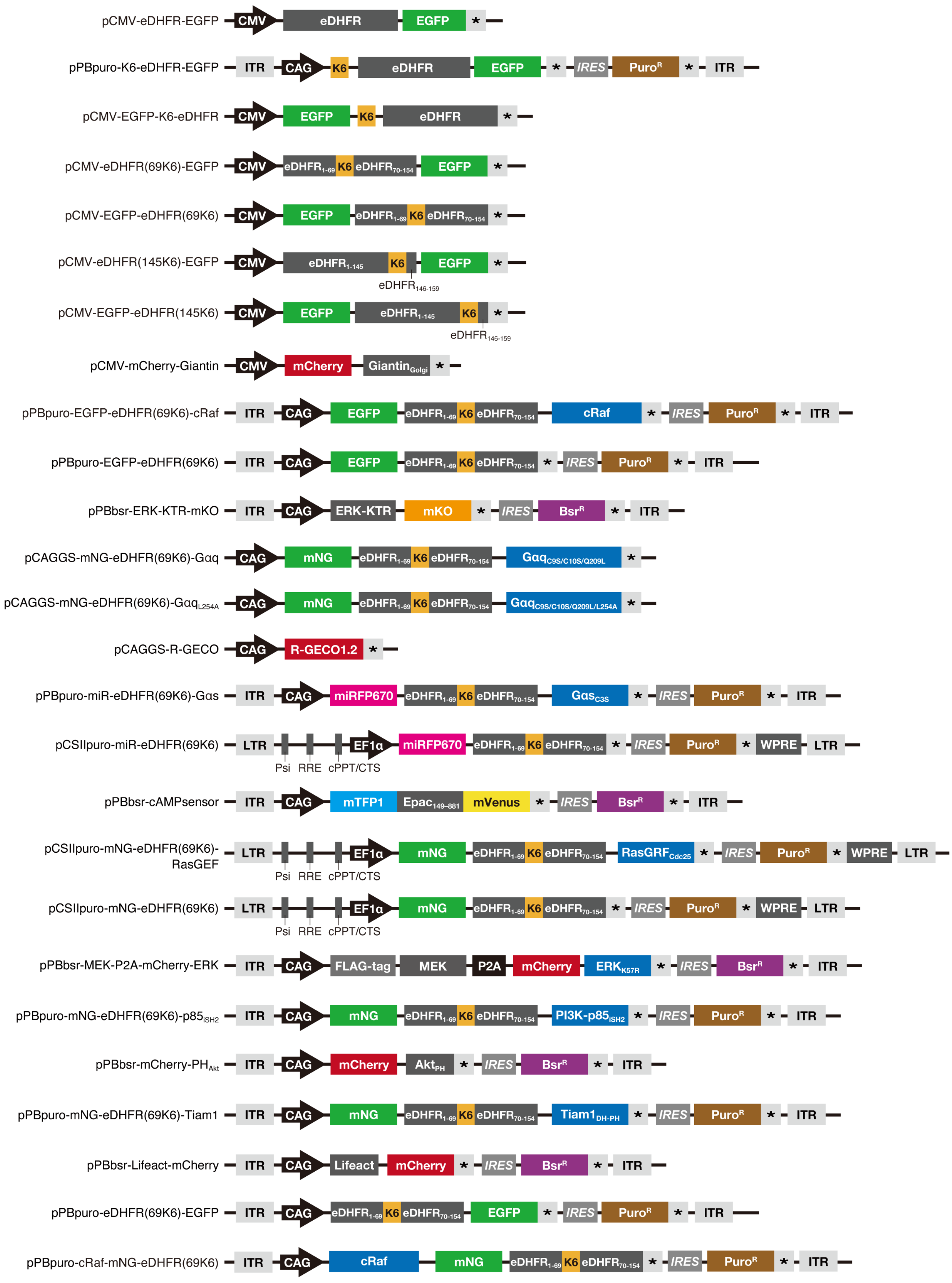


**Figure S14.** Schematic illustration of the domain structures of constructs used in this study. DNA and amino acid sequences of the constructs are shown in the "**Supplementary Sequences**" section.

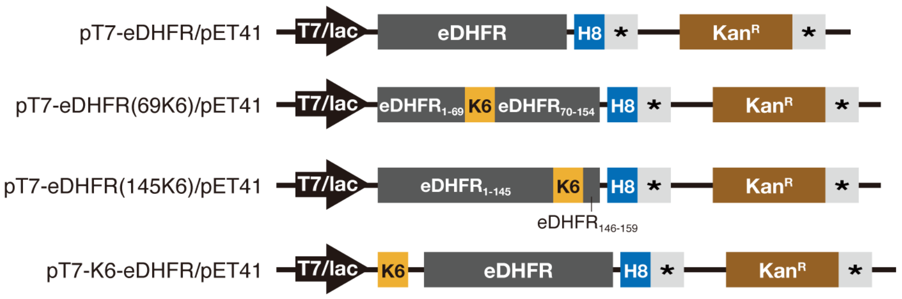


**Figure S15.** Schematic illustration of the domain structures of constructs used for bacterial expression. All vectors encoded aminoglycoside phosphotransferase ($Kan^R$) as a kanamycin resistance gene. DNA and amino acid sequences of the constructs are shown in the "**Supplementary Sequences**" section.

#### Supplementary Movies

**Movie S1.** m$^{D}$cTMP-induced PM recruitment of EGFP-eDHFR$^{69K6}$ in HeLa cells (time-lapse movie of **Figure 1d**). Scale bar, 10 µm.

**Movie S2.** Chemogenetic activation of the Raf/ERK pathway by PM recruitment of EGFP-$^{iK6}$DHFR-cRaf (time-lapse movie of **Figure 2b**). Scale bar, 10 µm.

**Movie S3.** Chemogenetic activation of Gαq signaling and Ca$^{2+}$ oscillations by PM recruitment of mNG-$^{iK6}$DHFR-Gαq (time-lapse movie of **Figure 3b**). Scale bar, 10 µm.

**Movie S4.** Chemogenetic control of Gαs signaling and cAMP production by PM recruitment of miR-$^{iK6}$DHFR-Gαs (time-lapse movie of **Figure S9b**). Scale bar, 20 µm.

**Movie S5.** Chemogenetic activation of the Ras/ERK pathway by PM recruitment of mNG-$^{iK6}$DHFR-RasGEF (time-lapse movie of **Figure S10b**). Scale bar, 20 µm.

**Movie S6.** Chemogenetic control of PI(3,4,5)P$_3$ production by PM recruitment of mNG-$^{iK6}$DHFR-p85$_{iSH2}$ (time-lapse movie of **Figure S11b**). Scale bar, 20 µm.

**Movie S7.** Chemogenetic control of Rac activation and lamellipodia formation by PM recruitment of mNG-$^{iK6}$DHFR-Tiam1 (time-lapse movie of **Figure S12b**). Scale bar, 20 µm.

**Movie S8.** Chemical induction of synthetic ERK signal oscillations by repeated PM-cytoplasm shuttling of cRaf-mNG-$^{iK6}$DHFR (time-lapse movie of **Figure 4b**). Scale bar, 20 µm.

### Supplementary Methods: Chemical Synthesis

#### General materials and methods

All chemical reagents and solvents were purchased from commercial supplies (Watanabe Chemical Industries, Tokyo Chemical Industry, FUJIFILM Wako Pure Chemical Corp., and Kanto Chemical) and used without further purification. Thin layer chromatography (TLC) was performed on silica gel 60 $F_{254}$ precoated aluminum sheets (Merck), and TLC plates were visualized by fluorescence quenching. Reverse-phase HPLC was performed on a Hitachi LaChrom Elite system with UV detection at 220 nm using a YMC-Pack ODS-A column (10 × 250 mm or 20 × 250 mm).

$^1$H NMR spectra were recorded on a Bruker AVANCE III HD400SJ (400 MHz) spectrometer. $^1$H NMR chemical shifts were referenced to tetramethylsilane (0 ppm). High-resolution mass spectra were measured on a Thermo Scientific Extractive Plus Orbitrap mass spectrometer by Dr. Keiko Kuwata (Nagoya University).

#### Reagent abbreviations

DIPEA, *N*,*N*-diisopropylethylamine; DMF, *N*,*N*-dimethylformamide; HBTU, *O*-(benzotriazole-1-yl)-*N*,*N*,*N*',*N*'-tetramethyluronium hexafluorophosphate; TFA, trifluoroacetic acid.

#### Synthesis of m$^D$cTMP and myrTMP

m$^D$cTMP and myrTMP were synthesized as described previously.[S2,S12]

#### Synthesis of Compound 3 (TMP-FL)

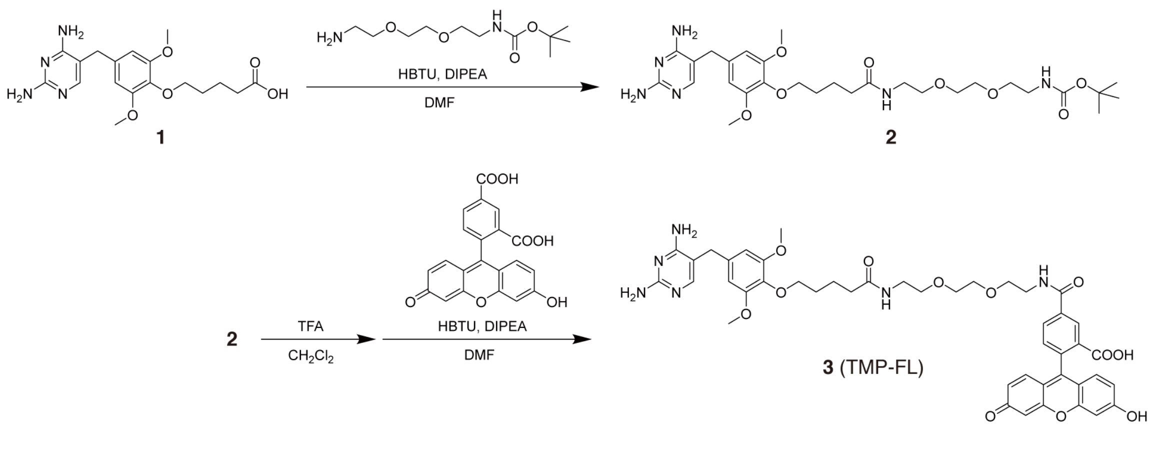


**Scheme S1.** Synthetic route of **3** (TMP-FL)

#### Synthesis of 2

Compound **1** was synthesized as described previously.[S13] To a stirred solution of **1** (102 mg, 0.27 mmol, 1.0 eq.), DIPEA (136 μL, 0.80 mmol, 3.0 eq.), and HBTU (114 mg, 0.30 mmol, 1.1 eq.) in DMF (1.5 mL) was added a solution of *N*-Boc-2,2'-(ethylenedioxy)diethylamine (84.2 mg, 0.34 mmol, 1.3 eq.) in DMF (0.5 mL). The mixture was stirred at room temperature for 2 h. The mixture was diluted with EtOAc (50 mL) and washed with saturated $NaHCO_3$ (30 mL × 2), water (30 mL × 2), and brine (30 mL). The organic layer was collected and dried over anhydrous $Na_2SO_4$. After filtration, the solvent was removed by evaporation to afford compound **2** as a pale-yellow solid (37 mg, 58 μmol, 23%),

which was used for the next reaction without further purification.
$^{1}$H NMR ($CDCl_3$): δ 7.76 (1H, s), 6.37 (2H, s), 4.96 (2H, s), 4.80 (2H, s), 3.94 (2H, t, *J* = 8.0 Hz), 3.78 (6H, s), 3.64 (2H, s), 3.56–3.59 (4H, m), 3.51–3.55 (4H, m), 3.29–3.32 (2H, m), 2.23 (2H, t, *J* = 8.0 Hz), 1.75–1.86 (4H, m), 1.44 (9H, s).

**Synthesis of 3 (TMP-FL)**

Compound **2** (10 mg, 17 μmol, 1.3 eq.) was dissolved in $CH_2Cl_2$ (2.0 mL) and TFA (1.0 mL). After stirring the mixture at room temperature for 1 h, the solvent was evaporated under reduced pressure. The residue was dissolved in DMF (200 μL). To this solution, 5-carboxyfluorescein (5.0 mg, 13 μmol, 1.0 eq.), DIPEA (23 μL, 132 μmol, 10 eq.), and HBTU (7.6 mg, 20 μmol, 1.5 eq.) were added. The mixture was stirred at room temperature for 6 h. After evaporating the solvent, the crude residue was purified by reverse-phase HPLC using a semi-preparative C18 column (a linear gradient of MeCN containing 0.1% TFA and 0.1% aqueous TFA) to afford **3** as a yellow solid (4.7 mg, 4.8 μmol, 36%).
$^{1}$H NMR ($CD_3OD$): δ 8.51 (1H, s), 8.22 (1H, dd, *J* = 1.6 and 8.0 Hz), 7.38 (1H, d, *J* = 8.0 Hz) 7.22 (1H, s), 6.65–6.85 (6H, m), 6.53 (1H, s), 3.87 (2H, t, *J* = 6.0 Hz), 3.37 (6H, s), 3.62–3.73 (8H, m), 3.56 (2H, t, *J* = 5.4 Hz), 3.35 (2H, t, *J* = 5.4 Hz), 2.25 (2H, 7.4 Hz), 1.73–1.81 (2H, m), 1.63–1.70 (2H, m).
HRMS (ESI): calculated for $[M+H]^+$, 865.3403; found, 865.3373.

### Supplementary Methods: Molecular and Cell Biology Experiments

#### Plasmid construction

All the cDNA and amino acid sequences of the constructs used in this study are listed in the "**Supplementary Sequences**" section. pPB-CAG.EBNXN (provided by Dr. Allan Bradley, Wellcome Trust Sanger Institute),[S14] pCSII-EF-MCS (provided by Dr. Hiroyuki Miyoshi, Keio University),[S15] and pCAGGS (provided by Dr. Jun-ichi Miyazaki, Osaka University)[S16] were used as vector backbones. pCSIIpuro[S17] was used as a lentiviral vector, and pPBbsr[S18] (blasticidin *S* resistance) and pPBpuro[S19] (puromycin resistance) were used as *piggyBac* donor vectors for the establishment of stable cell lines. For the protein expression in *E. coli* cells, pET-41a(+) (Novagen) was used as a vector backbone. All expression plasmids were generated using standard cloning procedures and the NEBuilder HiFi DNA assembly system (New England Biolabs). All PCR amplified sequences were verified by DNA sequencing. Complete plasmid sequences are available upon request.

#### Protein expression and purification

For *in vitro* experiments, wild-type eDHFR, K6-eDHFR, eDHFR$^{69K6}$, and eDHFR$^{145K6}$ were expressed as His-tag fusion proteins and purified as follows. The pET-41a(+) vector encoding the corresponding protein (**Figure S15**) was transformed into the *E. coli.* strain BL21(DE3), and the transformants were first cultured in 5 mL of LB broth containing 20 μg/mL kanamycin at 37 °C. The culture was transferred to 200 mL of Terrific Broth containing 20 μg/mL kanamycin and cells were grown at 37 °C until the $OD_{660}$ reached 0.6. Isopropyl-β-D-thiogalactoside was then added to a final concentration of 100 μM to induce protein expression. The cells expressing eDHFR, eDHFR$^{69K6}$, and K6-eDHFR were further cultured at 37 °C for 4 h. The cells expressing eDHFR$^{145K6}$ were further cultured at 16 °C for 24 h. The cells were collected by centrifugation, resuspended in buffer A (50 mM phosphate, 100 mM NaCl, 20 mM imidazole, pH 7.2), and disrupted by sonication on ice. The lysate was cleared by centrifugation and purified by a HisTrap FF column (GE Healthcare) according to the manufacture's protocol. The bound proteins were washed with buffer A and eluted with buffer B (50 mM phosphate, 100 mM NaCl, 250 mM imidazole, pH 7.2). The purified proteins were dialyzed and stored in 50 mM Tris-HCl, 100 mM NaCl, pH 7.4. The concentration of the proteins was determined by UV spectroscopy using the molar extinction coefficient at 280 nm of 33,710 $M^{-1}$ $cm^{-1}$.[S20]

#### Binding affinity assays (Figure S3)

The binding affinities of wild-type eDHFR, K6-eDHFR, eDHFR$^{69K6}$, and eDHFR$^{145K6}$ to TMP-FL (**Figure S3a**) were determined by fluorescence polarization assays using a Hitachi F-7000 fluorescence spectrophotometer equipped with polarizing filters. TMP-FL (10 nM) was titrated with varying concentrations of the purified eDHFR protein ranging from 1 nM to 1,000 nM in the presence of a 50-fold excess of NADPH in 50 mM Tris-HCl, 100 mM NaCl, pH 7.4. The fluorescence polarization value ($P$) was calculated by the following equation using the emission intensity at 526 nm (excitation at 488 nm):

$$P = \frac{I_{\mathrm{VV}} - \mathrm{G}I_{\mathrm{VH}}}{I_{\mathrm{VV}} + \mathrm{G}I_{\mathrm{VH}}}; \qquad \mathrm{G} = \frac{I_{\mathrm{HV}}}{I_{\mathrm{HH}}}$$

where $I_{\mathrm{VV}}$, $I_{\mathrm{VH}}$, $I_{\mathrm{HV}}$, and $I_{\mathrm{HH}}$ are the fluorescence intensity components, in which the subscripts refer to the vertical (V) or horizontal (H) position of the excitation and emission polarizers, respectively. G is a correction factor that detects the instrumental sensitivity of the polarization detection of the emission.

Fluorescence polarization values were plotted against the protein concentration, and the dissociation constants ($K_d$) were determined by fitting the titration data to the following equation based on a 1:1 binding model:

$$P = P_0 + (P_{max} - P_0)\frac{[TMP_{FL}] + [eDHFR] + K_d - \sqrt{([TMP_{FL}] + [eDHFR] + K_d)^2 - 4[TMP_{FL}][eDHFR]}}{2[TMP_{FL}]}$$

where $P$ is the polarization value of TMP-FL with varying concentrations of eDHFR, $P_0$ is the polarization value of TMP-FL in the absence of eDHFR, $P_{max}$ is the maximum polarization value with an infinite amount of eDHFR, $[TMP_{FL}]$ is the concentration of TMP-FL, and [eDHFR] is the concentration of eDHFR added.

**Cell culture and transfection**

HeLa, NIH3T3, and Cos-7 cells were obtained from the Cell Resource Center for Biomedical Research, Institute of Development, Aging and Cancer, Tohoku University. Lenti-X-293T cells were purchased from Clontech Laboratories. All cells were maintained in Dulbecco’s modified Eagle’s medium (DMEM) supplemented with 10% heat-inactivated fetal bovine serum (FBS), 100 U/mL penicillin, and 100 µg/mL streptomycin at 37 °C under a humidified 5% $CO_2$ atmosphere. For live-cell imaging, cells were seeded to 35 mm glass-bottomed dishes (Iwaki) coated with collagen type I-C (Nitta gelatin). For transient expression experiments, cells were transfected using 293fectin (Invitrogen) or Polyethylenimine “Max” (PEI-Max) (Polysciences Inc.) according to the manufacturer’s instructions.

**Establishment of stable cell lines**

A *piggyBac* transposon system[S14] was employed to establish cell lines stably expressing the constructs. Cells were cotransfected with *piggyBac* donor vector(s) (pPBpuro and/or pPBbsr) encoding the desired protein(s) and pCMV-mPBase encoding the *piggyBac* transposase (provided by Dr. Allan Bradley, Wellcome Trust Sanger Institute)[S14,S21] using PEI-Max or 293fectin. Cells were selected with 0.5–2 µg/mL puromycin and/or 5–10 µg/mL blasticidin S for at least 10 days. Bulk populations of selected cells were used.

For lentiviral production, Lenti-X-293T cells were cotransfected with the pCSIIpuro lentiviral vector encoding the desired protein, psPAX2 (Addgene plasmid #12260, provided by Dr. Didier Trono, Swiss Federal Institute of Technology in Lausanne), and pCMV-VSV-G-RSV-Rev (provided by Dr. Hiroyuki Miyoshi, Keio University)[S17] using PEI-Max. Virus-containing media were collected at 48 h after transfection, filtered, and used to infect target cells with 8 µg/mL polybrene. Cells were selected with 0.5 µg/mL puromycin for at least 7 days. Bulk populations of selected cells were used.

**Live-cell imaging**

Fluorescence imaging was performed on either (i) an IX83/FV3000 confocal laser-scanning microscope (Olympus) equipped with a PlanApo N 60×/1.42 NA oil objective lens (Olympus), a Z drift compensator system (IX3-ZDC2, Olympus), and a stage top incubator (Tokai Hit) or (ii) an IX83 (Olympus) equipped with a UPLXAPO60XO/1.42 NA oil objective lens (Olympus), a Z drift compensator system (IX3-ZDC2, Olympus), a sCMOS camera (Prime; Teledyne Photometrics), a spinning disk confocal unit (CSU-W1; Yokogawa Electric Corp.), and a stage top incubator (Tokai Hit). The microscope (ii) was controlled by MetaMorph software (Molecular Devices). Lasers used for excitation were as follows: 458 nm for mTFP1/mVenus FRET biosensor (ii), 488 nm for EGFP and mNeonGreen (i/ii), 561 nm for mCherry and mKO (i/ii), and 633 nm for miRFP670 (ii). The filters and dichroic mirrors used for the spinning disk confocal fluorescence imaging were as follows: two excitation dichroic mirrors (DM405/488/561/640 and DM 445/515/640) and four emission filters (482/35 for mTFP1, 525/50 for mNeonGreen and mVenus, 617/73 for mCherry, and 685/40 for miRFP670) (Yokogawa Electric Corp.). Unless otherwise noted, time-lapse live-cell imaging was performed at 37 °C under a 5% $CO_2$ atmosphere. Fluorescence images were analyzed using the Fiji distribution of ImageJ.[S22]

**SLIPT assays (Figure 1, Figure S1, and Figure S4)**

To conduct SLIPT assays of EGFP-tagged eDHFR variants, HeLa cells were plated at $1.0 \times 10^5$ cells in collagen-coated 35 mm glass-bottomed dishes and cultured for 24 h at 37 °C in 5% $CO_2$. The cells were transfected with pCMV-eDHFR-EGFP, pCMV-EGFP-K6-eDHFR, pCMV-EGFP-eDHFR(69K6), pCMV-eDHFR(69K6)-EGFP, pCMV-EGFP-eDHFR(145K6), or pCMV-eDHFR(145K6)-EGFP using PEI-Max. For the assay of K6-eDHFR-EGFP, HeLa cells stably expressing K6-eDHFR-EGFP (established in our previous work using pPBpuro-K6-eDHFR-EGFP)[S19] were used. Twenty-four hours after transfection, the medium was changed to serum-free DMEM supplemented with 100 U/mL penicillin and 100 μg/mL streptomycin [DMEM(–)], and the cells were observed by time-lapse imaging before and after addition of $m^{D}$cTMP (10 μM).

EGFP-$^{iK6}$DHFR translocation assays using NIH3T3 and Cos-7 cells were performed in the same manner as described above, except that cells were transfected with pCMV-EGFP-eDHFR(69K6) using 293fectin.

**Colocalization assays (Figure S2)**

For colocalization analysis, HeLa cells were plated at $1.0 \times 10^5$ cells in collagen-coated 35 mm glass-bottomed dishes and cultured for 24 h at 37 °C in 5% $CO_2$. The cells were cotransfected with pCMV-mCherry-Giantin and pCMV-eDHFR-EGFP, pCMV-EGFP-K6-eDHFR, pCMV-EGFP-eDHFR(69K6), pCMV-eDHFR(69K6)-EGFP, pCMV-EGFP-eDHFR(145K6), or pCMV-eDHFR(145K6)-EGFP at a 1:3 ratio using PEI-Max. HeLa cells stably expressing K6-eDHFR-EGFP were transfected with pCMV-mCherry-Giantin alone using PEI-Max. Twenty-four hours after transfection, the medium was changed to DMEM(–), and the cells were imaged before and 30 min after treatment with $m^{D}$cTMP (10 μM).

**Investigation of loop-K6-tag/PM interaction by lipid depletion (Figure S5)**

PS depletion experiments were performed as previously described.[S19] HeLa cells stably expressing EGFP-$^{iK6}$DHFR [established using pPBpuro-EGFP-eDHFR(69K6)] were plated at $1.5 \times 10^5$ cells in collagen-coated 35 mm glass-bottomed dishes and cultured for 24 h at 37 °C in 5% $CO_2$. The medium was changed to DMEM(–), and the cells were incubated with $m^{D}$cTMP (10 μM) at 37 °C for 30 min to recruit EGFP-$^{iK6}$DHFR to the PM. Then, the medium was changed to ionomycin buffer (20 mM

HEPES-KOH, 140 mM NaCl, 5 mM KCl, 1 mM $MgCl_2$, 0.7 mM $CaCl_2$, pH 7.4),[S23,S24] and the cells were imaged before and 10 min after treatment with ionomycin (10 μM) (Sigma).

PM PI4P and PI(4,5)$P_2$ depletion experiments were performed as previously described using a rapamycin-induced Pseudojanin recruitment system.[S1] HeLa cells stably expressing EGFP-eDHFR$^{iK6}$ were plated at 1.0 × $10^5$ cells in collagen-coated 35 mm glass-bottomed dishes and cultured for 24 h at 37 °C in 5% $CO_2$. The cells were cotransfected with plasmids encoding Pseudojanin[S1] (Addgene plasmids #20147, provided by Dr. Robin Irvine, University of Cambridge) [or PJ-DEAD[S1] (Addgene plasmid #38002, provided by Dr. Robin Irvine, University of Cambridge) for control] and Lyn11-targeted FRB[S7] (Addgene plasmid #20147, provided by Dr. Tobias Meyer, Stanford University) at a 1:1 ratio using 293fectin. Twenty-four hours after transfection, the medium was changed to DMEM(–), and the cells were incubated with $m^{D}$cTMP (10 μM) at 37 °C for 30 min to recruit EGFP-eDHFR$^{iK6}$ to the PM. Then, the cells were washed with DMEM(–) and imaged before and 10 min after treatment with rapamycin (200 nM).

**Investigation of the effect of $m^{D}$cTMP palmitoylation (Figure S6)**

HeLa cells stably expressing EGFP-$^{iK6}$DHFR were plated at 1.5 × $10^5$ cells in collagen-coated 35 mm glass-bottomed dishes and cultured for 24 h at 37 °C in 5% $CO_2$. The medium was changed to DMEM(–), and the cells were imaged before and 30 min after treatment with myrTMP (10 μM).

**Chemogenetic activation of the Raf/ERK pathway (Figure 2 and Figure S7)**

To evaluate chemogenetic Raf/ERK signal activation, HeLa cells stably expressing EGFP-$^{iK6}$DHFR-cRaf and ERK-KTR-mKO [established using pPBpuro-EGFP-eDHFR(69K6)-cRaf and pPBbsr-ERK-KTR-mKO] were plated at 1.5 × $10^5$ cells in collagen-coated 35 mm glass-bottomed dishes and cultured for 24 h at 37 °C in 5% $CO_2$. HeLa cells stably expressing EGFP-$^{iK6}$DHFR and ERK-KTR-mKO [established using pPBpuro-EGFP-eDHFR(69K6) and pPBbsr-ERK-KTR-mKO] were used as a control. After changing the medium to DMEM(–), the cells were serum-starved for 1 h. The cells were observed by time-lapse imaging before and after addition of $m^{D}$cTMP (10 μM).

For MEK inhibition experiments, cells expressing EGFP-$^{iK6}$DHFR-cRaf and ERK-KTR-mKO were preincubated with PD184352 (30 μM) (AdooQ Bioscience) in DMEM(–) for 4 h and subsequently observed by time-lapse imaging in the same manner as described above without washing.

**Chemogenetic activation of Gαq/$Ca^{2+}$ signaling (Figures 3 and Figure S8)**

To evaluate chemogenetic Gαq/$Ca^{2+}$ signal activation, HeLa cells were plated at 1.2 × $10^5$ cells in collagen-coated 35 mm glass-bottomed dishes and cultured for 24 h at 37 °C in 5% $CO_2$. The cells were cotransfected with pCAGGS-mNG-eDHFR(69K6)-Gαq and pCAGGS-R-GECO at a 1:2 ratio using 293fectin. Cells cotransfected with pCAGGS-mNG-eDHFR(69K6)-$Gαq_{L254A}$ and pCAGGS-R-GECO were used as a control. Twenty-four hours after transfection, the medium was changed to Hanks' Balanced Salt Solution (Gibco) containing 10 mM HEPES, and the cells were observed by time-lapse imaging before and after addition of $m^{D}$cTMP (5 μM).

**Chemogenetic activation of Gαs/cAMP signaling (Figure S9)**

To evaluate chemogenetic Gαs/cAMP signal activation, HeLa cells stably expressing miR-$^{iK6}$DHFR-Gαs and a cAMP sensor [established using pPBpuro-miR-eDHFR(69K6)-Gαs and pPBbsr-cAMPsensor] were plated at 2.5 × $10^5$ cells in 35 mm glass-bottomed dishes and cultured for 24 h at 37 °C in 5% $CO_2$. HeLa cells stably expressing miR-$^{iK6}$DHFR and a cAMP sensor [established using pCSIIpuro-miR-eDHFR(69K6) and pPBbsr-cAMPsensor] were used as a control. After changing the medium to DMEM(–), the cells were observed by time-lapse imaging before and after addition of m$^{D}$cTMP (10 μM).

**Chemogenetic activation of the Ras/ERK pathway** (**Figure S10**)

To evaluate chemogenetic Ras/ERK signal activation, HeLa cells stably expressing mNG-$^{iK6}$DHFR-RasGEF and mCh-ERK [established using pCSIIpuro-mNG-eDHFR(69K6)-RasGEF and pPBbsr-MEK-P2A-mCherry-ERK] were plated at 5 × $10^4$ cells in 35 mm glass-bottomed dishes and cultured for 24 h at 37 °C in 5% $CO_2$. HeLa cells stably expressing mNG-$^{iK6}$DHFR and mCh-ERK [established using pCSIIpuro-mNG-eDHFR(69K6) and pPBbsr-MEK-P2A-mCherry-ERK] were used as a control. After changing the medium to DMEM(–), the cells were observed by time-lapse imaging before and after addition of m$^{D}$cTMP (10 μM).

**Chemogenetic PI(3,4,5)$P_3$ production** (**Figure S11**)

To evaluate chemogenetic PI(3,4,5)$P_3$ production, HeLa cells stably expressing mNG-$^{iK6}$DHFR-$p85_{iSH2}$ and mCh-$PH_{Akt}$ [established using pPBpuro-mNG-eDHFR(69K6)-$p85_{iSH2}$ and pPBbsr-mCherry-$PH_{Akt}$] were plated at 5 × $10^4$ cells in 35 mm glass-bottomed dishes and cultured for 24 h at 37 °C in 5% $CO_2$. HeLa cells stably expressing mNG-$^{iK6}$DHFR and mCh-$PH_{Akt}$ [established using pCSIIpuro-mNG-eDHFR(69K6) and pPBbsr-mCherry-$PH_{Akt}$] were used as a control. After changing the medium to DMEM(–), the cells were observed by time-lapse imaging before and after addition of m$^{D}$cTMP (10 μM).

**Chemogenetic induction of lamellipodia formation** (**Figure S12**)

To evaluate chemogenetic Rac activation and lamellipodia formation, HeLa cells stably expressing mNG-$^{iK6}$DHFR-Tiam1 and Lifeact-mCh [established using pPBpuro-mNG-eDHFR(69K6)-Tiam1 and pPBbsr-Lifeact-mCherry] were plated at 5 × $10^4$ cells in 35 mm glass-bottomed dishes and cultured for 24 h at 37 °C in 5% $CO_2$. HeLa cells stably expressing mNG-$^{iK6}$DHFR and Lifeact-mCh [established using pCSIIpuro-mNG-eDHFR(69K6) and pPBbsr-Lifeact-mCherry] were used as a control. After changing the medium to DMEM(–), the cells were observed by time-lapse imaging before and after addition of m$^{D}$cTMP (5 μM).

**Chemical control of reversible protein translocation and signal oscillations** (**Figure 4** and **Figure S13**)

The culture-medium flow system was set up as shown in **Figure S13a**. In this system, a silicone attachment with a rhombic hole (Tokai Hit) was adsorbed on a collagen-coated 35 mm glass-bottomed dish (27Φ), and cells were plated in the rhombic area. Ligand solutions were added to the culture medium directly from above the dish, and the medium flow for cell washing was carried out using a peristaltic pump (ATTO) with a flow rate of 1 mL/min.

For the proof-of-principle experiment of repeatable SLIPT, HeLa cells stably expressing $^{iK6}$DHFR-EGFP [established using pPBpuro-eDHFR(69K6)-EGFP] were plated at 0.6 × $10^5$ cells in the rhombic area of collagen-coated 35 mm glass-bottomed dishes and cultured for 24 h at 37 °C in 5% $CO_2$. After changing the medium to DMEM(–), the dish was set on the microscope stage. Cells were observed by time-lapse imaging before and after inducing repeated protein translocation. First, PM recruitment of $^{iK6}$DHFR-EGFP was induced by incubating the cells with m$^{D}$cTMP (10 μM) for 20 min. Then, the cells were incubated with excess TMP (50 μM) for 20 min to return the PM-localized $^{iK6}$DHFR-EGFP to the cytoplasm. Next, the culture medium was replaced by continuously flowing fresh medium for 20 min, by which $^{iK6}$DHFR-EGFP was relocalized to the PM because of the removal of excess TMP. Subsequently, the cells were subjected to repeated cycles of TMP addition and medium exchange.

To demonstrate synthetic ERK signal oscillations, HeLa cells stably expressing cRaf -mNG-$^{iK6}$DHFR and mCh-ERK [established using pPBpuro-cRaf-mNG-eDHFR(69K6) and pPBbsr-mCherry-ERK] were plated at $0.6 \times 10^5$ cells in the rhombic area of collagen-coated 35 mm glass-bottomed dishes and cultured for 24 h at 37 °C in 5% $CO_2$. After changing the medium to DMEM(–), the cells were serum-starved for 1 h. The cells were observed by time-lapse imaging and subjected to repeated protein translocation as described above.

### Supplementary Sequences

#### pCMV-eDHFR-EGFP

```
>Amino acid sequence
MAISLIAALAVDRVIGMENAMPWNLPADLAWFKRNTLNKPVIMGRHTWESIGRPLPGRKNIILSSQPGTDDRV
TWVKSVDEAIAACGDVPEIMVIGGGRVYEQFLPKAQKLYLTHIDAEVEGDTHFPDYEPDDWESVFSEFHDADA
QNSHSYCFEILERRAAASDPPVATMVSKGEELFTGVVPILVELDGDVNGHKFSVSGEGEGDATYGKLTLKFIC
TTGKLPVPWPTLVTTLTYGVQCFSRYPDHMKQHDFFKSAMPEGYVQERTIFFKDDGNYKTRAEVKFEGDTLVN
RIELKGIDFKEDGNILGHKLEYNYNSHNVYIMADKQKNGIKVNFKIRHNIEDGSVQLADHYQQNTPIGDGPVL
LPDNHYLSTQSALSKDPNEKRDHMVLLEFVTAAGITLGMDELYK*

>DNA sequence
ATGGCTATCAGTCTGATTGCGGCGTTAGCGGTAGATCGCGTTATCGGCATGGAAAACGCCATGCCGTGGAACC
TGCCTGCCGATCTCGCCTGGTTTAAACGCAACACCTTAAATAAACCCGTGATTATGGGCCGCCATACCTGGGA
ATCAATCGGTCGTCCGTTGCCAGGACGCAAAAATATTATCCTCAGCAGTCAACCGGGTACGGACGATCGCGTA
ACGTGGGTGAAGTCGGTGGATGAAGCCATCGCGGCGTGTGGTGACGTACCAGAAATCATGGTGATTGGCGGCG
GTCGCGTTTATGAACAGTTCTTGCCAAAAGCGCAAAAACTGTATCTGACGCATATCGACGCAGAAGTGGAAGG
CGACACCCATTTCCCGGATTACGAGCCGGATGACTGGGAATCGGTATTCAGCGAATTCCACGATGCTGATGCG
CAGAACTCTCACAGCTATTGCTTTGAGATTCTGGAGCGGCGGGCGGCCGCTTCGGATCCACCGGTCGCCACCA
TGGTGAGCAAGGGCGAGGAGCTGTTCACCGGGGTGGTGCCCATCCTGGTCGAGCTGGACGGCGACGTAAACGG
CCACAAGTTCAGCGTGTCCGGCGAGGGCGAGGGCGATGCCACCTACGGCAAGCTGACCCTGAAGTTCATCTGC
ACCACCGGCAAGCTGCCCGTGCCCTGGCCCACCCTCGTGACCACCCTGACCTACGGCGTGCAGTGCTTCAGCC
GCTACCCCGACCACATGAAGCAGCACGACTTCTTCAAGTCCGCCATGCCCGAAGGCTACGTCCAGGAGCGCAC
CATCTTCTTCAAGGACGACGGCAACTACAAGACCCGCGCCGAGGTGAAGTTCGAGGGCGACACCCTGGTGAAC
CGCATCGAGCTGAAGGGCATCGACTTCAAGGAGGACGGCAACATCCTGGGGCACAAGCTGGAGTACAACTACA
ACAGCCACAACGTCTATATCATGGCCGACAAGCAGAAGAACGGCATCAAGGTGAACTTCAAGATCCGCCACAA
CATCGAGGACGGCAGCGTGCAGCTCGCCGACCACTACCAGCAGAACACCCCCATCGGCGACGGCCCCGTGCTG
CTGCCCGACAACCACTACCTGAGCACCCAGTCCGCCCTGAGCAAAGACCCCAACGAGAAGCGCGATCACATGG
TCCTGCTGGAGTTCGTGACCGCCGCCGGGATCACTCTCGGCATGGACGAGCTGTACAAGTAA

eDHFR EGFP
```

#### pPBpuro-K6-eDHFR-EGFP

>Amino acid sequence
MKKKKKKGSGASAGGGSGAGSGAISLIAALAVDRVIGMENAMPWNLPADLAWFKRNTLNKPVIMGRHTWESIG
RPLPGRKNIILSSQPGTDDRVTWVKSVDEAIAACGDVPEIMVIGGGRVYEQFLPKAQKLYLTHIDAEVEGDTH
FPDYEPDDWESVFSEFHDADAQNSHSYCFEILERRAAASDPPVATMVSKGEELFTGVVPILVELDGDVNGHKF
SVSGEGEGDATYGKLTLKFICTTGKLPVPWPTLVTTLTYGVQCFSRYPDHMKQHDFFKSAMPEGYVQERTIFF
KDDGNYKTRAEVKFEGDTLVNRIELKGIDFKEDGNILGHKLEYNYNSHNVYIMADKQKNGIKVNFKIRHNIED
GSVQLADHYQQNTPIGDGPVLLPDNHYLSTQSALSKDPNEKRDHMVLLEFVTAAGITLGMDELYKSGLRSRQG
SGAGSGAGSGAGSGAGSGAPRAQASNSAVDGTAGPG*-[*IRES*]-MTEYKPTVRLATRDDVPRAVRTLAAAFA
DYPATRHTVDPDRHIERVTELQELFLTRVGLDIGKVWVADDGAAVAVWTTPESVEAGAVFAEIGPRMAELSGS
RLAAQQQMEGLLAPHRPKEPAWFLATVGVSPDHQGKGLGSAVVLPGVEAAERAGVPAFLETSAPRNLPFYERL
GFTVTADVECPKDRATWCMTRKPGA*

>DNA sequence
ATGAAAAAAAAGAAAAAGAAAGGCTCCGGTGCCAGTGCTGGTGGTGGCAGCGGTGCTGGTTCCGGCGCTATCA
GTCTGATTGCGGCGTTAGCGGTAGATCGCGTTATCGGCATGGAAAACGCCATGCCGTGGAACCTGCCTGCCGA
TCTCGCCTGGTTTAAACGCAACACCTTAAATAAACCCGTGATTATGGGCCGCCATACCTGGGAATCAATCGGT
CGTCCGTTGCCAGGACGCAAAAATATTATCCTCAGCAGTCAACCGGGTACGGACGATCGCGTAACGTGGGTGA
AGTCGGTGGATGAAGCCATCGCGGCGTGTGGTGACGTACCAGAAATCATGGTGATTGGCGGCGGTCGCGTTTA
TGAACAGTTCTTGCCAAAAGCGCAAAAACTGTATCTGACGCATATCGACGCAGAAGTGGAAGGCGACACCCAT
TTCCCGGATTACGAGCCGGATGACTGGGAATCGGTATTCAGCGAATTCCACGATGCTGATGCGCAGAACTCTC
ACAGCTATTGCTTTGAGATTCTGGAGCGGCGGGCGGCCGCTTCGGATCCACCGGTCGCCACCATGGTGAGCAA
GGGCGAGGAGCTGTTCACCGGGGTGGTGCCCATCCTGGTCGAGCTGGACGGCGACGTAAACGGCCACAAGTTC
AGCGTGTCCGGCGAGGGCGAGGGCGATGCCACCTACGGCAAGCTGACCCTGAAGTTCATCTGCACCACCGGCA
AGCTGCCCGTGCCCTGGCCCACCCTCGTGACCACCCTGACCTACGGCGTGCAGTGCTTCAGCCGCTACCCCGA
CCACATGAAGCAGCACGACTTCTTCAAGTCCGCCATGCCCGAAGGCTACGTCCAGGAGCGCACCATCTTCTTC
AAGGACGACGGCAACTACAAGACCCGCGCCGAGGTGAAGTTCGAGGGCGACACCCTGGTGAACCGCATCGAGC
TGAAGGGCATCGACTTCAAGGAGGACGGCAACATCCTGGGGCACAAGCTGGAGTACAACTACAACAGCCACAA
CGTCTATATCATGGCCGACAAGCAGAAGAACGGCATCAAGGTGAACTTCAAGATCCGCCACAACATCGAGGAC
GGCAGCGTGCAGCTCGCCGACCACTACCAGCAGAACACCCCCATCGGCGACGGCCCCGTGCTGCTGCCCGACA
ACCACTACCTGAGCACCCAGTCCGCCCTGAGCAAAGACCCCAACGAGAAGCGCGATCACATGGTCCTGCTGGA
GTTCGTGACCGCCGCCGGGATCACTCTCGGCATGGACGAGCTGTACAAGTCCGGACTCAGATCTCGACAAGGT
AGTGGTGCTGGCTCTGGTGCTGGTAGTGGCGCTGGTTCCGGTGCTGGCTCTGGCGCGCCTCGAGCTCAAGCTT
CGAATTCTGCAGTCGACGGTACCGCGGGCCCGGGATAAGTCAACTAACTTAAGCTAGCAACGGTTTCCCTCTA
GCGGGATCAATTCCGCCCCCCCCCCCTAACGTTACTGGCCGAAGCCGCTTGGAATAAGGCCGGTGTGCGTTTG
TCTATATGTTATTTTCCACCATATTGCCGTCTTTTGGCAATGTGAGGGCCCGGAAACCTGGCCCTGTCTTCTT
GACGAGCATTCCTAGGGGTCTTTCCCCTCTCGCCAAAGGAATGCAAGGTCTGTTGAATGTCGTGAAGGAAGCA
GTTCCTCTGGAAGCTTCTTGAAGACAAACAACGTCTGTAGCGACCCTTTGCAGGCAGCGGAACCCCCCACCTG
GCGACAGGTGCCTCTGCGGCCAAAAGCCACGTGTATAAGATACACCTGCAAAGGCGGCACAACCCCAGTGCCA
CGTTGTGAGTTGGATAGTTGTGGAAAGAGTCAAATGGCTCTCCTCAAGCGTATTCAACAAGGGGCTGAAGGAT
GCCCAGAAGGTACCCCATTGTATGGGATCTGATCTGGGGCCTCGGTGCACATGCTTTACATGTGTTTAGTCGA
GGTTAAAAAACGTCTAGGCCCCCCGAACCACGGGGACGTGGTTTTCCTTTGAAAAACACGATAATACCATGAC
CGAGTACAAGCCCACGGTGCGCCTCGCCACCCGCGACGACGTCCCCAGGGCCGTACGCACCCTCGCCGCCGCG
TTCGCCGACTACCCCGCCACGCGCCACACCGTCGATCCGGACCGCCACATCGAGCGGGTCACCGAGCTGCAAG
AACTCTTCCTCACGCGCGTCGGGCTCGACATCGGCAAGGTGTGGGTCGCGGACGACGGCGCCGCGGTGGCGGT
CTGGACCACGCCGGAGAGCGTCGAAGCGGGGGCGGTGTTCGCCGAGATCGGCCCGCGCATGGCCGAGTTGAGC
GGTTCCCGGCTGGCCGCGCAGCAACAGATGGAAGGCCTCCTGGCGCCGCACCGGCCCAAGGAGCCCGCGTGGT
TCCTGGCCACCGTCGGCGTCTCGCCCGACCACCAGGGCAAGGGTCTGGGCAGCGCCGTCGTGCTCCCCGGAGT
GGAGGCGGCCGAGCGCGCCGGGGTGCCCGCCTTCCTGGAGACCTCCGCGCCCCGCAACCTCCCCTTCTACGAG
CGGCTCGGCTTCACCGTCACCGCCGACGTCGAGTGCCCGAAGGACCGCGCGACCTGGTGCATGACCCGCAAGC
CCGGTGCCTGA

K6-tag eDHFR EGFP *IRES* Puro$^{R}$ (puromycin *N*-acetyltransferase)

**pCMV-EGFP-K6-eDHFR**

>Amino acid sequence
MVSKGEELFTGVVPILVELDGDVNGHKFSVSGEGEGDATYGKLTLKFICTTGKLPVPWPTLVTTLTYGVQCFS
RYPDHMKQHDFFKSAMPEGYVQERTIFFKDDGNYKTRAEVKFEGDTLVNRIELKGIDFKEDGNILGHKLEYNY
NSHNVYIMADKQKNGIKVNFKIRHNIEDGSVQLADHYQQNTPIGDGPVLLPDNHYLSTQSALSKDPNEKRDHM
VLLEFVTAAGITLGMDELYKASAGSGAGSGKKKKKKGSGASAGGGSGAGSGAISLIAALAVDRVIGMENAMPW
NLPADLAWFKRNTLNKPVIMGRHTWESIGRPLPGRKNIILSSQPGTDDRVTWVKSVDEAIAACGDVPEIMVIG
GGRVYEQFLPKAQKLYLTHIDAEVEGDTHFPDYEPDDWESVFSEFHDADAQNSHSYCFEILERR*

>DNA sequence
ATGGTGAGCAAGGGCGAGGAGCTGTTCACCGGGGTGGTGCCCATCCTGGTCGAGCTGGACGGCGACGTAAACG
GCCACAAGTTCAGCGTGTCCGGCGAGGGCGAGGGCGATGCCACCTACGGCAAGCTGACCCTGAAGTTCATCTG
CACCACCGGCAAGCTGCCCGTGCCCTGGCCCACCCTCGTGACCACCCTGACCTACGGCGTGCAGTGCTTCAGC
CGCTACCCCGACCACATGAAGCAGCACGACTTCTTCAAGTCCGCCATGCCCGAAGGCTACGTCCAGGAGCGCA
CCATCTTCTTCAAGGACGACGGCAACTACAAGACCCGCGCCGAGGTGAAGTTCGAGGGCGACACCCTGGTGAA
CCGCATCGAGCTGAAGGGCATCGACTTCAAGGAGGACGGCAACATCCTGGGGCACAAGCTGGAGTACAACTAC
AACAGCCACAACGTCTATATCATGGCCGACAAGCAGAAGAACGGCATCAAGGTGAACTTCAAGATCCGCCACA
ACATCGAGGACGGCAGCGTGCAGCTCGCCGACCACTACCAGCAGAACACCCCCATCGGCGACGGCCCCGTGCT
GCTGCCCGACAACCACTACCTGAGCACCCAGTCCGCCCTGAGCAAAGACCCCAACGAGAAGCGCGATCACATG
GTCCTGCTGGAGTTCGTGACCGCCGCCGGGATCACTCTCGGCATGGACGAGCTGTACAAGGCTTCTGCGGGCT
CTGGTGCTGGTAGTGGCAAAAAAAAGAAAAAGAAAGGCTCCGGTGCCAGTGCTGGTGGTGGCAGCGGTGCTGG
TTCCGGCGCTATCAGTCTGATTGCGGCGTTAGCGGTAGATCGCGTTATCGGCATGGAAAACGCCATGCCGTGG
AACCTGCCTGCCGATCTCGCCTGGTTTAAACGCAACACCTTAAATAAACCCGTGATTATGGGCCGCCATACCT
GGGAATCAATCGGTCGTCCGTTGCCAGGACGCAAAAATATTATCCTCAGCAGTCAACCGGGTACGGACGATCG
CGTAACGTGGGTGAAGTCGGTGGATGAAGCCATCGCGGCGTGTGGTGACGTACCAGAAATCATGGTGATTGGC
GGCGGTCGCGTTTATGAACAGTTCTTGCCAAAAGCGCAAAAACTGTATCTGACGCATATCGACGCAGAAGTGG
AAGGCGACACCCATTTCCCGGATTACGAGCCGGATGACTGGGAATCGGTATTCAGCGAATTCCACGATGCTGA
TGCGCAGAACTCTCACAGCTATTGCTTTGAGATTCTGGAGCGGCGGTAA

EGFP K6-tag eDHFR

**pCMV-eDHFR(69K6)-EGFP**

>Amino acid sequence
MISLIAALAVDRVIGMENAMPWNLPADLAWFKRNTLNKPVIMGRHTWESIGRPLPGRKNIILSSQPGTDKKKK
KKDRVTWVKSVDEAIAACGDVPEIMVIGGGRVYEQFLPKAQKLYLTHIDAEVEGDTHFPDYEPDDWESVFSEF
HDADAQNSHSYCFEILERRSGSGDPPVATMVSKGEELFTGVVPILVELDGDVNGHKFSVSGEGEGDATYGKLT
LKFICTTGKLPVPWPTLVTTLTYGVQCFSRYPDHMKQHDFFKSAMPEGYVQERTIFFKDDGNYKTRAEVKFEG
DTLVNRIELKGIDFKEDGNILGHKLEYNYNSHNVYIMADKQKNGIKVNFKIRHNIEDGSVQLADHYQQNTPIG
DGPVLLPDNHYLSTQSALSKDPNEKRDHMVLLEFVTAAGITLGMDELYK*

>DNA sequence
ATGATCAGTCTGATTGCGGCGTTAGCGGTAGATCGCGTTATCGGCATGGAAAACGCCATGCCGTGGAACCTGC
CTGCCGATCTCGCCTGGTTTAAACGCAACACCTTAAATAAACCCGTGATTATGGGCCGCCATACCTGGGAATC
AATCGGTCGTCCGTTGCCAGGACGCAAAAATATTATCCTCAGCAGTCAACCGGGTACGGACAAAAAAAAGAAA
AAGAAAGATCGCGTAACGTGGGTGAAGTCGGTGGATGAAGCCATCGCGGCGTGTGGTGACGTACCAGAAATCA
TGGTGATTGGCGGCGGTCGCGTTTATGAACAGTTCTTGCCAAAAGCGCAAAAACTGTATCTGACGCATATCGA
CGCAGAAGTGGAAGGCGACACCCATTTCCCGGATTACGAGCCGGATGACTGGGAATCGGTATTCAGCGAATTC
CACGATGCTGATGCGCAGAACTCTCACAGCTATTGCTTTGAGATTCTGGAGCGGCGGAGCGGCTCCGGGGATC
CACCGGTCGCCACCATGGTGAGCAAGGGCGAGGAGCTGTTCACCGGGGTGGTGCCCATCCTGGTCGAGCTGGA
CGGCGACGTAAACGGCCACAAGTTCAGCGTGTCCGGCGAGGGCGAGGGCGATGCCACCTACGGCAAGCTGACC
CTGAAGTTCATCTGCACCACCGGCAAGCTGCCCGTGCCCTGGCCCACCCTCGTGACCACCCTGACCTACGGCG
TGCAGTGCTTCAGCCGCTACCCCGACCACATGAAGCAGCACGACTTCTTCAAGTCCGCCATGCCCGAAGGCTA
CGTCCAGGAGCGCACCATCTTCTTCAAGGACGACGGCAACTACAAGACCCGCGCCGAGGTGAAGTTCGAGGGC
GACACCCTGGTGAACCGCATCGAGCTGAAGGGCATCGACTTCAAGGAGGACGGCAACATCCTGGGGCACAAGC
TGGAGTACAACTACAACAGCCACAACGTCTATATCATGGCCGACAAGCAGAAGAACGGCATCAAGGTGAACTT
CAAGATCCGCCACAACATCGAGGACGGCAGCGTGCAGCTCGCCGACCACTACCAGCAGAACACCCCCATCGGC
GACGGCCCCGTGCTGCTGCCCGACAACCACTACCTGAGCACCCAGTCCGCCCTGAGCAAAGACCCCAACGAGA
AGCGCGATCACATGGTCCTGCTGGAGTTCGTGACCGCCGCCGGGATCACTCTCGGCATGGACGAGCTGTACAA
GTAA

eDHFR K6-tag(69K6) EGFP

**pCMV-EGFP-eDHFR(69K6)**

```
>Amino acid sequence
MVSKGEELFTGVVPILVELDGDVNGHKFSVSGEGEGDATYGKLTLKFICTTGKLPVPWPTLVTTLTYGVQCFS
RYPDHMKQHDFFKSAMPEGYVQERTIFFKDDGNYKTRAEVKFEGDTLVNRIELKGIDFKEDGNILGHKLEYNY
NSHNVYIMADKQKNGIKVNFKIRHNIEDGSVQLADHYQQNTPIGDGPVLLPDNHYLSTQSALSKDPNEKRDHM
VLLEFVTAAGITLGMDELYKSGSAASSAGSMISLIAALAVDRVIGMENAMPWNLPADLAWFKRNTLNKPVIMG
RHTWESIGRPLPGRKNIILSSQPGTDKKKKKKDRVTWVKSVDEAIAACGDVPEIMVIGGGRVYEQFLPKAQKL
YLTHIDAEVEGDTHFPDYEPDDWESVFSEFHDADAQNSHSYCFEILERR*

>DNA sequence
ATGGTGAGCAAGGGCGAGGAGCTGTTCACCGGGGTGGTGCCCATCCTGGTCGAGCTGGACGGCGACGTAAACG
GCCACAAGTTCAGCGTGTCCGGCGAGGGCGAGGGCGATGCCACCTACGGCAAGCTGACCCTGAAGTTCATCTG
CACCACCGGCAAGCTGCCCGTGCCCTGGCCCACCCTCGTGACCACCCTGACCTACGGCGTGCAGTGCTTCAGC
CGCTACCCCGACCACATGAAGCAGCACGACTTCTTCAAGTCCGCCATGCCCGAAGGCTACGTCCAGGAGCGCA
CCATCTTCTTCAAGGACGACGGCAACTACAAGACCCGCGCCGAGGTGAAGTTCGAGGGCGACACCCTGGTGAA
CCGCATCGAGCTGAAGGGCATCGACTTCAAGGAGGACGGCAACATCCTGGGGCACAAGCTGGAGTACAACTAC
AACAGCCACAACGTCTATATCATGGCCGACAAGCAGAAGAACGGCATCAAGGTGAACTTCAAGATCCGCCACA
ACATCGAGGACGGCAGCGTGCAGCTCGCCGACCACTACCAGCAGAACACCCCCATCGGCGACGGCCCCGTGCT
GCTGCCCGACAACCACTACCTGAGCACCCAGTCCGCCCTGAGCAAAGACCCCAACGAGAAGCGCGATCACATG
GTCCTGCTGGAGTTCGTGACCGCCGCCGGGATCACTCTCGGCATGGACGAGCTGTACAAGTCTGGGTCCGCAG
CCAGTAGCGCCGGAAGTATGATCAGTCTGATTGCGGCGTTAGCGGTAGATCGCGTTATCGGCATGGAAAACGC
CATGCCGTGGAACCTGCCTGCCGATCTCGCCTGGTTTAAACGCAACACCTTAAATAAACCCGTGATTATGGGC
CGCCATACCTGGGAATCAATCGGTCGTCCGTTGCCAGGACGCAAAAATATTATCCTCAGCAGTCAACCGGGTA
CGGACAAAAAAAAGAAAAAGAAAGATCGCGTAACGTGGGTGAAGTCGGTGGATGAAGCCATCGCGGCGTGTGG
TGACGTACCAGAAATCATGGTGATTGGCGGCGGTCGCGTTTATGAACAGTTCTTGCCAAAAGCGCAAAAACTG
TATCTGACGCATATCGACGCAGAAGTGGAAGGCGACACCCATTTCCCGGATTACGAGCCGGATGACTGGGAAT
CGGTATTCAGCGAATTCCACGATGCTGATGCGCAGAACTCTCACAGCTATTGCTTTGAGATTCTGGAGCGGCG
GTAA
```

EGFP eDHFR K6-tag(69K6)

**pCMV-eDHFR(145K6)-EGFP**

>Amino acid sequence
MISLIAALAVDRVIGMENAMPWNLPADLAWFKRNTLNKPVIMGRHTWESIGRPLPGRKNIILSSQPGTDDRVT
WVKSVDEAIAACGDVPEIMVIGGGRVYEQFLPKAQKLYLTHIDAEVEGDTHFPDYEPDDWESVFSEFHDADAK
KKKKKQNSHSYCFEILERRSGSGDPPVATMVSKGEELFTGVVPILVELDGDVNGHKFSVSGEGEGDATYGKLT
LKFICTTGKLPVPWPTLVTTLTYGVQCFSRYPDHMKQHDFFKSAMPEGYVQERTIFFKDDGNYKTRAEVKFEG
DTLVNRIELKGIDFKEDGNILGHKLEYNYNSHNVYIMADKQKNGIKVNFKIRHNIEDGSVQLADHYQQNTPIG
DGPVLLPDNHYLSTQSALSKDPNEKRDHMVLLEFVTAAGITLGMDELYK*

>DNA sequence
ATGATCAGTCTGATTGCGGCGTTAGCGGTAGATCGCGTTATCGGCATGGAAAACGCCATGCCGTGGAACCTGC
CTGCCGATCTCGCCTGGTTTAAACGCAACACCTTAAATAAACCCGTGATTATGGGCCGCCATACCTGGGAATC
AATCGGTCGTCCGTTGCCAGGACGCAAAAATATTATCCTCAGCAGTCAACCGGGTACGGACGATCGCGTAACG
TGGGTGAAGTCGGTGGATGAAGCCATCGCGGCGTGTGGTGACGTACCAGAAATCATGGTGATTGGCGGCGGTC
GCGTTTATGAACAGTTCTTGCCAAAAGCGCAAAAACTGTATCTGACGCATATCGACGCAGAAGTGGAAGGCGA
CACCCATTTCCCGGATTACGAGCCGGATGACTGGGAATCGGTATTCAGCGAATTCCACGATGCTGATGCGAAA
AAAAAGAAAAAGAAACAGAACTCTCACAGCTATTGCTTTGAGATTCTGGAGCGGCGGAGCGGCTCCGGGGATC
CACCGGTCGCCACCATGGTGAGCAAGGGCGAGGAGCTGTTCACCGGGGTGGTGCCCATCCTGGTCGAGCTGGA
CGGCGACGTAAACGGCCACAAGTTCAGCGTGTCCGGCGAGGGCGAGGGCGATGCCACCTACGGCAAGCTGACC
CTGAAGTTCATCTGCACCACCGGCAAGCTGCCCGTGCCCTGGCCCACCCTCGTGACCACCCTGACCTACGGCG
TGCAGTGCTTCAGCCGCTACCCCGACCACATGAAGCAGCACGACTTCTTCAAGTCCGCCATGCCCGAAGGCTA
CGTCCAGGAGCGCACCATCTTCTTCAAGGACGACGGCAACTACAAGACCCGCGCCGAGGTGAAGTTCGAGGGC
GACACCCTGGTGAACCGCATCGAGCTGAAGGGCATCGACTTCAAGGAGGACGGCAACATCCTGGGGCACAAGC
TGGAGTACAACTACAACAGCCACAACGTCTATATCATGGCCGACAAGCAGAAGAACGGCATCAAGGTGAACTT
CAAGATCCGCCACAACATCGAGGACGGCAGCGTGCAGCTCGCCGACCACTACCAGCAGAACACCCCCATCGGC
GACGGCCCCGTGCTGCTGCCCGACAACCACTACCTGAGCACCCAGTCCGCCCTGAGCAAAGACCCCAACGAGA
AGCGCGATCACATGGTCCTGCTGGAGTTCGTGACCGCCGCCGGGATCACTCTCGGCATGGACGAGCTGTACAA
GTAA

eDHFR K6-tag(145K6) EGFP

**pCMV-EGFP-eDHFR(145K6)**

>Amino acid sequence
MVSKGEELFTGVVPILVELDGDVNGHKFSVSGEGEGDATYGKLTLKFICTTGKLPVPWPTLVTTLTYGVQCFS
RYPDHMKQHDFFKSAMPEGYVQERTIFFKDDGNYKTRAEVKFEGDTLVNRIELKGIDFKEDGNILGHKLEYNY
NSHNVYIMADKQKNGIKVNFKIRHNIEDGSVQLADHYQQNTPIGDGPVLLPDNHYLSTQSALSKDPNEKRDHM
VLLEFVTAAGITLGMDELYKSGSAASSAGSMISLIAALAVDRVIGMENAMPWNLPADLAWFKRNTLNKPVIMG
RHTWESIGRPLPGRKNIILSSQPGTDDRVTWVKSVDEAIAACGDVPEIMVIGGGRVYEQFLPKAQKLYLTHID
AEVEGDTHFPDYEPDDWESVFSEFHDADAKKKKKKQNSHSYCFEILERR*

>DNA sequence
ATGGTGAGCAAGGGCGAGGAGCTGTTCACCGGGGTGGTGCCCATCCTGGTCGAGCTGGACGGCGACGTAAACG
GCCACAAGTTCAGCGTGTCCGGCGAGGGCGAGGGCGATGCCACCTACGGCAAGCTGACCCTGAAGTTCATCTG
CACCACCGGCAAGCTGCCCGTGCCCTGGCCCACCCTCGTGACCACCCTGACCTACGGCGTGCAGTGCTTCAGC
CGCTACCCCGACCACATGAAGCAGCACGACTTCTTCAAGTCCGCCATGCCCGAAGGCTACGTCCAGGAGCGCA
CCATCTTCTTCAAGGACGACGGCAACTACAAGACCCGCGCCGAGGTGAAGTTCGAGGGCGACACCCTGGTGAA
CCGCATCGAGCTGAAGGGCATCGACTTCAAGGAGGACGGCAACATCCTGGGGCACAAGCTGGAGTACAACTAC
AACAGCCACAACGTCTATATCATGGCCGACAAGCAGAAGAACGGCATCAAGGTGAACTTCAAGATCCGCCACA
ACATCGAGGACGGCAGCGTGCAGCTCGCCGACCACTACCAGCAGAACACCCCCATCGGCGACGGCCCCGTGCT
GCTGCCCGACAACCACTACCTGAGCACCCAGTCCGCCCTGAGCAAAGACCCCAACGAGAAGCGCGATCACATG
GTCCTGCTGGAGTTCGTGACCGCCGCCGGGATCACTCTCGGCATGGACGAGCTGTACAAGTCTGGGTCCGCAG
CCAGTAGCGCCGGAAGTATGATCAGTCTGATTGCGGCGTTAGCGGTAGATCGCGTTATCGGCATGGAAAACGC
CATGCCGTGGAACCTGCCTGCCGATCTCGCCTGGTTTAAACGCAACACCTTAAATAAACCCGTGATTATGGGC
CGCCATACCTGGGAATCAATCGGTCGTCCGTTGCCAGGACGCAAAAATATTATCCTCAGCAGTCAACCGGGTA
CGGACGATCGCGTAACGTGGGTGAAGTCGGTGGATGAAGCCATCGCGGCGTGTGGTGACGTACCAGAAATCAT
GGTGATTGGCGGCGGTCGCGTTTATGAACAGTTCTTGCCAAAAGCGCAAAAACTGTATCTGACGCATATCGAC
GCAGAAGTGGAAGGCGACACCCATTTCCCGGATTACGAGCCGGATGACTGGGAATCGGTATTCAGCGAATTCC
ACGATGCTGATGCGAAAAAAAAGAAAAAGAAACAGAACTCTCACAGCTATTGCTTTGAGATTCTGGAGCGGCG
GTAA

EGFP eDHFR K6-tag(145K6)

**pCMV-mCherry-Giantin**

>Amino acid sequence
MVSKGEEDNMAIIKEFMRFKVHMEGSVNGHEFEIEGEGEGRPYEGTQTAKLKVTKGGPLPFAWDILSPQFMYG
SKAYVKHPADIPDYLKLSFPEGFKWERVMNFEDGGVVTVTQDSSLQDGEFIYKVKLRGTNFPSDGPVMQKKTM
GWEASSERMYPEDGALKGEIKQRLKLKDGGHYDAEVKTTYKAKKPVQLPGAYNVNIKLDITSHNEDYTIVEQY
ERAEGRHSTGGMDELYKSGLRSRGEPQQSFSEAQQQLCNTRQEVNELRKLLEEERDQRVAAENALSVAEEQIR
RLEHSEWDSSRTPIIGSCGTQEQALLIDLTSNSCRRTRSGVGWKRVLRSLCHSRTRVPLLAAIYFLMIHVLLI
LCFTGHL*

>DNA sequence
ATGGTGAGCAAGGGCGAGGAGGATAACATGGCCATCATCAAGGAGTTCATGCGCTTCAAGGTGCACATGGAGG
GCTCCGTGAACGGCCACGAGTTCGAGATCGAGGGCGAGGGCGAGGGCCGCCCCTACGAGGGCACCCAGACCGC
CAAGCTGAAGGTGACCAAGGGTGGCCCCCTGCCCTTCGCCTGGGACATCCTGTCCCCTCAGTTCATGTACGGC
TCCAAGGCCTACGTGAAGCACCCCGCCGACATCCCCGACTACTTGAAGCTGTCCTTCCCCGAGGGCTTCAAGT
GGGAGCGCGTGATGAACTTCGAGGACGGCGGCGTGGTGACCGTGACCCAGGACTCCTCCCTGCAGGACGGCGA
GTTCATCTACAAGGTGAAGCTGCGCGGCACCAACTTCCCCTCCGACGGCCCCGTAATGCAGAAGAAGACCATG
GGCTGGGAGGCCTCCTCCGAGCGGATGTACCCCGAGGACGGCGCCCTGAAGGGCGAGATCAAGCAGAGGCTGA
AGCTGAAGGACGGCGGCCACTACGACGCTGAGGTCAAGACCACCTACAAGGCCAAGAAGCCCGTGCAGCTGCC
CGGCGCCTACAACGTCAACATCAAGTTGGACATCACCTCCCACAACGAGGACTACACCATCGTGGAACAGTAC
GAACGCGCCGAGGGCCGCCACTCCACCGGCGGCATGGACGAGCTGTACAAGTCCGGACTCAGATCTCGAGGAG
AACCGCAGCAAAGCTTTTCTGAAGCTCAGCAGCAGCTATGCAACACCAGACAGGAAGTGAATGAATTAAGGAA
GCTGCTGGAAGAAGAACGAGACCAAAGAGTGGCTGCTGAGAATGCTCTCTCTGTGGCCGAGGAGCAGATCAGA
CGGTTAGAGCACAGTGAATGGGACTCTTCCCGGACTCCTATCATTGGCTCCTGTGGCACTCAGGAGCAGGCAC
TGTTAATAGATCTTACAAGCAACAGTTGTCGAAGGACCCGGAGTGGCGTTGGATGGAAGCGAGTCCTGCGTTC
ACTCTGTCATTCACGGACCCGAGTGCCACTTCTAGCAGCCATCTACTTTCTAATGATTCATGTCCTGCTCATT
CTGTGTTTTACGGGCCATCTATAG

mCherry the Golgi-targeting motif from human giantin (residues 3131–3259)[S25]

**pPBpuro-EGFP-eDHFR(69K6)-cRaf**

>Amino acid sequence
MVSKGEELFTGVVPILVELDGDVNGHKFSVSGEGEGDATYGKLTLKFICTTGKLPVPWPTLVTTLTYGVQCFS
RYPDHMKQHDFFKSAMPEGYVQERTIFFKDDGNYKTRAEVKFEGDTLVNRIELKGIDFKEDGNILGHKLEYNY
NSHNVYIMADKQKNGIKVNFKIRHNIEDGSVQLADHYQQNTPIGDGPVLLPDNHYLSTQSALSKDPNEKRDHM
VLLEFVTAAGITLGMDELYKSGSAASSAGSMISLIAALAVDRVIGMENAMPWNLPADLAWFKRNTLNKPVIMG
RHTWESIGRPLPGRKNIILSSQPGTDKKKKKKDRVTWVKSVDEAIAACGDVPEIMVIGGGRVYEQFLPKAQKL
YLTHIDAEVEGDTHFPDYEPDDWESVFSEFHDADAQNSHSYCFEILERRSAGGSAGGSAGGSAGGSAGGPRSL
EMEHIQGAWKTISNGFGFKDAVFDGSSCISPTIVQQFGYQRRASDDGKLTDPSKTSNTIRVFLPNKQRTVVNV
RNGMSLHDCLMKALKVRGLQPECCAVFRLLHEHKGKKARLDWNTDAASLIGEELQVDFLDHVPLTTHNFARKT
FLKLAFCDICQKFLLNGFRCQTCGYKFHEHCSTKVPTMCVDWSNIRQLLLFPNSTIGDSGVPALPSLTMRRMR
ESVSRMPVSSQHRYSTPHAFTFNTSSPSSEGSLSQRQRSTSTPNVHMVSTTLPVDSRMIEDAIRSHSESASPS
ALSSSPNNLSPTGWSQPKTPVPAQRERAPVSGTQEKNKIRPRGQRDSSYYWEIEASEVMLSTRIGSGSFGTVY
KGKWHGDVAVKILKVVDPTPEQFQAFRNEVAVLRKTRHVNILLFMGYMTKDNLAIVTQWCEGSSLYKHLHVQE
TKFQMFQLIDIARQTAQGMDYLHAKNIIHRDMKSNNIFLHEGLTVKIGDFGLATVKSRWSGSQQVEQPTGSVL
WMAPEVIRMQDNNPFSFQSDVYSYGIVLYELMTGELPYSHINNRDQIIFMVGRGYASPDLSKLYKNCPKAMKR
LVADCVKKVKEERPLFPQILSSIELLQHSLPKINRSASEPSLHRAAHTEDINACTLTTSPRLPVFCGR*-[*IR*
*ES*]-MTEYKPTVRLATRDDVPRAVRTLAAAFADYPATRHTVDPDRHIERVTELQELFLTRVGLDIGKVWVADD
GAAVAVWTTPESVEAGAVFAEIGPRMAELSGSRLAAQQQMEGLLAPHRPKEPAWFLATVGVSPDHQGKGLGSA
VVLPGVEAAERAGVPAFLETSAPRNLPFYERLGFTVTADVECPKDRATWCMTRKPGA*

>DNA sequence
ATGGTGAGCAAGGGCGAGGAGCTGTTCACCGGGGTGGTGCCCATCCTGGTCGAGCTGGACGGCGACGTAAACG
GCCACAAGTTCAGCGTGTCCGGCGAGGGCGAGGGCGATGCCACCTACGGCAAGCTGACCCTGAAGTTCATCTG
CACCACCGGCAAGCTGCCCGTGCCCTGGCCCACCCTCGTGACCACCCTGACCTACGGCGTGCAGTGCTTCAGC
CGCTACCCCGACCACATGAAGCAGCACGACTTCTTCAAGTCCGCCATGCCCGAAGGCTACGTCCAGGAGCGCA
CCATCTTCTTCAAGGACGACGGCAACTACAAGACCCGCGCCGAGGTGAAGTTCGAGGGCGACACCCTGGTGAA
CCGCATCGAGCTGAAGGGCATCGACTTCAAGGAGGACGGCAACATCCTGGGGCACAAGCTGGAGTACAACTAC
AACAGCCACAACGTCTATATCATGGCCGACAAGCAGAAGAACGGCATCAAGGTGAACTTCAAGATCCGCCACA
ACATCGAGGACGGCAGCGTGCAGCTCGCCGACCACTACCAGCAGAACACCCCCATCGGCGACGGCCCCGTGCT
GCTGCCCGACAACCACTACCTGAGCACCCAGTCCGCCCTGAGCAAAGACCCCAACGAGAAGCGCGATCACATG
GTCCTGCTGGAGTTCGTGACCGCCGCCGGGATCACTCTCGGCATGGACGAGCTGTACAAGTCTGGGTCCGCAG
CCAGTAGCGCCGGAAGTATGATCAGTCTGATTGCGGCGTTAGCGGTAGATCGCGTTATCGGCATGGAAAACGC
CATGCCGTGGAACCTGCCTGCCGATCTCGCCTGGTTTAAACGCAACACCTTAAATAAACCCGTGATTATGGGC
CGCCATACCTGGGAATCAATCGGTCGTCCGTTGCCAGGACGCAAAAATATTATCCTCAGCAGTCAACCGGGTA
CGGACAAAAAAAAGAAAAAGAAAGATCGCGTAACGTGGGTGAAGTCGGTGGATGAAGCCATCGCGGCGTGTGG
TGACGTACCAGAAATCATGGTGATTGGCGGCGGTCGCGTTTATGAACAGTTCTTGCCAAAAGCGCAAAAACTG
TATCTGACGCATATCGACGCAGAAGTGGAAGGCGACACCCATTTCCCGGATTACGAGCCGGATGACTGGGAAT
CGGTATTCAGCGAATTCCACGATGCTGATGCGCAGAACTCTCACAGCTATTGCTTTGAGATTCTGGAGCGGCG
GAGTGCTGGTGGTAGTGCTGGTGGTAGTGCTGGTGGTAGTGCTGGTGGTAGTGCTGGTGGTCCTCGAAGCCTC
GAGATGGAGCACATACAGGGAGCTTGGAAGACGATCAGCAATGGTTTTGGATTCAAAGATGCCGTGTTTGATG
GCTCCAGCTGCATCTCTCCTACAATAGTTCAGCAGTTTGGCTATCAGCGCCGGGCATCAGATGATGGCAAACT
CACAGATCCTTCTAAGACAAGCAACACTATCCGTGTTTTCTTGCCGAACAAGCAAAGAACAGTGGTCAATGTG
CGAAATGGAATGAGCTTGCATGACTGCCTTATGAAAGCACTCAAGGTGAGGGGCCTGCAACCAGAGTGCTGTG
CAGTGTTCAGACTTCTCCACGAACACAAAGGTAAAAAAGCACGCTTAGATTGGAATACTGATGCTGCGTCTTT
GATTGGAGAAGAACTTCAAGTAGATTTCCTGGATCATGTTCCCCTCACAACACACAACTTTGCTCGGAAGACG
TTCCTGAAGCTTGCCTTCTGTGACATCTGTCAGAAATTCCTGCTCAATGGATTTCGATGTCAGACTTGTGGCT
ACAAATTTCATGAGCACTGTAGCACCAAAGTACCTACTATGTGTGTGGACTGGAGTAACATCAGACAACTCTT
ATTGTTTCCAAATTCCACTATTGGTGATAGTGGAGTCCCAGCACTACCTTCTTTGACTATGCGTCGTATGCGA
GAGTCTGTTTCCAGGATGCCTGTTAGTTCTCAGCACAGATATTCTACACCTCACGCCTTCACCTTTAACACCT
CCAGTCCCTCATCTGAAGGTTCCCTCTCCCAGAGGCAGAGGTCGACATCCACACCTAATGTCCACATGGTCAG
CACCACGCTGCCTGTGGACAGCAGGATGATTGAGGATGCAATTCGAAGTCACAGCGAATCAGCCTCACCTTCA
GCCCTGTCCAGTAGCCCCAACAATCTGAGCCCAACAGGCTGGTCACAGCCGAAAACCCCCGTGCCAGCACAAA
GAGAGCGGGCACCAGTATCTGGGACCCAGGAGAAAAACAAAATTAGGCCTCGTGGACAGAGAGATTCAAGCTA
TTATTGGGAAATAGAAGCCAGTGAAGTGATGCTGTCCACTCGGATTGGGTCAGGCTCTTTTGGAACTGTTTAT
AAGGGTAAATGGCACGGAGATGTTGCAGTAAAGATCCTAAAGGTTGTCGACCCAACCCCAGAGCAATTCCAGG

```
CCTTCAGGAATGAGGTGGCTGTTCTGCGCAAAACACGGCATGTGAACATTCTGCTTTTCATGGGGTACATGAC
AAAGGACAACCTGGCAATTGTGACCCAGTGGTGCGAGGGCAGCAGCCTCTACAAACACCTGCATGTCCAGGAG
ACCAAGTTTCAGATGTTCCAGCTAATTGACATTGCCCGGCAGACGGCTCAGGGAATGGACTATTTGCATGCAA
AGAACATCATCCATAGAGACATGAAATCCAACAATATATTTCTCCATGAAGGCTTAACAGTGAAAATTGGAGA
TTTTGGTTTGGCAACAGTAAAGTCACGCTGGAGTGGTTCTCAGCAGGTTGAACAACCTACTGGCTCTGTCCTC
TGGATGGCCCCAGAGGTGATCCGAATGCAGGATAACAACCCATTCAGTTTCCAGTCGGATGTCTACTCCTATG
GCATCGTATTGTATGAACTGATGACGGGGGAGCTTCCTTATTCTCACATCAACAACCGAGATCAGATCATCTT
CATGGTGGGCCGAGGATATGCCTCCCCAGATCTTAGTAAGCTATATAAGAACTGCCCCAAAGCAATGAAGAGG
CTGGTAGCTGACTGTGTGAAGAAAGTAAAGGAAGAGAGGCCTCTTTTTCCCCAGATCCTGTCTTCCATTGAGC
TGCTCCAACACTCTCTACCGAAGATCAACCGGAGCGCTTCCGAGCCATCCTTGCATCGGGCAGCCCACACTGA
GGATATCAATGCTTGCACGCTGACCACGTCCCCGAGGCTGCCTGTCTTCTGCGGCCGCTAGTCGACGGTACCG
CGGGCCCGGGATAAGTCAACTAACTTAAGCTAGCAACGGTTTCCCTCTAGCGGGATCAATTCCGCCCCCCCCC
CCTAACGTTACTGGCCGAAGCCGCTTGGAATAAGGCCGGTGTGCGTTTGTCTATATGTTATTTTCCACCATATTGCC
GTCTTTTGGCAATGTGAGGGCCCGGAAACCTGGCCCTGTCTTCTTGACGAGCATTCCTAGGGGTCTTTCCCCTCTCG
CCAAAGGAATGCAAGGTCTGTTGAATGTCGTGAAGGAAGCAGTTCCTCTGGAAGCTTCTTGAAGACAAACAACGTCT
GTAGCGACCCTTTGCAGGCAGCGGAACCCCCCACCTGGCGACAGGTGCCTCTGCGGCCAAAAGCCACGTGTATAAGA
TACACCTGCAAAGGCGGCACAACCCCAGTGCCACGTTGTGAGTTGGATAGTTGTGGAAAGAGTCAAATGGCTCTCCT
CAAGCGTATTCAACAAGGGGCTGAAGGATGCCCAGAAGGTACCCCATTGTATGGGATCTGATCTGGGGCCTCGGTGC
ACATGCTTTACATGTGTTTAGTCGAGGTTAAAAAACGTCTAGGCCCCCCGAACCACGGGGACGTGGTTTTCCTTTGA
AAAACACGATAATACCATGACCGAGTACAAGCCCACGGTGCGCCTCGCCACCCGCGACGACGTCCCCAGGGCCG
TACGCACCCTCGCCGCCGCGTTCGCCGACTACCCCGCCACGCGCCACACCGTCGATCCGGACCGCCACATCGA
GCGGGTCACCGAGCTGCAAGAACTCTTCCTCACGCGCGTCGGGCTCGACATCGGCAAGGTGTGGGTCGCGGAC
GACGGCGCCGCGGTGGCGGTCTGGACCACGCCGGAGAGCGTCGAAGCGGGGGCGGTGTTCGCCGAGATCGGCC
CGCGCATGGCCGAGTTGAGCGGTTCCCGGCTGGCCGCGCAGCAACAGATGGAAGGCCTCCTGGCGCCGCACCG
GCCCAAGGAGCCCGCGTGGTTCCTGGCCACCGTCGGCGTCTCGCCCGACCACCAGGGCAAGGGTCTGGGCAGC
GCCGTCGTGCTCCCCGGAGTGGAGGCGGCCGAGCGCGCCGGGGTGCCCGCCTTCCTGGAGACCTCCGCGCCCC
GCAACCTCCCCTTCTACGAGCGGCTCGGCTTCACCGTCACCGCCGACGTCGAGTGCCCGAAGGACCGCGCGAC
CTGGTGCATGACCCGCAAGCCCGGTGCCTGA
```

EGFP eDHFR K6-tag(69K6) cRaf(human) *IRES* Puro[R]

**pPBpuro-EGFP-eDHFR(69K6)**

>Amino acid sequence
MVSKGEELFTGVVPILVELDGDVNGHKFSVSGEGEGDATYGKLTLKFICTTGKLPVPWPTLVTTLTYGVQCFS
RYPDHMKQHDFFKSAMPEGYVQERTIFFKDDGNYKTRAEVKFEGDTLVNRIELKGIDFKEDGNILGHKLEYNY
NSHNVYIMADKQKNGIKVNFKIRHNIEDGSVQLADHYQQNTPIGDGPVLLPDNHYLSTQSALSKDPNEKRDHM
VLLEFVTAAGITLGMDELYKSGSAASSAGSMISLIAALAVDRVIGMENAMPWNLPADLAWFKRNTLNKPVIMG
RHTWESIGRPLPGRKNIILSSQPGTDKKKKKKDRVTWVKSVDEAIAACGDVPEIMVIGGGRVYEQFLPKAQKL
YLTHIDAEVEGDTHFPDYEPDDWESVFSEFHDADAQNSHSYCFEILERR-[*IRES*]-MTEYKPTVRLATRDDV
PRAVRTLAAAFADYPATRHTVDPDRHIERVTELQELFLTRVGLDIGKVWVADDGAAVAVWTTPESVEAGAVFA
EIGPRMAELSGSRLAAQQQMEGLLAPHRPKEPAWFLATVGVSPDHQGKGLGSAVVLPGVEAAERAGVPAFLET
SAPRNLPFYERLGFTVTADVECPKDRATWCMTRKPGA*

>DNA sequence
ATGGTGAGCAAGGGCGAGGAGCTGTTCACCGGGGTGGTGCCCATCCTGGTCGAGCTGGACGGCGACGTAAACG
GCCACAAGTTCAGCGTGTCCGGCGAGGGCGAGGGCGATGCCACCTACGGCAAGCTGACCCTGAAGTTCATCTG
CACCACCGGCAAGCTGCCCGTGCCCTGGCCCACCCTCGTGACCACCCTGACCTACGGCGTGCAGTGCTTCAGC
CGCTACCCCGACCACATGAAGCAGCACGACTTCTTCAAGTCCGCCATGCCCGAAGGCTACGTCCAGGAGCGCA
CCATCTTCTTCAAGGACGACGGCAACTACAAGACCCGCGCCGAGGTGAAGTTCGAGGGCGACACCCTGGTGAA
CCGCATCGAGCTGAAGGGCATCGACTTCAAGGAGGACGGCAACATCCTGGGGCACAAGCTGGAGTACAACTAC
AACAGCCACAACGTCTATATCATGGCCGACAAGCAGAAGAACGGCATCAAGGTGAACTTCAAGATCCGCCACA
ACATCGAGGACGGCAGCGTGCAGCTCGCCGACCACTACCAGCAGAACACCCCCATCGGCGACGGCCCCGTGCT
GCTGCCCGACAACCACTACCTGAGCACCCAGTCCGCCCTGAGCAAAGACCCCAACGAGAAGCGCGATCACATG
GTCCTGCTGGAGTTCGTGACCGCCGCCGGGATCACTCTCGGCATGGACGAGCTGTACAAGTCTGGGTCCGCAG
CCAGTAGCGCCGGAAGTATGATCAGTCTGATTGCGGCGTTAGCGGTAGATCGCGTTATCGGCATGGAAAACGC
CATGCCGTGGAACCTGCCTGCCGATCTCGCCTGGTTTAAACGCAACACCTTAAATAAACCCGTGATTATGGGC
CGCCATACCTGGGAATCAATCGGTCGTCCGTTGCCAGGACGCAAAAATATTATCCTCAGCAGTCAACCGGGTA
CGGACAAAAAAAAGAAAAAGAAAGATCGCGTAACGTGGGTGAAGTCGGTGGATGAAGCCATCGCGGCGTGTGG
TGACGTACCAGAAATCATGGTGATTGGCGGCGGTCGCGTTTATGAACAGTTCTTGCCAAAAGCGCAAAAACTG
TATCTGACGCATATCGACGCAGAAGTGGAAGGCGACACCCATTTCCCGGATTACGAGCCGGATGACTGGGAAT
CGGTATTCAGCGAATTCCACGATGCTGATGCGCAGAACTCTCACAGCTATTGCTTTGAGATTCTGGAGCGGCG
GTAACCCGGGATAAGTCAACTAACTTAAGCTAGCAACGGTTTCCCTCTAGCGGGATCAATTCCG*CCCCCCCCC*
*CCTAACGTTACTGGCCGAAGCCGCTTGGAATAAGGCCGGTGTGCGTTTGTCTATATGTTATTTTCCACCATAT*
*TGCCGTCTTTTGGCAATGTGAGGGCCCGGAAACCTGGCCCTGTCTTCTTGACGAGCATTCCTAGGGGTCTTTC*
*CCCTCTCGCCAAAGGAATGCAAGGTCTGTTGAATGTCGTGAAGGAAGCAGTTCCTCTGGAAGCTTCTTGAAGA*
*CAAACAACGTCTGTAGCGACCCTTTGCAGGCAGCGGAACCCCCCACCTGGCGACAGGTGCCTCTGCGGCCAAA*
*AGCCACGTGTATAAGATACACCTGCAAAGGCGGCACAACCCCAGTGCCACGTTGTGAGTTGGATAGTTGTGGA*
*AAGAGTCAAATGGCTCTCCTCAAGCGTATTCAACAAGGGGCTGAAGGATGCCCAGAAGGTACCCCATTGTATG*
*GGATCTGATCTGGGGCCTCGGTGCACATGCTTTACATGTGTTTAGTCGAGGTTAAAAAACGTCTAGGCCCCCC*
*GAACCACGGGGACGTGGTTTTCCTTTGAAAAACACGAT*AATACCATGACCGAGTACAAGCCCACGGTGCGCCT
CGCCACCCGCGACGACGTCCCCAGGGCCGTACGCACCCTCGCCGCCGCGTTCGCCGACTACCCCGCCACGCGC
CACACCGTCGATCCGGACCGCCACATCGAGCGGGTCACCGAGCTGCAAGAACTCTTCCTCACGCGCGTCGGGC
TCGACATCGGCAAGGTGTGGGTCGCGGACGACGGCGCCGCGGTGGCGGTCTGGACCACGCCGGAGAGCGTCGA
AGCGGGGGCGGTGTTCGCCGAGATCGGCCCGCGCATGGCCGAGTTGAGCGGTTCCCGGCTGGCCGCGCAGCAA
CAGATGGAAGGCCTCCTGGCGCCGCACCGGCCCAAGGAGCCCGCGTGGTTCCTGGCCACCGTCGGCGTCTCGC
CCGACCACCAGGGCAAGGGTCTGGGCAGCGCCGTCGTGCTCCCCGGAGTGGAGGCGGCCGAGCGCGCCGGGGT
GCCCGCCTTCCTGGAGACCTCCGCGCCCCGCAACCTCCCCTTCTACGAGCGGCTCGGCTTCACCGTCACCGCC
GACGTCGAGTGCCCGAAGGACCGCGCGACCTGGTGCATGACCCGCAAGCCCGGTGCCTGA

EGFP eDHFR K6-tag(69K6) *IRES* Puro$^R$

#### **pPBbsr-ERK-KTR-mKO**

>Amino acid sequence

MKGRKPRDLELPLSPSLLGGQGPERTPGSGTSSGLQAPGPALSPSKRSGLEDPATPSKKPRTPSVSSRLERLT
LQSSFQFPSGGRMVSVIKPEMKMRYYMDGSVNGHEFTIEGEGTGRPYEGHQEMTLRVTMAKGGPMPFAFDLVS
HVFCYGHRPFTKYPEEIPDYFKQAFPEGLSWERSLEFEDGGSASVSAHISLRGNTFYHKSKFTGVNFPADGPI
MQNQSVDWEPSTEKITASDGVLKGDVTMYLKLEGGGNHKCQFKTTYKAAKKILKMPGSHYISHRLVRKTEGNI
TELVEDAVAHS*-[*IRES*]-MVMKTFNISQQDLELVEVATEKITMLYEDNKHHVGAAIRTKTGEIISAVHIEA
YIGRVTVCAEAIAIGSAVSNGQKDFDTIVAVRHPYSDEVDRSIRVVSPCGMCRELISDYAPDCFVLIEMNGKL
VKTTIEELIPLKYTRN*

>DNA sequence

ATGAAAGGCCGCAAGCCTCGCGATCTGGAGTTACCTCTCAGCCCTAGCCTCCTGGGGGGACAGGGCCCTGAGC
GCACACCCGGGTCCGGCACCTCTAGTGGTTTGCAGGCTCCGGGACCTGCCCTGTCACCCTCCAAGCGCAGCGG
CCTGGAGGATCCTGCCACACCCAGCAAAAAACCAAGGACCCCCTCAGTCAGCTCGCGGTTAGAGCGACTGACG
CTCCAGTCTTCTTTCCAATTCCCGTCCGGCGGCCGCATGGTGAGCGTGATCAAGCCCGAGATGAAGATGAGGT
ACTACATGGACGGCTCCGTCAATGGGCATGAGTTCACAATCGAGGGTGAGGGCACAGGCAGACCTTACGAGGG
ACATCAGGAGATGACACTGCGCGTCACAATGGCCAAGGGCGGGCCAATGCCTTTCGCCTTCGACCTGGTGTCC
CACGTGTTCTGTTACGGCCACAGACCTTTTACTAAATATCCAGAAGAGATCCCAGACTATTTCAAGCAGGCCT
TTCCTGAGGGCCTGTCCTGGGAGAGGTCCCTGGAGTTCGAGGACGGCGGCTCCGCCTCCGTGAGCGCCCACAT
CAGCCTGAGGGGCAACACCTTCTACCACAAGTCCAAGTTCACCGGCGTGAACTTCCCCGCCGACGGCCCCATC
ATGCAGAACCAGAGCGTGGACTGGGAGCCCTCCACCGAGAAGATCACCGCCAGCGACGGCGTGCTGAAGGGCG
ACGTGACCATGTACCTGAAGCTGGAGGGCGGCGGCAACCACAAGTGCCAGTTCAAGACCACCTACAAGGCCGC
CAAGAAGATCCTGAAGATGCCCGGCAGCCACTACATCAGCCACAGGCTGGTGAGGAAGACCGAGGGCAACATC
ACCGAGCTGGTGGAGGACGCCGTGGCCCACTCCTAAGTCGACGGGCCGCGGTAACAATTGTTAACTAACTTAA
GCTAGCAACGGTTTCCCTCTAGCGGGATCAATTCCG*CCCCCCCCCCTAACGTTACTGGCCGAAGCCGCTTGGAA*
*TAAGGCCGGTGTGCGTTTGTCTATATGTTATTTTCCACCATATTGCCGTCTTTTGGCAATGTGAGGGCCCGGAAACC*
*TGGCCCTGTCTTCTTGACGAGCATTCCTAGGGGTCTTTCCCCTCTCGCCAAAGGAATGCAAGGTCTGTTGAATGTCG*
*TGAAGGAAGCAGTTCCTCTGGAAGCTTCTTGAAGACAAACAACGTCTGTAGCGACCCTTTGCAGGCAGCGGAACCCC*
*CCACCTGGCGACAGGTGCCTCTGCGGCCAAAAGCCACGTGTATAAGATACACCTGCAAAGGCGGCACAACCCCAGTG*
*CCACGTTGTGAGTTGGATAGTTGTGGAAAGAGTCAAATGGCTCTCCTCAAGCGTATTCAACAAGGGGCTGAAGGATG*
*CCCAGAAGGTACCCCATTGTATGGGATCTGATCTGGGGCCTCGGTGCACATGCTTTACATGTGTTTAGTCGAGGTTA*
*AAAAACGTCTAGGCCCCCCGAACCACGGGGACGTGGTTTTCCTTTGAAAAACACGAT*AATACCATGGTCATGAAAA
CATTTAACATTTCTCAACAAGATCTAGAATTAGTAGAAGTAGCGACAGAGAAGATTACAATGCTTTATGAGGA
TAATAAACATCATGTGGGAGCGGCAATTCGTACGAAAACAGGAGAAATCATTTCGGCAGTACATATTGAAGCG
TATATAGGACGAGTAACTGTTTGTGCAGAAGCCATTGCGATTGGTAGTGCAGTTTCGAATGGACAAAAGGATT
TTGACACGATTGTAGCTGTTAGACACCCTTATTCTGACGAAGTAGATAGAAGTATTCGAGTGGTAAGTCCTTG
TGGTATGTGTAGGGAGTTGATTTCAGACTATGCACCAGATTGTTTTGTGTTAATAGAAATGAATGGCAAGTTA
GTCAAAACTACGATTGAAGAACTCATTCCACTCAAATATACCCGAAATTAA

ERK-KTR[S26,S27] mKusabira-Orange *IRES* Bsr$^{R}$ (blasticidin *S*-deaminase)

**pCAGGS-mNG-eDHFR(69K6)-Gαq**

>Amino acid sequence
MVSKGEEDNMASLPATHELHIFGSINGVDFDMVGQGTGNPNDGYEELNLKSTKGDLQFSPWILVPHIGYGFHQ
YLPYPDGMSPFQAAMVDGSGYQVHRTMQFEDGASLTVNYRYTYEGSHIKGEAQVKGTGFPADGPVMTNSLTAA
DWCRSKKTYPNDKTIISTFKWSYTTGNGKRYRSTARTTYTFAKPMAANYLKNQPMYVFRKTELKHSKTELNFK
EWQKAFTDVMGMDELYKSGSAASSAGSMISLIAALAVDRVIGMENAMPWNLPADLAWFKRNTLNKPVIMGRHT
WESIGRPLPGRKNIILSSQPGTDKKKKKKDRVTWVKSVDEAIAACGDVPEIMVIGGGRVYEQFLPKAQKLYLT
HIDAEVEGDTHFPDYEPDDWESVFSEFHDADAQNSHSYCFEILERRLYSGGGGSGGGGSGGGGSSAGASLYKA
RGAAAGAGGAGRSGKKGKKGKKGTTMTLESIMASSLSEEAKEARRINDEIERQLRRDKRDARRELKLLLLGTG
ESGKSTFIKQMRIIHGSGYSDEDKRGFTKLVYQNIFTAMQAMIRAMDTLKIPYKYEHNKAHAQLVREVDVEKV
SAFENPYVDAIKSLWNDPGIQECYDRRREYQLSDSTKYYLNDLDRVADPSYLPTQQDVLRVRVPTTGIIEYPF
DLQSVIFRMVDVGGLRSERRKWIHCFENVTSIMFLVALSEYDQVLVESDNENRMEESKALFRTIITYPWFQNS
SVILFLNKKDLLEEKIMYSHLVDYFPEYDGPQRDAQAAREFILKMFVDLNPDSDKIIYSHFTCATDTENIRFV
FAAVKDTILQLNLKEYNLV*

>DNA sequence
ATGGTGAGCAAGGGCGAGGAGGATAACATGGCCTCTCTCCCAGCGACACATGAGTTACACATCTTTGGCTCCA
TCAACGGTGTGGACTTTGACATGGTGGGTCAGGGCACCGGCAATCCAAATGATGGTTATGAGGAGTTAAACCT
GAAGTCCACCAAGGGTGACCTCCAGTTCTCCCCCTGGATTCTGGTCCCTCATATCGGGTATGGCTTCCATCAG
TACCTGCCCTACCCTGACGGGATGTCGCCTTTCCAGGCCGCCATGGTAGATGGCTCCGGCTACCAAGTCCATC
GCACAATGCAGTTTGAAGATGGTGCCTCCCTTACTGTTAACTACCGCTACACCTACGAGGGAAGCCACATCAA
AGGAGAGGCCCAGGTGAAGGGGACTGGTTTCCCTGCTGACGGTCCTGTGATGACCAACTCGCTGACCGCTGCG
GACTGGTGCAGGTCGAAGAAGACTTACCCCAACGACAAAACCATCATCAGTACCTTTAAGTGGAGTTACACCA
CTGGAAATGGCAAGCGCTACCGGAGCACTGCGCGGACCACCTACACCTTTGCCAAGCCAATGGCGGCTAACTA
TCTGAAGAACCAGCCGATGTACGTGTTCCGTAAGACGGAGCTCAAGCACTCCAAGACCGAGCTCAACTTCAAG
GAGTGGCAAAAGGCCTTTACCGATGTGATGGGCATGGACGAGCTCTACAAGTCTGGGTCCGCAGCCAGTAGCG
CCGGAAGTATGATCAGTCTGATTGCGGCGTTAGCGGTAGATCGCGTTATCGGCATGGAAAACGCCATGCCGTG
GAACCTGCCTGCCGATCTCGCCTGGTTTAAACGCAACACCTTAAATAAACCCGTGATTATGGGCCGCCATACC
TGGGAATCAATCGGTCGTCCGTTGCCAGGACGCAAAAATATTATCCTCAGCAGTCAACCGGGTACGGACAAAA
AAAAGAAAAAGAAAGATCGCGTAACGTGGGTGAAGTCGGTGGATGAAGCCATCGCGGCGTGTGGTGACGTACC
AGAAATCATGGTGATTGGCGGCGGTCGCGTTTATGAACAGTTCTTGCCAAAAGCGCAAAAACTGTATCTGACG
CATATCGACGCAGAAGTGGAAGGCGACACCCATTTCCCGGATTACGAGCCGGATGACTGGGAATCGGTATTCA
GCGAATTCCACGATGCTGATGCGCAGAACTCTCACAGCTATTGCTTTGAGATTCTGGAGCGGCGGCTGTACTC
TGGTGGCGGAGGCTCGGGCGGAGGTGGGTCGGGTGGCGGCGGATCTTCAGCTGGAGCTTCGTTGTACAAGGCA
AGAGGAGCCGCTGCGGGTGCCGGAGGCGCTGGTAGGAGCGGCAAAAAGGGGAAGAAAGGGAAAAAGGGCACCA
CAATGACTCTGGAGTCCATCATGGCATCCTCCCTGAGCGAGGAGGCCAAGGAAGCCCGGAGGATCAACGACGA
GATCGAGCGGCAGCTGCGCAGGGACAAGCGCGACGCCCGCCGGGAGCTCAAGCTGCTGCTGCTGGGGACAGGG
GAGAGTGGCAAGTCGACCTTCATCAAGCAGATGAGGATCATCCACGGGTCGGGCTACTCTGACGAAGACAAGC
GCGGCTTCACCAAGCTGGTGTATCAGAACATCTTCACGGCCATGCAGGCCATGATCAGAGCGATGGACACACT
CAAGATCCCATACAAGTATGAACACAATAAGGCTCATGCACAATTGGTTCGAGAGGTTGATGTGGAGAAGGTG
TCTGCTTTTGAGAATCCATATGTAGATGCAATAAAGAGCTTGTGGAATGATCCTGGAATCCAGGAGTGCTACG
ACAGACGACGGGAATATCAGTTATCTGACTCTACCAAATACTATCTGAATGACTTGGACCGTGTAGCCGACCC
TTCCTATCTGCCTACACAACAAGACGTGCTTAGAGTTCGAGTCCCCACTACAGGGATCATCGAATACCCCTTT
GACTTACAAAGTGTCATTTTCAGAATGGTCGATGTAGGGGGCCTGAGGTCAGAGAGAAGAAAATGGATACACT
GCTTTGAAAATGTCACCTCCATCATGTTTCTAGTAGCGCTTAGCGAATATGATCAAGTTCTTGTGGAGTCAGA
CAATGAGAACCGCATGGAGGAGAGCAAAGCACTCTTTAGAACAATTATCACCTACCCCTGGTTCCAGAACTCC
TCTGTGATTCTGTTCTTAAACAAGAAAGATCTTCTAGAGGAGAAAATCATGTATTCCCACCTAGTCGACTACT
TCCCAGAATATGATGGACCCCAGAGAGATGCCCAGGCAGCTCGAGAATTCATCCTGAAAATGTTCGTGGACCT
GAACCCCGACAGTGACAAAATCATCTACTCCCACTTCACGTGCGCCACAGATACCGAGAACATCCGCTTCGTC
TTTGCAGCCGTCAAGGACACCATCCTGCAGCTGAACCTGAAGGAGTACAATCTGGTCTAA

mNeonGreen eDHFR K6-tag(69K6) Gαq (human Gαq with C9S/C10S/Q209L mutations)[S3,S28] Lysine-rich linker[S29]

**pCAGGS-mNG-eDHFR(69K6)-Gαq$_{L254A}$**

```
>Amino acid sequence
MVSKGEEDNMASLPATHELHIFGSINGVDFDMVGQGTGNPNDGYEELNLKSTKGDLQFSPWILVPHIGYGFHQ
YLPYPDGMSPFQAAMVDGSGYQVHRTMQFEDGASLTVNYRYTYEGSHIKGEAQVKGTGFPADGPVMTNSLTAA
DWCRSKKTYPNDKTIISTFKWSYTTGNGKRYRSTARTTYTFAKPMAANYLKNQPMYVFRKTELKHSKTELNFK
EWQKAFTDVMGMDELYKSGSAASSAGSMISLIAALAVDRVIGMENAMPWNLPADLAWFKRNTLNKPVIMGRHT
WESIGRPLPGRKNIILSSQPGTDKKKKKKDRVTWVKSVDEAIAACGDVPEIMVIGGGRVYEQFLPKAQKLYLT
HIDAEVEGDTHFPDYEPDDWESVFSEFHDADAQNSHSYCFEILERRLYSGGGGSGGGGSGGGGSSAGASLYKA
RGAAAGAGGAGRSGKKGKKGKKGTTMTLESIMASSLSEEAKEARRINDEIERQLRRDKRDARRELKLLLLGTG
ESGKSTFIKQMRIIHGSGYSDEDKRGFTKLVYQNIFTAMQAMIRAMDTLKIPYKYEHNKAHAQLVREVDVEKV
SAFENPYVDAIKSLWNDPGIQECYDRRREYQLSDSTKYYLNDLDRVADPSYLPTQQDVLRVRVPTTGIIEYPF
DLQSVIFRMVDVGGLRSERRKWIHCFENVTSIMFLVALSEYDQVLVESDNENRMEESKAAFRTIITYPWFQNS
SVILFLNKKDLLEEKIMYSHLVDYFPEYDGPQRDAQAAREFILKMFVDLNPDSDKIIYSHFTCATDTENIRFV
FAAVKDTILQLNLKEYNLV*

>DNA sequence
ATGGTGAGCAAGGGCGAGGAGGATAACATGGCCTCTCTCCCAGCGACACATGAGTTACACATCTTTGGCTCCA
TCAACGGTGTGGACTTTGACATGGTGGGTCAGGGCACCGGCAATCCAAATGATGGTTATGAGGAGTTAAACCT
GAAGTCCACCAAGGGTGACCTCCAGTTCTCCCCCTGGATTCTGGTCCCTCATATCGGGTATGGCTTCCATCAG
TACCTGCCCTACCCTGACGGGATGTCGCCTTTCCAGGCCGCCATGGTAGATGGCTCCGGCTACCAAGTCCATC
GCACAATGCAGTTTGAAGATGGTGCCTCCCTTACTGTTAACTACCGCTACACCTACGAGGGAAGCCACATCAA
AGGAGAGGCCCAGGTGAAGGGGACTGGTTTCCCTGCTGACGGTCCTGTGATGACCAACTCGCTGACCGCTGCG
GACTGGTGCAGGTCGAAGAAGACTTACCCCAACGACAAAACCATCATCAGTACCTTTAAGTGGAGTTACACCA
CTGGAAATGGCAAGCGCTACCGGAGCACTGCGCGGACCACCTACACCTTTGCCAAGCCAATGGCGGCTAACTA
TCTGAAGAACCAGCCGATGTACGTGTTCCGTAAGACGGAGCTCAAGCACTCCAAGACCGAGCTCAACTTCAAG
GAGTGGCAAAAGGCCTTTACCGATGTGATGGGCATGGACGAGCTCTACAAGTCTGGGTCCGCAGCCAGTAGCG
CCGGAAGTATGATCAGTCTGATTGCGGCGTTAGCGGTAGATCGCGTTATCGGCATGGAAAACGCCATGCCGTG
GAACCTGCCTGCCGATCTCGCCTGGTTTAAACGCAACACCTTAAATAAACCCGTGATTATGGGCCGCCATACC
TGGGAATCAATCGGTCGTCCGTTGCCAGGACGCAAAAATATTATCCTCAGCAGTCAACCGGGTACGGACAAAA
AAAAGAAAAAGAAAGATCGCGTAACGTGGGTGAAGTCGGTGGATGAAGCCATCGCGGCGTGTGGTGACGTACC
AGAAATCATGGTGATTGGCGGCGGTCGCGTTTATGAACAGTTCTTGCCAAAAGCGCAAAAACTGTATCTGACG
CATATCGACGCAGAAGTGGAAGGCGACACCCATTTCCCGGATTACGAGCCGGATGACTGGGAATCGGTATTCA
GCGAATTCCACGATGCTGATGCGCAGAACTCTCACAGCTATTGCTTTGAGATTCTGGAGCGGCGGCTGTACTC
TGGTGGCGGAGGCTCGGGCGGAGGTGGGTCGGGTGGCGGCGGATCTTCAGCTGGAGCTTCGTTGTACAAGGCA
AGAGGAGCCGCTGCGGGTGCCGGAGGCGCTGGTAGGAGCGGCAAAAAGGGGAAGAAAGGGAAAAAGGGCACCA
CAATGACTCTGGAGTCCATCATGGCATCCTCCCTGAGCGAGGAGGCCAAGGAAGCCCGGAGGATCAACGACGA
GATCGAGCGGCAGCTGCGCAGGGACAAGCGCGACGCCCGCCGGGAGCTCAAGCTGCTGCTGCTGGGGACAGGG
GAGAGTGGCAAGTCGACCTTCATCAAGCAGATGAGGATCATCCACGGGTCGGGCTACTCTGACGAAGACAAGC
GCGGCTTCACCAAGCTGGTGTATCAGAACATCTTCACGGCCATGCAGGCCATGATCAGAGCGATGGACACACT
CAAGATCCCATACAAGTATGAACACAATAAGGCTCATGCACAATTGGTTCGAGAGGTTGATGTGGAGAAGGTG
TCTGCTTTTGAGAATCCATATGTAGATGCAATAAAGAGCTTGTGGAATGATCCTGGAATCCAGGAGTGCTACG
ACAGACGACGGGAATATCAGTTATCTGACTCTACCAAATACTATCTGAATGACTTGGACCGTGTAGCCGACCC
TTCCTATCTGCCTACACAACAAGACGTGCTTAGAGTTCGAGTCCCCACTACAGGGATCATCGAATACCCCTTT
GACTTACAAAGTGTCATTTTCAGAATGGTCGATGTAGGGGGCCTGAGGTCAGAGAGAAGAAAATGGATACACT
GCTTTGAAAATGTCACCTCCATCATGTTTCTAGTAGCGCTTAGCGAATATGATCAAGTTCTTGTGGAGTCAGA
CAATGAGAACCGCATGGAGGAGAGCAAAGCAGCCTTTAGAACAATTATCACCTACCCCTGGTTCCAGAACTCC
TCTGTGATTCTGTTCTTAAACAAGAAAGATCTTCTAGAGGAGAAAATCATGTATTCCCACCTAGTCGACTACT
TCCCAGAATATGATGGACCCCAGAGAGATGCCCAGGCAGCTCGAGAATTCATCCTGAAAATGTTCGTGGACCT
GAACCCCGACAGTGACAAAATCATCTACTCCCACTTCACGTGCGCCACAGATACCGAGAACATCCGCTTCGTC
TTTGCAGCCGTCAAGGACACCATCCTGCAGCTGAACCTGAAGGAGTACAATCTGGTCTAA
```

mNeonGreen eDHFR K6-tag(69K6) Gαq$_{L254A}$ (human Gαq with C9S/C10S/Q209L/L254A mutations)[S3,S28,S30] Lysine-rich linker[S29]

**pCAGGS-R-GECO**

>Amino acid sequence
MVDSSRRKWNKAGHAVRAIGRLSSPVVSERMYPEDGALKSEIKKGLRLKDGGHYAAEVKTTYKAKKPVQLPGA
YIVDIKLDIVSHNEDYTIVEQCERAEGRHSTGGMDELYKGGTGGSLVSKGEEDNRAIVKEFMRFKLHMEGSVN
GHEFEIEGEGEGRPYEAFQTAKLKVTKGGPLPFAWDILSPQLMYGSKAYIKHPADIPDYFKLSFPEGFRWERV
MNFEDGGIIHVSQDTSLQDGVFIYKVKLRGTNFPPDGPVMQKKTMGWEATRDQLTEEQIAEFKEAFSLFDKDG
DGTMTTKELGTVMRSLGQNPTEAELQDMINEVDADGDGTFDFPEFLTMMARKMNDTDSEEEIREAFRVFDKDG
NGYIGAAELRHVMTDLGEKITDEEVDEMIRVADIDGDGQVNYEEFVQMMTAK*

>DNA sequence
ATGGTCGACTCATCACGTCGTAAGTGGAATAAGGCAGGTCACGCAGTCAGAGCTATAGGTCGGCTGAGCTCAC
CCGTGGTTTCCGAGCGGATGTACCCCGAGGACGGCGCCCTGAAGAGCGAGATCAAGAAGGGGCTGAGGCTGAA
GGACGGCGGCCACTACGCCGCCGAGGTCAAGACCACCTACAAGGCCAAGAAGCCCGTGCAGCTGCCCGGCGCC
TACATCGTCGACATCAAGTTGGACATCGTGTCCCACAACGAGGACTACACCATCGTGGAACAGTGCGAACGCG
CCGAGGGCCGCCACTCCACCGGCGGCATGGACGAGCTGTACAAGGGAGGTACAGGCGGGAGTCTGGTGAGCAA
GGGCGAGGAGGATAACAGGGCCATCGTCAAGGAGTTCATGCGCTTCAAGTTGCACATGGAGGGCTCCGTGAAC
GGCCACGAGTTCGAGATCGAGGGCGAGGGCGAGGGCCGCCCCTACGAGGCCTTTCAGACCGCTAAGCTGAAGG
TGACCAAGGGTGGCCCCCTGCCCTTCGCATGGGACATCCTGTCCCCTCAGTTGATGTACGGCTCCAAGGCCTA
CATTAAGCACCCAGCCGACATCCCCGACTACTTCAAGCTGTCCTTCCCCGAGGGCTTCAGGTGGGAGCGCGTG
ATGAACTTCGAGGACGGCGGCATTATTCACGTTAGCCAGGACACCTCCCTGCAGGACGGCGTATTCATCTACA
AGGTGAAGCTGCGCGGCACCAACTTCCCCCCCGACGGCCCCGTAATGCAGAAGAAGACCATGGGCTGGGAGGC
TACGCGTGACCAACTGACTGAAGAGCAGATCGCAGAATTTAAAGAGGCTTTCTCCCTATTTGACAAGGACGGG
GATGGGACGATGACAACCAAGGAGCTGGGGACGGTGATGCGGTCTCTGGGGCAGAACCCCACAGAAGCAGAGC
TGCAGGACATGATCAATGAAGTAGATGCCGACGGTGACGGCACATTCGACTTCCCTGAGTTCCTGACGATGAT
GGCAAGAAAAATGAATGACACAGACAGTGAAGAGGAAATTAGAGAAGCGTTCCGCGTGTTTGATAAGGACGGC
AATGGCTACATCGGCGCAGCAGAGCTTCGCCACGTGATGACAGACCTTGGAGAGAAGATAACAGATGAGGAGG
TTGATGAAATGATCAGGGTAGCAGACATCGATGGGGATGGTCAGGTAAACTATGAAGAGTTTGTACAAATGAT
GACAGCGAAGTAA

R-GECO1.2[S31]

**pPBpuro-miR-eDHFR(69K6)-G$\alpha$s**

>Amino acid sequence
MVAGHASGSPAFGTASHSNCEHEEIHLAGSIQPHGALLVVSEHDHRVIQASANAAEFLNLGSVLGVPLAEIDG
DLLIKILPHLDPTAEGMPVAVRCRIGNPSTEYCGLMHRPPEGGLIIELERAGPSIDLSGTLAPALERIRTAGS
LRALCDDTVLLFQQCTGYDRVMVYRFDEQGHGLVFSECHVPGLESYFGNRYPSSTVPQMARQLYVRQRVRVLV
DVTYQPVPLEPRLSPLTGRDLDMSGCFLRSMSPCHLQFLKDMGVRATLAVSLVVGGKLWGLVVCHHYLPRFIR
FELRAICKRLAERIATRITALESLYKSGSAASSAGSMISLIAALAVDRVIGMENAMPWNLPADLAWFKRNTLN
KPVIMGRHTWESIGRPLPGRKNIILSSQPGTDKKKKKKDRVTWVKSVDEAIAACGDVPEIMVIGGGRVYEQFL
PKAQKLYLTHIDAEVEGDTHFPDYEPDDWESVFSEFHDADAQNSHSYCFEILERRLYKSGLRSRQGSGAGSGA
GSGAGSGAGSGAPRAQASNSAVDGTAGPGMGSLGNSKTEDQRNEEKAQREANKKIEKQLQKDKQVYRATHRLL
LLGAGESGKSTIVKQMRILHVNGFNGDEKATKVQDIKNNLKEAIETIVAAMSNLVPPVELANPENQFRVDYIL
SVMNVPNFDFPPEFYEHAKALWEDEGVRACYERSNEYQLIDCAQYFLDKIDVIKQADYVPSDQDLLRCRVLTS
GIFETKFQVDKVNFHMFDVGGQRDERRKWIQCFNDVTAIIFVVASSSYNMVIREDNQTNRLQEALNLFKSIWN
NRWLRTISVILFLNKQDLLAEKVLAGKSKIEDYFPEFARYTTPEDATPEPGEDPRVTRAKYFIRDEFLRISTA
SGDGRHYCYPHFTCAVDTENIRRVFNDCRDIIQRMHLRQYELL*-[*IRES*]-MTEYKPTVRLATRDDVPRAVR
TLAAAFADYPATRHTVDPDRHIERVTELQELFLTRVGLDIGKVWVADDGAAVAVWTTPESVEAGAVFAEIGPR
MAELSGSRLAAQQQMEGLLAPHRPKEPAWFLATVGVSPDHQGKGLGSAVVLPGVEAAERAGVPAFLETSAPRN
LPFYERLGFTVTADVECPKDRATWCMTRKPGA*

>DNA sequence
ATGGTAGCAGGTCATGCCTCTGGCAGCCCCGCATTCGGGACCGCCTCTCATTCGAATTGCGAACATGAAGAGA
TCCACCTCGCCGGCTCGATCCAGCCGCATGGCGCGCTTCTGGTCGTCAGCGAACATGATCATCGCGTCATCCA
GGCCAGCGCCAACGCCGCGGAATTTCTGAATCTCGGAAGCGTACTCGGCGTTCCGCTCGCCGAGATCGACGGC
GATCTGTTGATCAAGATCCTGCCGCATCTCGATCCCACCGCCGAAGGCATGCCGGTCGCGGTGCGCTGCCGGA
TCGGCAATCCCTCTACGGAGTACTGCGGTCTGATGCATCGGCCTCCGGAAGGCGGGCTGATCATCGAACTCGA
ACGTGCCGGCCCGTCGATCGATCTGTCAGGCACGCTGGCGCCGGCGCTGGAGCGGATCCGCACGGCGGGTTCA
CTGCGCGCGCTGTGCGATGACACCGTGCTGCTGTTTCAGCAGTGCACCGGCTACGACCGGGTGATGGTGTATC
GTTTCGATGAGCAAGGCCACGGCCTGGTATTCTCCGAGTGCCATGTGCCTGGGCTCGAATCCTATTTCGGCAA
CCGCTATCCGTCGTCGACTGTCCCGCAGATGGCGCGGCAGCTGTACGTGCGGCAGCGCGTCCGCGTGCTGGTC
GACGTCACCTATCAGCCGGTGCCGCTGGAGCCGCGGCTGTCGCCGCTGACCGGGCGCGATCTCGACATGTCGG
GCTGCTTCCTGCGCTCGATGTCGCCGTGCCATCTGCAGTTCCTGAAGGACATGGGCGTGCGCGCCACCCTGGC
GGTGTCGCTGGTGGTCGGCGGCAAGCTGTGGGGCCTGGTTGTCTGTCACCATTATCTGCCGCGCTTCATCCGT
TTCGAGCTGCGGGCGATCTGCAAACGGCTCGCCGAAAGGATCGCGACGCGGATCACCGCGCTTGAGAGCCTCT
ACAAGTCTGGGTCCGCAGCCAGTAGCGCCGGAAGTATGATCAGTCTGATTGCGGCGTTAGCGGTAGATCGCGT
TATCGGCATGGAAAACGCCATGCCGTGGAACCTGCCTGCCGATCTCGCCTGGTTTAAACGCAACACCTTAAAT
AAACCCGTGATTATGGGCCGCCATACCTGGGAATCAATCGGTCGTCCGTTGCCAGGACGCAAAAATATTATCC
TCAGCAGTCAACCGGGTACGGACAAAAAAAAGAAAAAGAAAGATCGCGTAACGTGGGTGAAGTCGGTGGATGA
AGCCATCGCGGCGTGTGGTGACGTACCAGAAATCATGGTGATTGGCGGCGGTCGCGTTTATGAACAGTTCTTG
CCAAAAGCGCAAAAACTGTATCTGACGCATATCGACGCAGAAGTGGAAGGCGACACCCATTTCCCGGATTACG
AGCCGGATGACTGGGAATCGGTATTCAGCGAATTCCACGATGCTGATGCGCAGAACTCTCACAGCTATTGCTT
TGAGATTCTGGAGCGGCGGCTGTACAAGTCCGGACTCAGATCTCGACAAGGTAGTGGTGCTGGCTCTGGTGCT
GGTAGTGGCGCTGGTTCCGGTGCTGGCTCTGGCGCGCCTCGAGCTCAAGCTTCGAATTCTGCAGTCGACGGTA
CCGCGGGCCCGGGTATGGGCAGCCTCGGCAACAGTAAGACCGAGGACCAGCGCAACGAGGAGAAGGCGCAGCG
CGAGGCCAACAAAAAGATCGAGAAGCAGCTGCAGAAGGACAAGCAGGTCTACCGGGCCACGCACCGCCTGCTG
CTGCTGGGTGCTGGAGAGTCTGGCAAAAGCACCATTGTGAAGCAGATGAGGATCCTACATGTTAATGGGTTTA
ACGGAGACGAGAAGGCCACCAAAGTGCAGGACATCAAAAACAACCTGAAGGAGGCCATTGAAACCATTGTGGC
CGCCATGAGCAACCTGGTGCCCCCCGTGGAGCTGGCCAACCCTGAGAACCAGTTCAGAGTGGACTACATTCTG
AGCGTGATGAACGTGCCAAACTTTGACTTCCCACCTGAATTCTATGAGCATGCCAAGGCTCTGTGGGAGGATG
AGGGAGTTCGTGCCTGCTACGAGCGCTCCAACGAGTACCAGCTGATCGACTGTGCCCAGTACTTCCTGGACAA
GATTGATGTGATCAAGCAGGCCGACTACGTGCCAAGTGACCAGGACCTGCTTCGCTGCCGCGTCCTGACCTCT
GGAATCTTTGAGACCAAGTTCCAGGTGGACAAAGTCAACTTCCACATGTTCGATGTGGGCGGCCAGCGCGATG
AACGCCGCAAGTGGATCCAGTGCTTCAATGATGTGACTGCCATCATCTTCGTGGTGGCCAGCAGCAGCTACAA
CATGGTCATCCGGGAGGACAACCAGACCAACCGTCTGCAGGAGGCTCTGAACCTCTTCAAGAGCATCTGGAAC
AACAGATGGCTGCGTACCATCTCTGTGATCCTCTTCCTCAACAAGCAAGATCTGCTTGCTGAGAAGGTCCTCG
CTGGGAAATCGAAGATTGAGGACTACTTTCCAGAGTTCGCTCGCTACACCACTCCTGAGGATGCGACTCCCGA
GCCCGGAGAGGACCCACGCGTGACCCGGGCCAAGTACTTCATCCGGGATGAGTTTCTGAGAATCAGCACTGCT

AGTGGAGATGGACGTCACTACTGCTACCCTCACTTTACCTGCGCCGTGGACACTGAGAACATCCGCCGTGTCT
TCAACGACTGCCGTGACATCATCCAGCGCATGCATCTTCGCCAATACGAGCTGCTCTAAACCGGGATAAGTCA
ACTAACTTAAGCTAGCAACGGTTTCCCTCTAGCGGGATCAATTCCG*CCCCCCCCCCCTAACGTTACTGGCCGA*
*AGCCGCTTGGAATAAGGCCGGTGTGCGTTTGTCTATATGTTATTTTCCACCATATTGCCGTCTTTTGGCAATG*
*TGAGGGCCCGGAAACCTGGCCCTGTCTTCTTGACGAGCATTCCTAGGGGTCTTTCCCCTCTCGCCAAAGGAAT*
*GCAAGGTCTGTTGAATGTCGTGAAGGAAGCAGTTCCTCTGGAAGCTTCTTGAAGACAAACAACGTCTGTAGCG*
*ACCCTTTGCAGGCAGCGGAACCCCCCACCTGGCGACAGGTGCCTCTGCGGCCAAAAGCCACGTGTATAAGATA*
*CACCTGCAAAGGCGGCACAACCCCAGTGCCACGTTGTGAGTTGGATAGTTGTGGAAAGAGTCAAATGGCTCTC*
*CTCAAGCGTATTCAACAAGGGGCTGAAGGATGCCCAGAAGGTACCCCATTGTATGGGATCTGATCTGGGGCCT*
*CGGTGCACATGCTTTACATGTGTTTAGTCGAGGTTAAAAAACGTCTAGGCCCCCCGAACCACGGGGACGTGGT*
*TTTCCTTTGAAAAACACGAT*AATACCATGACCGAGTACAAGCCCACGGTGCGCCTCGCCACCCGCGACGACGT
CCCCAGGGCCGTACGCACCCTCGCCGCCGCGTTCGCCGACTACCCCGCCACGCGCCACACCGTCGATCCGGAC
CGCCACATCGAGCGGGTCACCGAGCTGCAAGAACTCTTCCTCACGCGCGTCGGGCTCGACATCGGCAAGGTGT
GGGTCGCGGACGACGGCGCCGCGGTGGCGGTCTGGACCACGCCGGAGAGCGTCGAAGCGGGGGCGGTGTTCGC
CGAGATCGGCCCGCGCATGGCCGAGTTGAGCGGTTCCCGGCTGGCCGCGCAGCAACAGATGGAAGGCCTCCTG
GCGCCGCACCGGCCCAAGGAGCCCGCGTGGTTCCTGGCCACCGTCGGCGTCTCGCCCGACCACCAGGGCAAGG
GTCTGGGCAGCGCCGTCGTGCTCCCCGGAGTGGAGGCGGCCGAGCGCGCCGGGGTGCCCGCCTTCCTGGAGAC
CTCCGCGCCCCGCAACCTCCCCTTCTACGAGCGGCTCGGCTTCACCGTCACCGCCGACGTCGAGTGCCCGAAG
GACCGCGCGACCTGGTGCATGACCCGCAAGCCCGGTGCCTGA

miRFP670 eDHFR K6-tag(69K6) Gαs (human Gαs with a C3S mutation)[S3] *IRES* Puro$^{R}$

**pCSIIpuro-miR-eDHFR(69K6)**

```
>Amino acid sequence
MVAGHASGSPAFGTASHSNCEHEEIHLAGSIQPHGALLVVSEHDHRVIQASANAAEFLNLGSVLGVPLAEIDG
DLLIKILPHLDPTAEGMPVAVRCRIGNPSTEYCGLMHRPPEGGLIIELERAGPSIDLSGTLAPALERIRTAGS
LRALCDDTVLLFQQCTGYDRVMVYRFDEQGHGLVFSECHVPGLESYFGNRYPSSTVPQMARQLYVRQRVRVLV
DVTYQPVPLEPRLSPLTGRDLDMSGCFLRSMSPCHLQFLKDMGVRATLAVSLVVGGKLWGLVVCHHYLPRFIR
FELRAICKRLAERIATRITALESLYKSGSAASSAGSMISLIAALAVDRVIGMENAMPWNLPADLAWFKRNTLN
KPVIMGRHTWESIGRPLPGRKNIILSSQPGTDKKKKKKDRVTWVKSVDEAIAACGDVPEIMVIGGGRVYEQFL
PKAQKLYLTHIDAEVEGDTHFPDYEPDDWESVFSEFHDADAQNSHSYCFEILERRLYK*-[IRES]-MTEYKP
TVRLATRDDVPRAVRTLAAAFADYPATRHTVDPDRHIERVTELQELFLTRVGLDIGKVWVADDGAAVAVWTTP
ESVEAGAVFAEIGPRMAELSGSRLAAQQQMEGLLAPHRPKEPAWFLATVGVSPDHQGKGLGSAVVLPGVEAAE
RAGVPAFLETSAPRNLPFYERLGFTVTADVECPKDRATWCMTRKPGA*

>DNA sequence
ATGGTAGCAGGTCATGCCTCTGGCAGCCCCGCATTCGGGACCGCCTCTCATTCGAATTGCGAACATGAAGAGA
TCCACCTCGCCGGCTCGATCCAGCCGCATGGCGCGCTTCTGGTCGTCAGCGAACATGATCATCGCGTCATCCA
GGCCAGCGCCAACGCCGCGGAATTTCTGAATCTCGGAAGCGTACTCGGCGTTCCGCTCGCCGAGATCGACGGC
GATCTGTTGATCAAGATCCTGCCGCATCTCGATCCCACCGCCGAAGGCATGCCGGTCGCGGTGCGCTGCCGGA
TCGGCAATCCCTCTACGGAGTACTGCGGTCTGATGCATCGGCCTCCGGAAGGCGGGCTGATCATCGAACTCGA
ACGTGCCGGCCCGTCGATCGATCTGTCAGGCACGCTGGCGCCGGCGCTGGAGCGGATCCGCACGGCGGGTTCA
CTGCGCGCGCTGTGCGATGACACCGTGCTGCTGTTTCAGCAGTGCACCGGCTACGACCGGGTGATGGTGTATC
GTTTCGATGAGCAAGGCCACGGCCTGGTATTCTCCGAGTGCCATGTGCCTGGGCTCGAATCCTATTTCGGCAA
CCGCTATCCGTCGTCGACTGTCCCGCAGATGGCGCGGCAGCTGTACGTGCGGCAGCGCGTCCGCGTGCTGGTC
GACGTCACCTATCAGCCGGTGCCGCTGGAGCCGCGGCTGTCGCCGCTGACCGGGCGCGATCTCGACATGTCGG
GCTGCTTCCTGCGCTCGATGTCGCCGTGCCATCTGCAGTTCCTGAAGGACATGGGCGTGCGCGCCACCCTGGC
GGTGTCGCTGGTGGTCGGCGGCAAGCTGTGGGGCCTGGTTGTCTGTCACCATTATCTGCCGCGCTTCATCCGT
TTCGAGCTGCGGGCGATCTGCAAACGGCTCGCCGAAAGGATCGCGACGCGGATCACCGCGCTTGAGAGCCTCT
ACAAGTCTGGGTCCGCAGCCAGTAGCGCCGGAAGTATGATCAGTCTGATTGCGGCGTTAGCGGTAGATCGCGT
TATCGGCATGGAAAACGCCATGCCGTGGAACCTGCCTGCCGATCTCGCCTGGTTTAAACGCAACACCTTAAAT
AAACCCGTGATTATGGGCCGCCATACCTGGGAATCAATCGGTCGTCCGTTGCCAGGACGCAAAAATATTATCC
TCAGCAGTCAACCGGGTACGGACAAAAAAAAGAAAAAGAAAGATCGCGTAACGTGGGTGAAGTCGGTGGATGA
AGCCATCGCGGCGTGTGGTGACGTACCAGAAATCATGGTGATTGGCGGCGGTCGCGTTTATGAACAGTTCTTG
CCAAAAGCGCAAAAACTGTATCTGACGCATATCGACGCAGAAGTGGAAGGCGACACCCATTTCCCGGATTACG
AGCCGGATGACTGGGAATCGGTATTCAGCGAATTCCACGATGCTGATGCGCAGAACTCTCACAGCTATTGCTT
TGAGATTCTGGAGCGGCGGCTGTACAAGTAAAGCGGCCGCGGCTCTAGAGGATCCGTTAACTAACTTAAGCTA
GCAACGGTTTCCCTCTAGCGGGATCAATTCCGCCCCCCCCCCCTAACGTTACTGGCCGAAGCCGCTTGGAATA
AGGCCGGTGTGCGTTTGTCTATATGTTATTTTCCACCATATTGCCGTCTTTTGGCAATGTGAGGGCCCGGAAA
CCTGGCCCTGTCTTCTTGACGAGCATTCCTAGGGGTCTTTCCCCTCTCGCCAAAGGAATGCAAGGTCTGTTGA
ATGTCGTGAAGGAAGCAGTTCCTCTGGAAGCTTCTTGAAGACAAACAACGTCTGTAGCGACCCTTTGCAGGCA
GCGGAACCCCCCACCTGGCGACAGGTGCCTCTGCGGCCAAAAGCCACGTGTATAAGATACACCTGCAAAGGCG
GCACAACCCCAGTGCCACGTTGTGAGTTGGATAGTTGTGGAAAGAGTCAAATGGCTCTCCTCAAGCGTATTCA
ACAAGGGGCTGAAGGATGCCCAGAAGGTACCCCATTGTATGGGATCTGATCTGGGGCCTCGGTGCACATGCTT
TACATGTGTTTAGTCGAGGTTAAAAAACGTCTAGGCCCCCCGAACCACGGGGACGTGGTTTTCCTTTGAAAAA
CACGATAATACCATGACCGAGTACAAGCCCACGGTGCGCCTCGCCACCCGCGACGACGTCCCCCGGGCCGTAC
GCACCCTCGCCGCCGCGTTCGCCGACTACCCCGCCACGCGCCACACCGTCGACCCGGACCGCCACATCGAGCG
GGTCACCGAGCTGCAAGAACTCTTCCTCACGCGCGTCGGGCTCGACATCGGCAAGGTGTGGGTCGCGGACGAC
GGCGCCGCGGTGGCGGTCTGGACCACGCCGGAGAGCGTCGAAGCGGGGGCGGTGTTCGCCGAGATCGGCCCGC
GCATGGCCGAGTTGAGCGGTTCCCGGCTGGCCGCGCAGCAACAGATGGAAGGCCTCCTGGCGCCGCACCGGCC
CAAGGAGCCCGCGTGGTTCCTGGCCACCGTCGGCGTCTCGCCCGACCACCAGGGCAAGGGTCTGGGCAGCGCC
GTCGTGCTCCCCGGAGTGGAGGCGGCCGAGCGCGCCGGGGTGCCCGCCTTCCTGGAGACCTCCGCGCCCCGCA
ACCTCCCCTTCTACGAGCGGCTCGGCTTCACCGTCACCGCCGACGTCGAGTGCCCGAAGGACCGCGCGACCTG
GTGCATGACCCGCAAGCCCGGTGCCTGA
```

miRFP670 eDHFR K6-tag(69K6) *IRES* Puro$^R$

#### pPBbsr-cAMPsensor

```
>Amino acid sequence
MVSKGEETTMGVIKPDMKIKLKMEGNVNGHAFVIEGEGEGKPYDGTNTINLEVKEGAPLPFSYDILTTAFAYG
NRAFTKYPDDIPNYFKQSFPEGYSWERTMTFEDKGIVKVKSDISMEEDSFIYEIHLKGENFPPNGPVMQKKTT
GWDASTERMYVRDGVLKGDVKHKLLLEGGGHHRVDFKTIYRAKKAVKLPDYHFVDHRIEILNHDKDYNKVTVY
ESAVARNSTDGMDELYKLDPVGTHEMEEELAEAVALLSQRGPDALLTVALRKPPGQRTDEELDLIFEELLHIK
AVAHLSNSVKRELAAVLLFEPHSKAGTVLFSQGDKGTSWYIIWKGSVNVVTHGKGLVTTLHEGDDFGQLALVN
DAPRAATIILREDNCHFLRVDKQDFNRIIKDVEAKTMRLEEHGKVVLVLERASQGAGPSRPPTPGRNRYTVMS
GTPEKILELLLEAMGPDSSAHDPTETFLSDFLLTHRVFMPSAQLCAALLHHFHVEPAGGSEQERSTYVCNKRQ
QILRLVSQWVALYGSMLHTDPVATSFLQKLSDLVGRDTRLSNLLREQWPERRRCHRLENGCGNASPQMKARNL
PVWLPNQDEPLPGSSCAIQVGDKVPYDICRPDHSVLTLQLPVTASVREVMAALAQEDGWTKGQVLVKVNSAGD
AIGLQPDARGVATSLGLNERLFVVNPQEAHELIPHPDQLGPTVGSAEGLDLVSAKDLAGQLTDHDWSLFNSIH
QVELIHYVLGPQHLRDVTTANLERFMRRFNELQYWVATELCLCPVPGPRAQLLRKFIKLAAHLKEQKNLNSFF
AVMFGLSNSAISRLAHTWERLPHKVRKLYSALERLLDPSWNHRVYRLALAKLSPPVIPFMPLLLKDMTFIHEG
NHTLVENLINFEKMRMMARAARMLHHCRSHNPVPLSPLRSRVSHLHEDSQVARISTCSEQSLSTRSPASTWAY
VQQLKVIDNQRELSRLSRELEPGGRMVSKGEELFTGVVPILVELDGDVNGHKFSVSGEGEGDATYGKLTLKLI
CTTGKLPVPWPTLVTTLGYGLQCFARYPDHMKQHDFFKSAMPEGYVQERTIFFKDDGNYKTRAEVKFEGDTLV
NRIELKGIDFKEDGNILGHKLEYNYNSHNVYITADKQKNGIKANFKIRHNIEDGGVQLADHYQQNTPIGDGPV
LLPDNHYLSYQSKLSKDPNEKRDHMVLLEFVTAASR*-[IRES]-MLYEDNKHHVGAAIRTKTGEIISAVHIE
AYIGRVTVCAEAIAIGSAVSNGQKDFDTIVAVRHPYSDEVDRSIRVVSPCGMCRELISDYAPDCFVLIEMNGK
LVKTTIEELIPLKYTRN*
```

```
>DNA sequence
ATGGTGAGCAAGGGCGAGGAGACCACAATGGGCGTAATCAAGCCCGACATGAAGATCAAGCTGAAGATGGAGG
GCAACGTGAATGGCCACGCCTTCGTGATCGAGGGCGAGGGCGAGGGCAAGCCCTACGACGGCACCAACACCAT
CAACCTGGAGGTGAAGGAGGGAGCCCCCCTGCCCTTCTCCTACGACATTCTGACCACCGCGTTCGCCTACGGC
AACAGGGCCTTCACCAAGTACCCCGACGACATCCCCAACTACTTCAAGCAGTCCTTCCCCGAGGGCTACTCTT
GGGAGCGCACCATGACCTTCGAGGACAAGGGCATCGTGAAGGTGAAGTCCGACATCTCCATGGAGGAGGACTC
CTTCATCTACGAGATACACCTCAAGGGCGAGAACTTCCCCCCCAACGGCCCCGTGATGCAGAAGAAGACCACC
GGCTGGGACGCCTCCACCGAGAGGATGTACGTGCGCGACGGCGTGCTGAAGGGCGACGTCAAGCACAAGCTGC
TGCTGGAGGGCGGCGGCCACCACCGCGTTGACTTCAAGACCATCTACAGGGCCAAGAAGGCGGTGAAGCTGCC
CGACTATCACTTTGTGGACCACCGCATCGAGATCCTGAACCACGACAAGGACTACAACAAGGTGACCGTTTAC
GAGAGCGCCGTGGCCCGCAACTCCACCGACGGCATGGACGAGCTGTACAAGCTCGACCCCGTGGGAACTCATG
AGATGGAGGAGGAGTTGGCCGAAGCTGTGGCCCTGCTCTCCCAGCGGGGGCCTGACGCCCTGCTCACTGTGGC
ACTTCGAAAGCCCCCAGGTCAGCGCACGGATGAAGAGCTGGACCTCATCTTTGAGGAGCTGCTGCACATCAAG
GCTGTGGCCCACCTCTCCAACTCGGTGAAGCGAGAATTAGCGGCTGTTCTGCTCTTTGAACCACACAGCAAGG
CAGGGACCGTGTTGTTCAGCCAGGGGGACAAGGGCACTTCGTGGTACATTATCTGGAAGGGATCTGTCAACGT
GGTGACCCATGGCAAGGGGCTGGTGACCACCCTGCATGAGGGAGATGATTTTGGACAGCTGGCTCTGGTGAAT
GATGCACCCCGGGCAGCCACCATCATCCTGCGAGAAGACAACTGTCATTTCCTGCGTGTGGACAAGCAGGACT
TCAACCGTATCATCAAGGATGTGGAGGCAAAGACCATGCGGCTGGAAGAACATGGCAAAGTGGTGCTGGTGCT
GGAGAGAGCCTCTCAGGGCGCCGGCCCTTCCCGACCCCCAACCCCAGGCAGGAACCGGTATACAGTGATGTCT
GGCACCCCAGAGAAGATCCTAGAGCTTCTGTTGGAGGCCATGGGACCAGATTCCAGTGCTCATGACCCAACAG
AGACATTCCTCAGCGACTTCCTCCTGACCCACAGGGTCTTCATGCCCAGCGCCCAACTCTGCGCTGCCCTTCT
GCACCACTTCCATGTGGAGCCTGCGGGTGGCAGCGAGCAGGAGCGCAGCACCTACGTCTGCAACAAGAGGCAG
CAGATCTTGCGGCTGGTCAGCCAGTGGGTGGCCCTGTATGGCTCCATGCTCCACACTGACCCTGTGGCCACCA
GCTTCCTCCAGAAACTCTCAGACCTGGTGGGCAGGGACACCCGACTCAGCAACCTGCTGAGGGAGCAGTGGCC
AGAGAGGCGGCGATGCCACAGGTTGGAGAATGGCTGTGGGAATGCATCTCCTCAGATGAAGGCCCGGAACTTG
CCTGTTTGGCTCCCCAACCAGGACGAGCCCCTTCCTGGCAGCAGCTGTGCCATCCAAGTTGGGGATAAAGTCC
CCTATGACATCTGCCGGCCAGACCACTCAGTGTTGACCCTGCAGCTGCCTGTGACAGCCTCCGTGAGAGAGGT
GATGGCAGCGTTGGCCCAGGAGGATGGCTGGACCAAGGGGCAGGTGCTGGTGAAGGTCAATTCTGCAGGTGAT
GCCATTGGCCTGCAGCCAGATGCCCGTGGTGTGGCCACATCTCTGGGGCTCAATGAGCGTCTCTTTGTTGTCA
ACCCACAGGAAGCGCATGAGCTGATCCCACACCCTGACCAGCTGGGGCCCACTGTGGGCTCTGCTGAGGGGCT
GGACCTGGTGAGTGCCAAGGACCTGGCAGGCCAGCTGACGGACCACGACTGGAGCCTCTTCAACAGTATCCAC
CAGGTGGAGCTGATCCACTATGTGCTGGGCCCCCAGCATCTGCGGGATGTCACCACCGCCAACCTGGAGCGCT
TCATGCGCCGCTTCAATGAGCTGCAGTACTGGGTGGCCACCGAGCTGTGTCTCTGCCCCGTGCCCGGCCCCCG
GGCCCAGCTGCTCAGGAAGTTCATTAAGCTGGCGGCCCACCTCAAGGAGCAGAAGAATCTCAATTCCTTCTTT
```

GCCGTCATGTTTGGCCTCAGCAACTCGGCCATCAGCCGCCTAGCCCACACCTGGGAGCGGCTGCCCCACAAAG
TCCGGAAGCTGTACTCCGCCCTCGAGAGGCTGCTGGATCCCTCATGGAACCACCGGGTATACCGACTGGCCCT
CGCCAAGCTCTCCCCTCCTGTCATCCCCTTCATGCCCCTTCTTCTCAAAGACATGACCTTCATTCATGAGGGA
AACCACACACTAGTGGAGAATCTCATCAACTTTGAGAAGATGAGAATGATGGCCAGAGCCGCGCGGATGCTGC
ACCACTGCCGAAGCCACAACCCTGTGCCTCTCTCACCACTCAGAAGCCGAGTTTCCCACCTCCACGAGGACAG
CCAGGTGGCGAGGATTTCCACATGCTCGGAGCAGTCCCTGAGCACCCGGAGTCCAGCCAGCACCTGGGCTTAT
GTCCAGCAGCTGAAGGTCATTGACAACCAGCGGGAACTCTCCCGCCTCTCCCGAGAGCTGGAGCCAGGCGGCC
GCATGGTGAGCAAGGGCGAGGAGCTGTTCACCGGGGTGGTGCCCATCCTGGTCGAGCTGGACGGCGACGTAAA
CGGCCACAAGTTCAGCGTGTCCGGCGAGGGCGAGGGCGATGCCACCTACGGCAAGCTGACCCTGAAGCTGATC
TGCACCACCGGCAAGCTGCCCGTGCCCTGGCCCACCCTCGTGACCACCCTGGGCTACGGCCTGCAGTGCTTCG
CCCGCTACCCCGACCACATGAAGCAGCACGACTTCTTCAAGTCCGCCATGCCCGAAGGCTACGTCCAGGAGCG
CACCATCTTCTTCAAGGACGACGGCAACTACAAGACCCGCGCCGAGGTGAAGTTCGAGGGCGACACCCTGGTG
AACCGCATCGAGCTGAAGGGCATCGACTTCAAGGAGGACGGCAACATCCTGGGGCACAAGCTGGAGTACAACT
ACAACAGCCACAACGTCTATATCACCGCCGACAAGCAGAAGAACGGCATCAAGGCCAACTTCAAGATCCGCCA
CAACATCGAGGACGGCGGCGTGCAGCTCGCCGACCACTACCAGCAGAACACCCCCATCGGCGACGGCCCCGTG
CTGCTGCCCGACAACCACTACCTGAGCTACCAGTCCAAGCTTAGCAAAGACCCCAACGAGAAGCGCGATCACA
TGGTCCTGCTGGAGTTCGTGACCGCCGCCTCTAGATAAGTCGACGGGCCGCGGTAACAATTGTTAACTAACTT
AAGCTAGCAACGGTTTCCCTCTAGCGGGATCAATTCCG*CCCCCCCCCCCTAACGTTACTGGCCGAAGCCGCTT*
*GGAATAAGGCCGGTGTGCGTTTGTCTATATGTTATTTTCCACCATATTGCCGTCTTTTGGCAATGTGAGGGCC*
*CGGAAACCTGGCCCTGTCTTCTTGACGAGCATTCCTAGGGGTCTTTCCCCTCTCGCCAAAGGAATGCAAGGTC*
*TGTTGAATGTCGTGAAGGAAGCAGTTCCTCTGGAAGCTTCTTGAAGACAAACAACGTCTGTAGCGACCCTTTG*
*CAGGCAGCGGAACCCCCCACCTGGCGACAGGTGCCTCTGCGGCCAAAAGCCACGTGTATAAGATACACCTGCA*
*AAGGCGGCACAACCCCAGTGCCACGTTGTGAGTTGGATAGTTGTGGAAAGAGTCAAATGGCTCTCCTCAAGCG*
*TATTCAACAAGGGGCTGAAGGATGCCCAGAAGGTACCCCATTGTATGGGATCTGATCTGGGGCCTCGGTGCAC*
*ATGCTTTACATGTGTTTAGTCGAGGTTAAAAAACGTCTAGGCCCCCCGAACCACGGGGACGTGGTTTTCCTTT*
*GAAAAACACGAT*AATACCATGGTCATGAAAACATTTAACATTTCTCAACAAGATCTAGAATTAGTAGAAGTAG
CGACAGAGAAGATTACAATGCTTTATGAGGATAATAAACATCATGTGGGAGCGGCAATTCGTACGAAAACAGG
AGAAATCATTTCGGCAGTACATATTGAAGCGTATATAGGACGAGTAACTGTTTGTGCAGAAGCCATTGCGATT
GGTAGTGCAGTTTCGAATGGACAAAAGGATTTTGACACGATTGTAGCTGTTAGACACCCTTATTCTGACGAAG
TAGATAGAAGTATTCGAGTGGTAAGTCCTTGTGGTATGTGTAGGGAGTTGATTTCAGACTATGCACCAGATTG
TTTTGTGTTAATAGAAATGAATGGCAAGTTAGTCAAAACTACGATTGAAGAACTCATTCCACTCAAATATACC
CGAAATTAA

cAMPsensor [mTFP1 the regulatory domain of human Epac (residues 149–881) mVenus][S4,S5] *IRES* Bsr$^R$

**pCSIIpuro-mNG-eDHFR(69K6)-RasGEF**

>Amino acid sequence
MVSKGEEDNMASLPATHELHIFGSINGVDFDMVGQGTGNPNDGYEELNLKSTKGDLQFSPWILVPHIGYGFHQ
YLPYPDGMSPFQAAMVDGSGYQVHRTMQFEDGASLTVNYRYTYEGSHIKGEAQVKGTGFPADGPVMTNSLTAA
DWCRSKKTYPNDKTIISTFKWSYTTGNGKRYRSTARTTYTFAKPMAANYLKNQPMYVFRKTELKHSKTELNFK
EWQKAFTDVMGMDELYKSGSAASSAGSMISLIAALAVDRVIGMENAMPWNLPADLAWFKRNTLNKPVIMGRHT
WESIGRPLPGRKNIILSSQPGTDKKKKKKDRVTWVKSVDEAIAACGDVPEIMVIGGGRVYEQFLPKAQKLYLT
HIDAEVEGDTHFPDYEPDDWESVFSEFHDADAQNSHSYCFEILERRLYKSGLRSRQGSGAGSGAGSGAGSGAG
SGAPRAQASNSTLEEITQMAEGVKAEPFENHSALEIAEQLTLLDHLVFKKIPYEEFFGQGWMKLEKNERTPYI
MKTTKHFNDISNLIASEIIRNEDINARVSAIEKWVAVADICRCLHNYNAVLEITSSMNRSAIFRLKKTWLKVS
KQTKALIDKLQKLVSSEGRFKNLREALKNCDPPCVPYLGMYLTDLAFIEEGTPNYTEDGLVNFSKMRMISHII
REIRQFQQTAYKIEHQAKVTQYLLDQSFVMDEESLYESSLRIEPKLPT*-[*IRES*]-MTEYKPTVRLATRDDV
PRAVRTLAAAFADYPATRHTVDPDRHIERVTELQELFLTRVGLDIGKVWVADDGAAVAVWTTPESVEAGAVFA
EIGPRMAELSGSRLAAQQQMEGLLAPHRPKEPAWFLATVGVSPDHQGKGLGSAVVLPGVEAAERAGVPAFLET
SAPRNLPFYERLGFTVTADVECPKDRATWCMTRKPGA*

>DNA sequence
ATGGTGAGCAAGGGCGAGGAGGATAACATGGCCTCTCTCCCAGCGACACATGAGTTACACATCTTTGGCTCCA
TCAACGGTGTGGACTTTGACATGGTGGGTCAGGGCACCGGCAATCCAAATGATGGTTATGAGGAGTTAAACCT
GAAGTCCACCAAGGGTGACCTCCAGTTCTCCCCCTGGATTCTGGTCCCTCATATCGGGTATGGCTTCCATCAG
TACCTGCCCTACCCTGACGGGATGTCGCCTTTCCAGGCCGCCATGGTAGATGGCTCCGGCTACCAAGTCCATC
GCACAATGCAGTTTGAAGATGGTGCCTCCCTTACTGTTAACTACCGCTACACCTACGAGGGAAGCCACATCAA
AGGAGAGGCCCAGGTGAAGGGGACTGGTTTCCCTGCTGACGGTCCTGTGATGACCAACTCGCTGACCGCTGCG
GACTGGTGCAGGTCGAAGAAGACTTACCCCAACGACAAAACCATCATCAGTACCTTTAAGTGGAGTTACACCA
CTGGAAATGGCAAGCGCTACCGGAGCACTGCGCGGACCACCTACACCTTTGCCAAGCCAATGGCGGCTAACTA
TCTGAAGAACCAGCCGATGTACGTGTTCCGTAAGACGGAGCTCAAGCACTCCAAGACCGAGCTCAACTTCAAG
GAGTGGCAAAAGGCCTTTACCGATGTGATGGGCATGGACGAGCTCTACAAGTCTGGGTCCGCAGCCAGTAGCG
CCGGAAGTATGATCAGTCTGATTGCGGCGTTAGCGGTAGATCGCGTTATCGGCATGGAAAACGCCATGCCGTG
GAACCTGCCTGCCGATCTCGCCTGGTTTAAACGCAACACCTTAAATAAACCCGTGATTATGGGCCGCCATACC
TGGGAATCAATCGGTCGTCCGTTGCCAGGACGCAAAAATATTATCCTCAGCAGTCAACCGGGTACGGACAAAA
AAAAGAAAAAGAAAGATCGCGTAACGTGGGTGAAGTCGGTGGATGAAGCCATCGCGGCGTGTGGTGACGTACC
AGAAATCATGGTGATTGGCGGCGGTCGCGTTTATGAACAGTTCTTGCCAAAAGCGCAAAAACTGTATCTGACG
CATATCGACGCAGAAGTGGAAGGCGACACCCATTTCCCGGATTACGAGCCGGATGACTGGGAATCGGTATTCA
GCGAATTCCACGATGCTGATGCGCAGAACTCTCACAGCTATTGCTTTGAGATTCTGGAGCGGCGGCTGTACAA
GTCCGGACTCAGATCTCGACAAGGTAGTGGTGCTGGCTCTGGTGCTGGTAGTGGCGCTGGTTCCGGTGCTGGC
TCTGGCGCGCCTCGAGCTCAAGCTTCGAATTCTACGCTGGAGGAGATCACGCAGATGGCTGAAGGCGTGAAGG
CTGAGCCCTTTGAAAACCACTCAGCCCTGGAGATCGCGGAGCAGCTGACCCTGCTAGATCACCTCGTCTTCAA
GAAGATTCCTTATGAGGAGTTCTTCGGACAAGGATGGATGAAACTGGAAAAGAATGAAAGGACCCCTTATATC
ATGAAAACCACTAAGCACTTCAATGACATCAGTAACTTGATTGCTTCAGAAATCATCCGCAATGAGGACATCA
ACGCCAGGGTGAGCGCCATCGAGAAGTGGGTGGCCGTAGCTGACATATGCCGCTGCCTCCACAACTACAATGC
CGTACTGGAGATCACCTCGTCCATGAACCGCAGTGCAATCTTCCGGCTCAAAAAGACGTGGCTCAAAGTCTCT
AAGCAGACTAAAGCTTTGATTGATAAGCTCCAAAAGCTTGTGTCATCTGAGGGCAGATTTAAGAATCTCAGAG
AAGCTCTGAAAAATTGTGACCCACCCTGTGTCCCTTACCTGGGGATGTACCTCACCGACCTGGCCTTCATCGA
GGAGGGGACGCCCAATTACACGGAAGACGGCCTGGTCAACTTCTCCAAGATGAGGATGATATCCCATATTATC
CGAGAGATTCGCCAGTTTCAACAAACTGCCTACAAAATAGAGCACCAAGCAAAGGTAACGCAATATTTACTGG
ACCAATCTTTTGTAATGGATGAAGAAAGCCTCTACGAGTCTTCTCTCCGAATAGAACCAAAACTCCCCACCTA
GGTCAACTAACTTAAGCTAGCAACGGTTTCCCTCTAGCGGGATCAATTCCG*CCCCCCCCCCCTAACGTTACTG*
*GCCGAAGCCGCTTGGAATAAGGCCGGTGTGCGTTTGTCTATATGTTATTTTCCACCATATTGCCGTCTTTTGG*
*CAATGTGAGGGCCCGGAAACCTGGCCCTGTCTTCTTGACGAGCATTCCTAGGGGTCTTTCCCCTCTCGCCAAA*
*GGAATGCAAGGTCTGTTGAATGTCGTGAAGGAAGCAGTTCCTCTGGAAGCTTCTTGAAGACAAACAACGTCTG*
*TAGCGACCCTTTGCAGGCAGCGGAACCCCCCACCTGGCGACAGGTGCCTCTGCGGCCAAAAGCCACGTGTATA*
*AGATACACCTGCAAAGGCGGCACAACCCCAGTGCCACGTTGTGAGTTGGATAGTTGTGGAAAGAGTCAAATGG*
*CTCTCCTCAAGCGTATTCAACAAGGGGCTGAAGGATGCCCAGAAGGTACCCCATTGTATGGGATCTGATCTGG*
*GGCCTCGGTGCACATGCTTTACATGTGTTTAGTCGAGGTTAAAAAACGTCTAGGCCCCCCGAACCACGGGGAC*
*GTGGTTTTCCTTTGAAAAACACGAT*AATACCATGACCGAGTACAAGCCCACGGTGCGCCTCGCCACCCGCGAC
GACGTCCCCCGGGCCGTACGCACCCTCGCCGCCGCGTTCGCCGACTACCCCGCCACGCGCCACACCGTCGACC

CGGACCGCCACATCGAGCGGGTCACCGAGCTGCAAGAACTCTTCCTCACGCGCGTCGGGCTCGACATCGGCAA
GGTGTGGGTCGCGGACGACGGCGCCGCGGTGGCGGTCTGGACCACGCCGGAGAGCGTCGAAGCGGGGGCGGTG
TTCGCCGAGATCGGCCCGCGCATGGCCGAGTTGAGCGGTTCCCGGCTGGCCGCGCAGCAACAGATGGAAGGCC
TCCTGGCGCCGCACCGGCCCAAGGAGCCCGCGTGGTTCCTGGCCACCGTCGGCGTCTCGCCCGACCACCAGGG
CAAGGGTCTGGGCAGCGCCGTCGTGCTCCCCGGAGTGGAGGCGGCCGAGCGCGCCGGGGTGCCCGCCTTCCTG
GAGACCTCCGCGCCCCGCAACCTCCCCTTCTACGAGCGGCTCGGCTTCACCGTCACCGCCGACGTCGAGTGCC
CGAAGGACCGCGCGACCTGGTGCATGACCCGCAAGCCCGGTGCCTGA

mNeonGreen eDHFR K6-tag(69K6) RasGEF [the Cdc25 domain of human RasGRF1 (residues 1018–1273)][S6] *IRES* Puro$^R$

### pCSIIpuro-mNG-eDHFR(69K6)

>Amino acid sequence
MVSKGEEDNMASLPATHELHIFGSINGVDFDMVGQGTGNPNDGYEELNLKSTKGDLQFSPWILVPHIGYGFHQ
YLPYPDGMSPFQAAMVDGSGYQVHRTMQFEDGASLTVNYRYTYEGSHIKGEAQVKGTGFPADGPVMTNSLTAA
DWCRSKKTYPNDKTIISTFKWSYTTGNGKRYRSTARTTYTFAKPMAANYLKNQPMYVFRKTELKHSKTELNFK
EWQKAFTDVMGMDELYKSGSAASSAGSMISLIAALAVDRVIGMENAMPWNLPADLAWFKRNTLNKPVIMGRHT
WESIGRPLPGRKNIILSSQPGTDKKKKKKDRVTWVKSVDEAIAACGDVPEIMVIGGGRVYEQFLPKAQKLYLT
HIDAEVEGDTHFPDYEPDDWESVFSEFHDADAQNSHSYCFEILERRLYK*-[*IRES*]-MTEYKPTVRLATRDD
VPRAVRTLAAAFADYPATRHTVDPDRHIERVTELQELFLTRVGLDIGKVWVADDGAAVAVWTTPESVEAGAVF
AEIGPRMAELSGSRLAAQQQMEGLLAPHRPKEPAWFLATVGVSPDHQGKGLGSAVVLPGVEAAERAGVPAFLE
TSAPRNLPFYERLGFTVTADVECPKDRATWCMTRKPGA*

>DNA sequence
ATGGTGAGCAAGGGCGAGGAGGATAACATGGCCTCTCTCCCAGCGACACATGAGTTACACATCTTTGGCTCCA
TCAACGGTGTGGACTTTGACATGGTGGGTCAGGGCACCGGCAATCCAAATGATGGTTATGAGGAGTTAAACCT
GAAGTCCACCAAGGGTGACCTCCAGTTCTCCCCCTGGATTCTGGTCCCTCATATCGGGTATGGCTTCCATCAG
TACCTGCCCTACCCTGACGGGATGTCGCCTTTCCAGGCCGCCATGGTAGATGGCTCCGGCTACCAAGTCCATC
GCACAATGCAGTTTGAAGATGGTGCCTCCCTTACTGTTAACTACCGCTACACCTACGAGGGAAGCCACATCAA
AGGAGAGGCCCAGGTGAAGGGGACTGGTTTCCCTGCTGACGGTCCTGTGATGACCAACTCGCTGACCGCTGCG
GACTGGTGCAGGTCGAAGAAGACTTACCCCAACGACAAAACCATCATCAGTACCTTTAAGTGGAGTTACACCA
CTGGAAATGGCAAGCGCTACCGGAGCACTGCGCGGACCACCTACACCTTTGCCAAGCCAATGGCGGCTAACTA
TCTGAAGAACCAGCCGATGTACGTGTTCCGTAAGACGGAGCTCAAGCACTCCAAGACCGAGCTCAACTTCAAG
GAGTGGCAAAAGGCCTTTACCGATGTGATGGGCATGGACGAGCTCTACAAGTCTGGGTCCGCAGCCAGTAGCG
CCGGAAGTATGATCAGTCTGATTGCGGCGTTAGCGGTAGATCGCGTTATCGGCATGGAAAACGCCATGCCGTG
GAACCTGCCTGCCGATCTCGCCTGGTTTAAACGCAACACCTTAAATAAACCCGTGATTATGGGCCGCCATACC
TGGGAATCAATCGGTCGTCCGTTGCCAGGACGCAAAAATATTATCCTCAGCAGTCAACCGGGTACGGACAAAA
AAAAGAAAAAGAAAGATCGCGTAACGTGGGTGAAGTCGGTGGATGAAGCCATCGCGGCGTGTGGTGACGTACC
AGAAATCATGGTGATTGGCGGCGGTCGCGTTTATGAACAGTTCTTGCCAAAAGCGCAAAAACTGTATCTGACG
CATATCGACGCAGAAGTGGAAGGCGACACCCATTTCCCGGATTACGAGCCGGATGACTGGGAATCGGTATTCA
GCGAATTCCACGATGCTGATGCGCAGAACTCTCACAGCTATTGCTTTGAGATTCTGGAGCGGCGGCTGTACAA
GTAAAGCGGCCGCGGCTCTAGAGGATCCGTTAACTAACTTAAGCTAGCAACGGTTTCCCTCTAGCGGGATCAA
TTCCG*CCCCCCCCCCCTAACGTTACTGGCCGAAGCCGCTTGGAATAAGGCCGGTGTGCGTTTGTCTATATGTT*
*ATTTTCCACCATATTGCCGTCTTTTGGCAATGTGAGGGCCCGGAAACCTGGCCCTGTCTTCTTGACGAGCATT*
*CCTAGGGGTCTTTCCCCTCTCGCCAAAGGAATGCAAGGTCTGTTGAATGTCGTGAAGGAAGCAGTTCCTCTGG*
*AAGCTTCTTGAAGACAAACAACGTCTGTAGCGACCCTTTGCAGGCAGCGGAACCCCCCACCTGGCGACAGGTG*
*CCTCTGCGGCCAAAAGCCACGTGTATAAGATACACCTGCAAAGGCGGCACAACCCCAGTGCCACGTTGTGAGT*
*TGGATAGTTGTGGAAAGAGTCAAATGGCTCTCCTCAAGCGTATTCAACAAGGGGCTGAAGGATGCCCAGAAGG*
*TACCCCATTGTATGGATCTGATCTGGGGCCTCGGTGCACATGCTTTACATGTGTTTAGTCGAGGTTAAAAAA*
*CGTCTAGGCCCCCCGAACCACGGGGACGTGGTTTTCCTTTGAAAAACACGAT*AATACCATGACCGAGTACAAG
CCCACGGTGCGCCTCGCCACCCGCGACGACGTCCCCCGGGCCGTACGCACCCTCGCCGCCGCGTTCGCCGACT
ACCCCGCCACGCGCCACACCGTCGACCCGGACCGCCACATCGAGCGGGTCACCGAGCTGCAAGAACTCTTCCT
CACGCGCGTCGGGCTCGACATCGGCAAGGTGTGGGTCGCGGACGACGGCGCCGCGGTGGCGGTCTGGACCACG
CCGGAGAGCGTCGAAGCGGGGGCGGTGTTCGCCGAGATCGGCCCGCGCATGGCCGAGTTGAGCGGTTCCCGGC
TGGCCGCGCAGCAACAGATGGAAGGCCTCCTGGCGCCGCACCGGCCCAAGGAGCCCGCGTGGTTCCTGGCCAC
CGTCGGCGTCTCGCCCGACCACCAGGGCAAGGGTCTGGGCAGCGCCGTCGTGCTCCCCGGAGTGGAGGCGGCC
GAGCGCGCCGGGGTGCCCGCCTTCCTGGAGACCTCCGCGCCCCGCAACCTCCCCTTCTACGAGCGGCTCGGCT
TCACCGTCACCGCCGACGTCGAGTGCCCGAAGGACCGCGCGACCTGGTGCATGACCCGCAAGCCCGGTGCCTG
A

mNeonGreen eDHFR K6-tag(69K6) *IRES* Puro$^R$

**pPBbsr-MEK-P2A-mCherry-ERK**

>Amino acid sequence
MDYKDDDDKARLEMPKKKPTPIQLNPNPEGTAVNGTPTAETNLEALQKKLEELELDEQQRKRLEAFLTQKQKV
GELKDDDFEKVSELGAGNGGVVFKVSHKPTSLIMARKLIHLEIKPAIRNQIIRELQVLHECNSPYIVGFYGAF
YSDGEISICMEHMDGGSLDQVLKKAGKIPEKILGKVSIAVIKGLTYLREKHKIMHRDVKPSNILVNSRGEIKL
CDFGVSGQLIDSMANSFVGTRSYMSPERLQGTHYSVQSDIWSMGLSLVEMAIGRYPIPPPDAKELELIFGCSV
ERDPASSELAPRPRPPGRPISSYGPDSRPPMAIFELLDYIVNEPPPKLPSGVFGAEFQDFVNKCLVKNPAERA
DLKQLMVHSFIKQSELEEVDFAGWLCSTMGLKQPSTPTHAAGVGGRGTSGSGATNFSLLKQAGDVEENPGPQL
IKGAMVSKGEEDNMAIIKEFMRFKVHMEGSVNGHEFEIEGEGEGRPYEGTQTAKLKVTKGGPLPFAWDILSPQ
FMYGSKAYVKHPADIPDYLKLSFPEGFKWERVMNFEDGGVVTVTQDSSLQDGEFIYKVKLRGTNFPSDGPVMQ
KKTMGWEASSERMYPEDGALKGEIKQRLKLKDGGHYDAEVKTTYKAKKPVQLPGAYNVNIKLDITSHNEDYTI
VEQYDRAEGRHSTGGMDELYLEMAAAGAASNPGGGPEMVRGQAFDVGPRYINLAYIGEGAYGMVCSAHDNVNK
VRVAIRKISPFEHQTYCQRTLREIKILLRFKHENIIGINDIIRAPTIEQMKDVYIVQDLMETDLYKLLKTQHL
SNDHICYFLYQILRGLKYIHSANVLHRDLKPSNLLLNTTCDLKICDFGLARVADPDHDHTGFLTEYVATRWYR
APEIMLNSKGYTKSIDIWSVGCILAEMLSNRPIFPGKHYLDQLNHILGILGSPSQEDLNCIINLKARNYLLSL
PHKNKVPWNRLFPNADPKALDLLDKMLTFNPHKRIEVEAALAHPYLEQYYDPSDEPVAEAPFKFEMELDDLPK
ETLKELIFEETARFQPGY*-[*IRES*]-MVMKTFNISQQDLELVEVATEKITMLYEDNKHHVGAAIRTKTGEII
SAVHIEAYIGRVTVCAEAIAIGSAVSNGQKDFDTIVAVRHPYSDEVDRSIRVVSPCGMCRELISDYAPDCFVL
IEMNGKLVKTTIEELIPLKYTRN*

>DNA sequence
ATGGACTACAAAGACGATGACGATAAAGCAAGGCTCGAGATGCCTAAAAAGAAGCCTACGCCCATACAGCTGA
ATCCCAACCCCGAAGGGACTGCTGTGAACGGGACCCCTACAGCCGAGACAAACCTTGAAGCTCTGCAGAAAAA
GTTGGAAGAGCTTGAGCTGGATGAGCAGCAGAGGAAGCGTCTGGAGGCTTTTCTCACCCAGAAGCAGAAAGTT
GGGGAACTGAAGGATGACGACTTTGAAAAAGTTTCAGAGCTTGGAGCAGGCAACGGAGGAGTGGTGTTTAAGG
TGTCCCACAAGCCAACCAGCTTGATTATGGCCAGGAAGTTGATTCATCTGGAGATTAAGCCTGCAATCCGAAA
CCAGATTATCCGAGAGTTGCAGGTTCTGCATGAATGTAACTCCCCATACATTGTGGGGTTCTATGGGGCCTTC
TACAGTGATGGAGAGATCAGCATTTGCATGGAACACATGGATGGAGGCTCCCTTGATCAGGTTCTGAAGAAAG
CTGGCAAAATCCCAGAAAAGATTTTGGGAAAAGTCAGCATTGCAGTGATAAAAGGTCTAACCTACCTGAGAGA
AAAGCATAAGATAATGCACAGAGATGTGAAACCTTCTAACATCCTGGTCAACTCTAGAGGAGAGATAAAACTC
TGCGACTTTGGGGTCAGCGGGCAACTCATAGACTCCATGGCAAATTCCTTTGTTGGGACAAGATCCTATATGT
CACCGGAGCGACTACAGGGCACTCATTATTCTGTGCAATCAGACATCTGGAGCATGGGGCTGTCGCTGGTGGA
AATGGCCATTGGAAGGTATCCCATTCCACCCCCTGATGCCAAAGAGCTGGAACTTATCTTTGGGTGTTCTGTA
GAAAGGGATCCAGCGTCTTCTGAACTGGCACCTCGCCCCCGGCCACCCGGACGTCCAATAAGCTCATACGGTC
CTGATAGTCGACCACCCATGGCTATTTTTGAACTTCTGGATTATATCGTGAACGAGCCGCCTCCAAAATTGCC
CAGTGGAGTATTTGGAGCTGAGTTCCAGGACTTTGTGAATAAATGTCTTGTGAAGAATCCGGCAGAGAGAGCA
GACCTTAAACAGCTAATGGTTCACAGCTTCATTAAGCAGTCAGAGTTGGAGGAAGTGGATTTTGCTGGATGGC
TCTGTTCCACTATGGGCCTTAAGCAGCCCAGTACCCCAACCCATGCCGCCGGAGTGGGCGGCCGCGGCACTAG
TGGAAGCGGAGCTACTAACTTCAGCCTGCTGAAGCAGGCTGGAGACGTGGAGGAGAACCCTGGACCTCAATTA
ATTAAGGGCGCAATGGTGAGCAAGGGCGAGGAGGATAACATGGCCATCATCAAGGAGTTCATGCGCTTCAAGG
TGCACATGGAGGGCTCCGTGAACGGCCACGAGTTCGAGATCGAGGGCGAGGGCGAGGGCCGCCCCTACGAGGG
CACCCAGACCGCCAAGCTGAAGGTGACCAAGGGTGGCCCCCTGCCCTTCGCCTGGGACATCCTGTCCCCTCAG
TTCATGTACGGCTCCAAGGCCTACGTGAAGCACCCCGCCGACATCCCCGACTACTTGAAGCTGTCCTTCCCCG
AGGGCTTCAAGTGGGAGCGCGTGATGAACTTCGAGGACGGCGGCGTGGTGACCGTGACCCAGGACTCCTCCCT
GCAGGACGGCGAGTTCATCTACAAGGTGAAGCTGCGCGGCACCAACTTCCCCTCCGACGGCCCCGTAATGCAG
AAGAAGACCATGGGCTGGGAGGCCTCCTCCGAGCGGATGTACCCCGAGGACGGCGCCCTGAAGGGCGAGATCA
AGCAGAGGCTGAAGCTGAAGGACGGCGGCCACTACGACGCTGAGGTCAAGACCACCTACAAGGCCAAGAAGCC
CGTGCAGCTGCCCGGCGCCTACAACGTCAACATCAAGTTGGACATCACCTCCCACAACGAGGACTACACCATC
GTGGAACAGTACGACCGCGCCGAGGGCCGCCACTCCACCGGCGGCATGGACGAGCTGTACCTCGAGATGGCAG
CGGCAGGAGCTGCGTCTAACCCCGGCGGGGGTCCGGAGATGGTGCGGGGCCAGGCGTTCGACGTAGGCCCTCG
ATACATCAATCTGGCTTATATCGGCGAGGGAGCGTACGGCATGGTGTGTTCTGCCCATGACAATGTTAACAAA
GTTCGAGTTGCTATCAGGAAAATCAGCCCATTTGAGCATCAGACATACTGCCAGCGAACATTGCGGGAGATCA
AAATCTTGCTACGTTTTAAACATGAAAACATCATTGGGATAAACGACATTATTCGCGCTCCAACCATTGAGCA
GATGAAAGATGTGTACATTGTGCAGGACCTCATGGAGACAGACCTCTATAAGCTCCTGAAGACTCAGCATCTT
AGCAATGACCATATCTGCTATTTCTTGTACCAGATTCTGAGAGGATTAAAGTACATCCATTCAGCCAATGTTC
TACATCGTGATCTTAAGCCTTCAAATTTGCTGCTTAACACTACCTGTGATCTCAAGATCTGTGATTTTGGATT

GGCTCGTGTTGCAGACCCAGATCATGATCACACTGGCTTTCTCACAGAATATGTAGCCACTCGCTGGTACAGA
GCTCCTGAGATCATGCTGAATTCCAAGGGCTATACCAAATCAATTGACATCTGGTCTGTTGGCTGCATTCTTG
CTGAGATGCTTTCTAATAGACCCATATTTCCTGGGAAACATTATCTTGACCAGCTTAATCACATACTTGGTAT
TCTTGGATCTCCATCTCAAGAGGACCTAAACTGTATAATCAATTTAAAAGCTAGGAATTACTTGCTTTCCCTT
CCTCACAAAAATAAGGTGCCATGGAACAGACTTTTCCCCAATGCAGATCCCAAAGCTCTAGACTTACTGGACA
AGATGCTGACTTTCAACCCCCATAAAAGAATTGAAGTAGAGGCAGCTTTGGCTCATCCTTATCTGGAGCAGTA
TTATGACCCAAGTGATGAGCCTGTAGCTGAAGCTCCCTTTAAATTTGAAATGGAGCTTGATGATTTGCCCAAG
GAGACTCTTAAGGAGCTAATTTTTGAAGAAACCGCTAGATTCCAGCCAGGGTACTAATCGCGCCTCTAGAGGA
TCCGTTAACTAACTTAAGCTAGCGTCGACGGGCCGCGGTAACAATTGTTAACTAACTTAAGCTAGCAACGGTT
TCCCTCTAGCGGGATCAATTCCG*CCCCCCCCCCCTAACGTTACTGGCCGAAGCCGCTTGGAATAAGGCCGGTG*
*TGCGTTTGTCTATATGTTATTTTCCACCATATTGCCGTCTTTTGGCAATGTGAGGGCCCGGAAACCTGGCCCT*
*GTCTTCTTGACGAGCATTCCTAGGGGTCTTTCCCCTCTCGCCAAAGGAATGCAAGGTCTGTTGAATGTCGTGA*
*AGGAAGCAGTTCCTCTGGAAGCTTCTTGAAGACAAACAACGTCTGTAGCGACCCTTTGCAGGCAGCGGAACCC*
*CCCACCTGGCGACAGGTGCCTCTGCGGCCAAAAGCCACGTGTATAAGATACACCTGCAAAGGCGGCACAACCC*
*CAGTGCCACGTTGTGAGTTGGATAGTTGTGGAAAGAGTCAAATGGCTCTCCTCAAGCGTATTCAACAAGGGGC*
*TGAAGGATGCCCAGAAGGTACCCCATTGTATGGGATCTGATCTGGGGCCTCGGTGCACATGCTTTACATGTGT*
*TTAGTCGAGGTTAAAAAACGTCTAGGCCCCCCGAACCACGGGGACGTGGTTTTCCTTTGAAAAACACGAT*AAT
ACCATGGTCATGAAAACATTTAACATTTCTCAACAAGATCTAGAATTAGTAGAAGTAGCGACAGAGAAGATTA
CAATGCTTTATGAGGATAATAAACATCATGTGGGAGCGGCAATTCGTACGAAAACAGGAGAAATCATTTCGGC
AGTACATATTGAAGCGTATATAGGACGAGTAACTGTTTGTGCAGAAGCCATTGCGATTGGTAGTGCAGTTTCG
AATGGACAAAAGGATTTTGACACGATTGTAGCTGTTAGACACCCTTATTCTGACGAAGTAGATAGAAGTATTC
GAGTGGTAAGTCCTTGTGGTATGTGTAGGGAGTTGATTTCAGACTATGCACCAGATTGTTTTGTGTTAATAGA
AATGAATGGCAAGTTAGTCAAAACTACGATTGAAGAACTCATTCCACTCAAATATACCCGAAATTAA

FLAG-tag MEK1(*Xenopus laevis*) 2A self-cleaving peptide (P2A)[S32] mCherry
ERK2(K57R)(*Xenopus laevis*)[S33] *IRES* Bsr$^R$

**pPBpuro-mNG-eDHFR(69K6)-p85$_{iSH2}$**

```
>Amino acid sequence
MVSKGEEDNMASLPATHELHIFGSINGVDFDMVGQGTGNPNDGYEELNLKSTKGDLQFSPWILVPHIGYGFHQ
YLPYPDGMSPFQAAMVDGSGYQVHRTMQFEDGASLTVNYRYTYEGSHIKGEAQVKGTGFPADGPVMTNSLTAA
DWCRSKKTYPNDKTIISTFKWSYTTGNGKRYRSTARTTYTFAKPMAANYLKNQPMYVFRKTELKHSKTELNFK
EWQKAFTDVMGMDELYKSGSAASSAGSMISLIAALAVDRVIGMENAMPWNLPADLAWFKRNTLNKPVIMGRHT
WESIGRPLPGRKNIILSSQPGTDKKKKKKDRVTWVKSVDEAIAACGDVPEIMVIGGGRVYEQFLPKAQKLYLT
HIDAEVEGDTHFPDYEPDDWESVFSEFHDADAQNSHSYCFEILERRLYKSGLRSRQGSGAGSGAGSGAGSGAG
SGAPRASKYQQDQVVKEDSVEAVGAQLKVYHQQYQDKSREYDQLYEEYTRTSQELQMKRTAIEAFNETIKIFE
EQGQTQEKCSKEYLERFRREGNEKEMQRILLNSERLKSRIAEIHESRTKLEQDLRAQASDNREIDKRMNSLKP
DLMQLRKIRDQYLVWLTQKGARQRKINEWLGIKNETEDQYSLMEDEDALPHHEERT*-[IRES]-MTEYKPTV
RLATRDDVPRAVRTLAAAFADYPATRHTVDPDRHIERVTELQELFLTRVGLDIGKVWVADDGAAVAVWTTPES
VEAGAVFAEIGPRMAELSGSRLAAQQQMEGLLAPHRPKEPAWFLATVGVSPDHQGKGLGSAVVLPGVEAAERA
GVPAFLETSAPRNLPFYERLGFTVTADVECPKDRATWCMTRKPGA*

>DNA sequence
ATGGTGAGCAAGGGCGAGGAGGATAACATGGCCTCTCTCCCAGCGACACATGAGTTACACATCTTTGGCTCCA
TCAACGGTGTGGACTTTGACATGGTGGGTCAGGGCACCGGCAATCCAAATGATGGTTATGAGGAGTTAAACCT
GAAGTCCACCAAGGGTGACCTCCAGTTCTCCCCCTGGATTCTGGTCCCTCATATCGGGTATGGCTTCCATCAG
TACCTGCCCTACCCTGACGGGATGTCGCCTTTCCAGGCCGCCATGGTAGATGGCTCCGGCTACCAAGTCCATC
GCACAATGCAGTTTGAAGATGGTGCCTCCCTTACTGTTAACTACCGCTACACCTACGAGGGAAGCCACATCAA
AGGAGAGGCCCAGGTGAAGGGGACTGGTTTCCCTGCTGACGGTCCTGTGATGACCAACTCGCTGACCGCTGCG
GACTGGTGCAGGTCGAAGAAGACTTACCCCAACGACAAAACCATCATCAGTACCTTTAAGTGGAGTTACACCA
CTGGAAATGGCAAGCGCTACCGGAGCACTGCGCGGACCACCTACACCTTTGCCAAGCCAATGGCGGCTAACTA
TCTGAAGAACCAGCCGATGTACGTGTTCCGTAAGACGGAGCTCAAGCACTCCAAGACCGAGCTCAACTTCAAG
GAGTGGCAAAAGGCCTTTACCGATGTGATGGGCATGGACGAGCTCTACAAGTCTGGGTCCGCAGCCAGTAGCG
CCGGAAGTATGATCAGTCTGATTGCGGCGTTAGCGGTAGATCGCGTTATCGGCATGGAAAACGCCATGCCGTG
GAACCTGCCTGCCGATCTCGCCTGGTTTAAACGCAACACCTTAAATAAACCCGTGATTATGGGCCGCCATACC
TGGGAATCAATCGGTCGTCCGTTGCCAGGACGCAAAAATATTATCCTCAGCAGTCAACCGGGTACGGACAAAA
AAAAGAAAAAGAAAGATCGCGTAACGTGGGTGAAGTCGGTGGATGAAGCCATCGCGGCGTGTGGTGACGTACC
AGAAATCATGGTGATTGGCGGCGGTCGCGTTTATGAACAGTTCTTGCCAAAAGCGCAAAAACTGTATCTGACG
CATATCGACGCAGAAGTGGAAGGCGACACCCATTTCCCGGATTACGAGCCGGATGACTGGGAATCGGTATTCA
GCGAATTCCACGATGCTGATGCGCAGAACTCTCACAGCTATTGCTTTGAGATTCTGGAGCGGCGGCTGTACAA
GTCCGGACTCAGATCTCGACAAGGTAGTGGTGCTGGCTCTGGTGCTGGTAGTGGCGCTGGTTCCGGTGCTGGC
TCTGGCGCGCCTCGAGCATCCAAGTACCAACAAGACCAGGTGGTGAAGGAGGACAGCGTAGAGGCTGTGGGCG
CCCAGCTCAAGGTCTACCACCAGCAGTACCAGGACAAGAGCCGCGAATATGACCAGCTGTATGAAGAATACAC
ACGGACCTCCCAGGAGCTGCAGATGAAGCGCACAGCCATAGAGGCCTTCAACGAGACCATCAAGATCTTCGAA
GAGCAGGGCCAGACACAGGAGAAGTGCAGCAAGGAGTATTTGGAGCGCTTCCGGCGAGAGGGAAATGAGAAGG
AGATGCAGAGGATCCTGCTGAACTCCGAGCGACTCAAGTCTCGCATCGCGGAGATACACGAAAGCCGCACGAA
GTTGGAGCAGGATCTGCGGGCGCAGGCCTCCGACAACCGTGAGATCGACAAGCGCATGAACAGCCTCAAACCT
GACCTCATGCAGCTGCGCAAGATCAGGGACCAGTACCTCGTGTGGCTCACCCAGAAAGGTGCCCGACAGAGGA
AGATCAACGAATGGCTGGGAATCAAGAACGAGACTGAGGACCAGTATTCACTGATGGAGGATGAGGACGCCCT
CCCCCACCACGAGGAGCGCACGTGAGAATTCTGCAGTCGACGGTACCGCGGGCCCGGGATAAGTCAACTAACT
TAAGCTAGCAACGGTTTCCCTCTAGCGGGATCAATTCCGCCCCCCCCCCTAACGTTACTGGCCGAAGCCGCT
TGGAATAAGGCCGGTGTGCGTTTGTCTATATGTTATTTTCCACCATATTGCCGTCTTTTGGCAATGTGAGGGC
CCGGAAACCTGGCCCTGTCTTCTTGACGAGCATTCCTAGGGGTCTTTCCCCTCTCGCCAAAGGAATGCAAGGT
CTGTTGAATGTCGTGAAGGAAGCAGTTCCTCTGGAAGCTTCTTGAAGACAAACAACGTCTGTAGCGACCCTTT
GCAGGCAGCGGAACCCCCCACCTGGCGACAGGTGCCTCTGCGGCCAAAAGCCACGTGTATAAGATACACCTGC
AAAGGCGGCACAACCCCAGTGCCACGTTGTGAGTTGGATAGTTGTGGAAAGAGTCAAATGGCTCTCCTCAAGC
GTATTCAACAAGGGGCTGAAGGATGCCCAGAAGGTACCCCATTGTATGGGATCTGATCTGGGGCCTCGGTGCA
CATGCTTTACATGTGTTTAGTCGAGGTTAAAAAACGTCTAGGCCCCCCGAACCACGGGGACGTGGTTTTCCTT
TGAAAAACACGATAATACCATGACCGAGTACAAGCCCACGGTGCGCCTCGCCACCCGCGACGACGTCCCCAGG
GCCGTACGCACCCTCGCCGCCGCGTTCGCCGACTACCCCGCCACGCGCCACACCGTCGATCCGGACCGCCACA
TCGAGCGGGTCACCGAGCTGCAAGAACTCTTCCTCACGCGCGTCGGGCTCGACATCGGCAAGGTGTGGGTCGC
GGACGACGGCGCCGCGGTGGCGGTCTGGACCACGCCGGAGAGCGTCGAAGCGGGGGCGGTGTTCGCCGAGATC
GGCCCGCGCATGGCCGAGTTGAGCGGTTCCCGGCTGGCCGCGCAGCAACAGATGGAAGGCCTCCTGGCGCCGC
```

ACCGGCCCAAGGAGCCCGCGTGGTTCCTGGCCACCGTCGGCGTCTCGCCCGACCACCAGGGCAAGGGTCTGGG
CAGCGCCGTCGTGCTCCCCGGAGTGGAGGCGGCCGAGCGCGCCGGGGTGCCCGCCTTCCTGGAGACCTCCGCG
CCCCGCAACCTCCCCTTCTACGAGCGGCTCGGCTTCACCGTCACCGCCGACGTCGAGTGCCCGAAGGACCGCG
CGACCTGGTGCATGACCCGCAAGCCCGGTGCCTGA

mNeonGreen eDHFR K6-tag(69K6) p85$_{iSH2}$ [the iSH2 domain of human PI3K/p85α (residues 617–724)][S7,S8] *IRES* Puro$^R$

**pPBbsr-mCherry-PH$_{Akt}$**

>Amino acid sequence
MVSKGEEDNMAIIKEFMRFKVHMEGSVNGHEFEIEGEGEGRPYEGTQTAKLKVTKGGPLPFAWDILSPQFMYG
SKAYVKHPADIPDYLKLSFPEGFKWERVMNFEDGGVVTVTQDSSLQDGEFIYKVKLRGTNFPSDGPVMQKKTM
GWEASSERMYPEDGALKGEIKQRLKLKDGGHYDAEVKTTYKAKKPVQLPGAYNVNIKLDITSHNEDYTIVEQY
ERAEGRHSTGGMDELYKSGLRSRAQASNSAVDGTAGPGSMSDVAIVKEGWLHKRGEYIKTWRPRYFLLKNDGT
FIGYKERPQDVDQREAPLNNFSVAQCQLMKTERPRPNTFIIRCLQWTTVIERTFHVETPEEREEWTTAIQTVA
DGLKKQEEEEMDFRSGSPSDNSGAEEMEVSLAKPKHRVTMN*-[*IRES*]-MVMKTFNISQQDLELVEVATEKI
TMLYEDNKHHVGAAIRTKTGEIISAVHIEAYIGRVTVCAEAIAIGSAVSNGQKDFDTIVAVRHPYSDEVDRSI
RVVSPCGMCRELISDYAPDCFVLIEMNGKLVKTTIEELIPLKYTRN*

>DNA sequence
ATGGTGAGCAAGGGCGAGGAGGATAACATGGCCATCATCAAGGAGTTCATGCGCTTCAAGGTGCACATGGAGG
GCTCCGTGAACGGCCACGAGTTCGAGATCGAGGGCGAGGGCGAGGGCCGCCCCTACGAGGGCACCCAGACCGC
CAAGCTGAAGGTGACCAAGGGTGGCCCCCTGCCCTTCGCCTGGGACATCCTGTCCCCTCAGTTCATGTACGGC
TCCAAGGCCTACGTGAAGCACCCCGCCGACATCCCCGACTACTTGAAGCTGTCCTTCCCCGAGGGCTTCAAGT
GGGAGCGCGTGATGAACTTCGAGGACGGCGGCGTGGTGACCGTGACCCAGGACTCCTCCCTGCAGGACGGCGA
GTTCATCTACAAGGTGAAGCTGCGCGGCACCAACTTCCCCTCCGACGGCCCCGTAATGCAGAAGAAGACCATG
GGCTGGGAGGCCTCCTCCGAGCGGATGTACCCCGAGGACGGCGCCCTGAAGGGCGAGATCAAGCAGAGGCTGA
AGCTGAAGGACGGCGGCCACTACGACGCTGAGGTCAAGACCACCTACAAGGCCAAGAAGCCCGTGCAGCTGCC
CGGCGCCTACAACGTCAACATCAAGTTGGACATCACCTCCCACAACGAGGACTACACCATCGTGGAACAGTAC
GAACGCGCCGAGGGCCGCCACTCCACCGGCGGCATGGACGAGCTGTACAAGTCCGGACTCAGATCTCGAGCTC
AAGCTTCGAATTCTGCAGTCGACGGTACCGCGGGCCCGGGATCCATGAGCGACGTGGCTATTGTGAAGGAGGG
TTGGCTGCACAAACGAGGGGAGTACATCAAGACCTGGCGGCCACGCTACTTCCTCCTCAAGAATGATGGCACC
TTCATTGGCTACAAGGAGCGGCCGCAGGATGTGGACCAACGTGAGGCTCCCCTCAACAACTTCTCTGTGGCGC
AGTGCCAGCTGATGAAGACGGAGCGGCCCCGGCCCAACACCTTCATCATCCGCTGCCTGCAGTGGACCACTGT
CATCGAACGCACCTTCCATGTGGAGACTCCTGAGGAGCGGGAGGAGTGGACAACCGCCATCCAGACTGTGGCT
GACGGCCTCAAGAAGCAGGAGGAGGAGGAGATGGACTTCCGGTCGGGCTCACCCAGTGACAACTCAGGGGCTG
AAGAGATGGAGGTGTCCCTGGCCAAGCCCAAGCACCGCGTGACCATGAACTAAGCTCTAGAGTCGACGGGCCG
CGGTAACAATTGTTAACTAACTTAAGCTAGCAACGGTTTCCCTCTAGCGGGATCAATTCCG*CCCCCCCCCCCT*
*AACGTTACTGGCCGAAGCCGCTTGGAATAAGGCCGGTGTGCGTTTGTCTATATGTTATTTTCCACCATATTGC*
*CGTCTTTTGGCAATGTGAGGGCCCGGAAACCTGGCCCTGTCTTCTTGACGAGCATTCCTAGGGGTCTTTCCCC*
*TCTCGCCAAAGGAATGCAAGGTCTGTTGAATGTCGTGAAGGAAGCAGTTCCTCTGGAAGCTTCTTGAAGACAA*
*ACAACGTCTGTAGCGACCCTTTGCAGGCAGCGGAACCCCCCACCTGGCGACAGGTGCCTCTGCGGCCAAAAGC*
*CACGTGTATAAGATACACCTGCAAAGGCGGCACAACCCCAGTGCCACGTTGTGAGTTGGATAGTTGTGGAAAG*
*AGTCAAATGGCTCTCCTCAAGCGTATTCAACAAGGGGCTGAAGGATGCCCAGAAGGTACCCCATTGTATGGGA*
*TCTGATCTGGGGCCTCGGTGCACATGCTTTACATGTGTTTAGTCGAGGTTAAAAAACGTCTAGGCCCCCCGAA*
*CCACGGGGACGTGGTTTTCCTTTGAAAAACACGAT*AATACCATGGTCATGAAAACATTTAACATTTCTCAACA
AGATCTAGAATTAGTAGAAGTAGCGACAGAGAAGATTACAATGCTTTATGAGGATAATAAACATCATGTGGGA
GCGGCAATTCGTACGAAAACAGGAGAAATCATTTCGGCAGTACATATTGAAGCGTATATAGGACGAGTAACTG
TTTGTGCAGAAGCCATTGCGATTGGTAGTGCAGTTTCGAATGGACAAAAGGATTTTGACACGATTGTAGCTGT
TAGACACCCTTATTCTGACGAAGTAGATAGAAGTATTCGAGTGGTAAGTCCTTGTGGTATGTGTAGGGAGTTG
ATTTCAGACTATGCACCAGATTGTTTTGTGTTAATAGAAATGAATGGCAAGTTAGTCAAAACTACGATTGAAG
AACTCATTCCACTCAAATATACCCGAAATTAA

mCherry PH$_{Akt}$ [the PH domain of human Akt1 (residues 1–148)][S9] *IRES* Bsr$^{R}$

#### pPBpuro-mNG-eDHFR(69K6)-Tiam1

>Amino acid sequence
MVSKGEEDNMASLPATHELHIFGSINGVDFDMVGQGTGNPNDGYEELNLKSTKGDLQFSPWILVPHIGYGFHQ
YLPYPDGMSPFQAAMVDGSGYQVHRTMQFEDGASLTVNYRYTYEGSHIKGEAQVKGTGFPADGPVMTNSLTAA
DWCRSKKTYPNDKTIISTFKWSYTTGNGKRYRSTARTTYTFAKPMAANYLKNQPMYVFRKTELKHSKTELNFK
EWQKAFTDVMGMDELYKSGSAASSAGSMISLIAALAVDRVIGMENAMPWNLPADLAWFKRNTLNKPVIMGRHT
WESIGRPLPGRKNIILSSQPGTDKKKKKKDRVTWVKSVDEAIAACGDVPEIMVIGGGRVYEQFLPKAQKLYLT
HIDAEVEGDTHFPDYEPDDWESVFSEFHDADAQNSHSYCFEILERRLYKSGLRSRQGSGAGSGAGSGAGSGAG
SGAPRAMNPSDQNPSPQDSTGPQLATMRQLSDADNVRKVICELLETERTYVKDLNCLMERYLKPLQKETFLTQ
DELDVLFGNLTEMVEFQVEFLKTLEDGVRLVPDLEKLEKVDQFKKVLFSLGGSFLYYADRFKLYSAFCAIHTK
VPKVLVKAKTDTAFKAFLDAQNPKQQHSSTLESYLIKPIQRILKYPLLLRELFALTDAESEEHYHLDVAIKTM
NKVASHINEMQKIHEEFGAVFDQLIAEQTGEKKEVADLSMGDLLLHTTVIWLNPPASLGKWKKEPELAAFVFK
TAVVLVYKDGSKQKKKLVGSHRLSIYEDWDPFRFRHMIPTEALQVRALASADAEANAVCEIVHVKSESEGRPE
RVFHLCCSSPESRKDFLKAVHSILRDKHRRQLLKTESLPSSQQYVPFGGKRLCALKGARPAMSRAVSAPSKSL
GRRRRRLARNRFTIDSDAVSASSPEKESQQPPGGGDTDRWVEEQFDLAQYEEQDDIKETDILSDDDEFCESVK
GASVDRDLQERLQATSISQRERGRKTLDSHASRMAQLKKQAALSGINGGLESASEEVIWVRREDFAPSRKLNT
EI*-[IRES]-MTEYKPTVRLATRDDVPRAVRTLAAAFADYPATRHTVDPDRHIERVTELQELFLTRVGLDIG
KVWVADDGAAVAVWTTPESVEAGAVFAEIGPRMAELSGSRLAAQQQMEGLLAPHRPKEPAWFLATVGVSPDHQ
GKGLGSAVVLPGVEAAERAGVPAFLETSAPRNLPFYERLGFTVTADVECPKDRATWCMTRKPGA*

>DNA sequence
ATGGTGAGCAAGGGCGAGGAGGATAACATGGCCTCTCTCCCAGCGACACATGAGTTACACATCTTTGGCTCCA
TCAACGGTGTGGACTTTGACATGGTGGGTCAGGGCACCGGCAATCCAAATGATGGTTATGAGGAGTTAAACCT
GAAGTCCACCAAGGGTGACCTCCAGTTCTCCCCCTGGATTCTGGTCCCTCATATCGGGTATGGCTTCCATCAG
TACCTGCCCTACCCTGACGGGATGTCGCCTTTCCAGGCCGCCATGGTAGATGGCTCCGGCTACCAAGTCCATC
GCACAATGCAGTTTGAAGATGGTGCCTCCCTTACTGTTAACTACCGCTACACCTACGAGGGAAGCCACATCAA
AGGAGAGGCCCAGGTGAAGGGGACTGGTTTCCCTGCTGACGGTCCTGTGATGACCAACTCGCTGACCGCTGCG
GACTGGTGCAGGTCGAAGAAGACTTACCCCAACGACAAAACCATCATCAGTACCTTTAAGTGGAGTTACACCA
CTGGAAATGGCAAGCGCTACCGGAGCACTGCGCGGACCACCTACACCTTTGCCAAGCCAATGGCGGCTAACTA
TCTGAAGAACCAGCCGATGTACGTGTTCCGTAAGACGGAGCTCAAGCACTCCAAGACCGAGCTCAACTTCAAG
GAGTGGCAAAAGGCCTTTACCGATGTGATGGGCATGGACGAGCTCTACAAGTCTGGGTCCGCAGCCAGTAGCG
CCGGAAGTATGATCAGTCTGATTGCGGCGTTAGCGGTAGATCGCGTTATCGGCATGGAAAACGCCATGCCGTG
GAACCTGCCTGCCGATCTCGCCTGGTTTAAACGCAACACCTTAAATAAACCCGTGATTATGGGCCGCCATACC
TGGGAATCAATCGGTCGTCCGTTGCCAGGACGCAAAAATATTATCCTCAGCAGTCAACCGGGTACGGACAAAA
AAAAGAAAAAGAAAGATCGCGTAACGTGGGTGAAGTCGGTGGATGAAGCCATCGCGGCGTGTGGTGACGTACC
AGAAATCATGGTGATTGGCGGCGGTCGCGTTTATGAACAGTTCTTGCCAAAAGCGCAAAAACTGTATCTGACG
CATATCGACGCAGAAGTGGAAGGCGACACCCATTTCCCGGATTACGAGCCGGATGACTGGGAATCGGTATTCA
GCGAATTCCACGATGCTGATGCGCAGAACTCTCACAGCTATTGCTTTGAGATTCTGGAGCGGCGGCTGTACAA
GTCCGGACTCAGATCTCGACAAGGTAGTGGTGCTGGCTCTGGTGCTGGTAGTGGCGCTGGTTCCGGTGCTGGC
TCTGGCGCGCCTCGAGCAATGAACCCCTCTGACCAGAACCCATCTCCTCAGGACTCCACGGGGCCTCAGCTGG
CGACCATGAGACAACTCTCGGATGCAGATAACGTGCGCAAGGTGATCTGCGAGCTCCTGGAGACGGAGCGCAC
CTACGTGAAGGATTTAAACTGTCTTATGGAGAGATACCTAAAGCCTCTTCAAAAAGAAACTTTTCTCACCCAG
GATGAGCTTGACGTGCTTTTTGGAAATTTAACGGAAATGGTAGAGTTTCAAGTAGAATTCCTTAAAACTCTAG
AAGATGGAGTGAGACTGGTACCTGATTTGGAAAAGCTTGAGAAGGTTGATCAATTTAAGAAAGTGCTGTTCTC
TCTGGGGGGATCATTCCTGTATTATGCTGACCGCTTCAAGCTCTACAGTGCCTTCTGCGCCATCCACACAAAA
GTTCCCAAGGTCCTGGTGAAAGCCAAGACAGACACGGCTTTCAAGGCATTCTTGGATGCCCAGAACCCGAAGC
AGCAGCACTCATCCACGCTGGAGTCGTACCTCATCAAGCCCATCCAGAGGATCCTCAAGTACCCACTTCTGCT
CAGGGAGCTGTTCGCCCTGACCGATGCGGAGAGCGAGGAGCACTACCACCTGGACGTGGCCATCAAGACCATG
AACAAGGTTGCCAGTCACATCAATGAGATGCAGAAAATCCATGAAGAGTTTGGGGCTGTGTTTGACCAGCTGA
TTGCTGAACAGACTGGTGAGAAAAAAGAGGTTGCAGATCTGAGCATGGGAGACCTGCTTTTGCACACTACCGT
GATCTGGCTGAACCCGCCGGCCTCGCTGGGCAAGTGGAAAAAGGAACCAGAGTTGGCAGCATTCGTCTTCAAA
ACTGCTGTGGTCCTTGTGTATAAAGATGGTTCCAAACAGAAGAAGAAACTTGTAGGATCTCACAGGCTTTCCA
TTTATGAGGACTGGGACCCCTTCAGATTTCGACACATGATCCCCACGGAAGCGCTGCAGGTTCGAGCTTTGGC
GAGTGCAGATGCAGAGGCAAATGCCGTGTGTGAAATTGTCCATGTAAAATCCGAGTCTGAAGGGAGGCCGGAG
AGGGTCTTTCACTTGTGCTGCAGCTCCCCAGAGAGCCGAAAGGATTTCCTAAAGGCTGTGCATTCAATCCTGC
GTGATAAGCACAGAAGACAGCTCCTCAAAACCGAGAGCCTTCCCTCATCCCAGCAATATGTCCCTTTTGGAGG

CAAAAGATTGTGTGCACTGAAGGGGGCCAGGCCGGCCATGAGCAGGGCAGTGTCTGCCCCAAGCAAGTCTCTT
GGGAGGAGGAGGCGGCGGCTGGCTCGAAACAGGTTTACCATTGATTCTGATGCCGTCTCCGCAAGCAGCCCGG
AGAAAGAGTCCCAGCAGCCCCCGGTGGTGGGACACTGACCGATGGGTAGAGGAGCAGTTTGATCTTGCTCA
GTATGAGGAGCAAGATGACATCAAGGAGACAGACATCCTCAGTGACGATGATGAGTTCTGTGAGTCCGTGAAG
GGTGCCTCAGTGGACAGAGACCTGCAGGAGCGGCTTCAGGCCACCTCCATCAGTCAGCGGGAAAGAGGCCGGA
AAACCCTGGATAGTCACGCGTCCCGCATGGCACAGCTCAAGAAGCAAGCTGCCCTGTCGGGGATCAATGGAGG
CCTGGAGAGCGCAAGCGAGGAAGTCATTTGGGTTAGGCGTGAAGACTTTGCCCCCTCCAGGAAACTGAACACT
GAGATCTGACCGCGGGCCCGGGATAAGTCAACTAACTTAAGCTAGCAACGGTTTCCCTCTAGCGGGATCAATT
CCG*CCCCCCCCCCCTAACGTTACTGGCCGAAGCCGCTTGGAATAAGGCCGGTGTGCGTTTGTCTATATGTTAT*
*TTTCCACCATATTGCCGTCTTTTGGCAATGTGAGGGCCCGGAAACCTGGCCCTGTCTTCTTGACGAGCATTCC*
*TAGGGGTCTTTCCCCTCTCGCCAAAGGAATGCAAGGTCTGTTGAATGTCGTGAAGGAAGCAGTTCCTCTGGAA*
*GCTTCTTGAAGACAAACAACGTCTGTAGCGACCCTTTGCAGGCAGCGGAACCCCCCACCTGGCGACAGGTGCC*
*TCTGCGGCCAAAAGCCACGTGTATAAGATACACCTGCAAAGGCGGCACAACCCCAGTGCCACGTTGTGAGTTG*
*GATAGTTGTGGAAAGAGTCAAATGGCTCTCCTCAAGCGTATTCAACAAGGGGCTGAAGGATGCCCAGAAGGTA*
*CCCCATTGTATGGATCTGATCTGGGGCCTCGGTGCACATGCTTTACATGTGTTTAGTCGAGGTTAAAAAACG*
*TCTAGGCCCCCCGAACCACGGGGACGTGGTTTTCCTTTGAAAAACACGAT*AATACCATGACCGAGTACAAGCC
CACGGTGCGCCTCGCCACCCGCGACGACGTCCCCAGGGCCGTACGCACCCTCGCCGCCGCGTTCGCCGACTAC
CCCGCCACGCGCCACACCGTCGATCCGGACCGCCACATCGAGCGGGTCACCGAGCTGCAAGAACTCTTCCTCA
CGCGCGTCGGGCTCGACATCGGCAAGGTGTGGGTCGCGGACGACGGCGCCGCGGTGGCGGTCTGGACCACGCC
GGAGAGCGTCGAAGCGGGGGCGGTGTTCGCCGAGATCGGCCCGCGCATGGCCGAGTTGAGCGGTTCCCGGCTG
GCCGCGCAGCAACAGATGGAAGGCCTCCTGGCGCCGCACCGGCCCAAGGAGCCCGCGTGGTTCCTGGCCACCG
TCGGCGTCTCGCCCGACCACCAGGGCAAGGGTCTGGGCAGCGCCGTCGTGCTCCCCGGAGTGGAGGCGGCCGA
GCGCGCCGGGGTGCCCGCCTTCCTGGAGACCTCCGCGCCCCGCAACCTCCCCTTCTACGAGCGGCTCGGCTTC
ACCGTCACCGCCGACGTCGAGTGCCCGAAGGACCGCGCGACCTGGTGCATGACCCGCAAGCCCGGTGCCTGA

mNeonGreen eDHFR K6-tag(69K6) Tiam1 [the DH-PH domain of human Tiam1 (residues 1012–1591)][S10] *IRES* Puro$^R$

**pPBbsr-Lifeact-mCherry**

>Amino acid sequence
MGVADLIKKFESISKEEGDPPVATMVSKGEEDNMAIIKEFMRFKVHMEGSVNGHEFEIEGEGEGRPYEGTQTA
KLKVTKGGPLPFAWDILSPQFMYGSKAYVKHPADIPDYLKLSFPEGFKWERVMNFEDGGVVTVTQDSSLQDGE
FIYKVKLRGTNFPSDGPVMQKKTMGWEASSERMYPEDGALKGEIKQRLKLKDGGHYDAEVKTTYKAKKPVQLP
GAYNVNIKLDITSHNEDYTIVEQYERAEGRHSTGGMDELYK*-[*IRES*]-MLYEDNKHHVGAAIRTKTGEIIS
AVHIEAYIGRVTVCAEAIAIGSAVSNGQKDFDTIVAVRHPYSDEVDRSIRVVSPCGMCRELISDYAPDCFVLI
EMNGKLVKTTIEELIPLKYTRN*

>DNA sequence
ATGGGCGTGGCCGACTTGATCAAGAAGTTCGAGTCCATCTCCAAGGAGGAGGGGGATCCACCGGTCGCCACCA
TGGTGAGCAAGGGCGAGGAGGATAACATGGCCATCATCAAGGAGTTCATGCGCTTCAAGGTGCACATGGAGGG
CTCCGTGAACGGCCACGAGTTCGAGATCGAGGGCGAGGGCGAGGGCCGCCCCTACGAGGGCACCCAGACCGCC
AAGCTGAAGGTGACCAAGGGTGGCCCCCTGCCCTTCGCCTGGGACATCCTGTCCCCTCAGTTCATGTACGGCT
CCAAGGCCTACGTGAAGCACCCCGCCGACATCCCCGACTACTTGAAGCTGTCCTTCCCCGAGGGCTTCAAGTG
GGAGCGCGTGATGAACTTCGAGGACGGCGGCGTGGTGACCGTGACCCAGGACTCCTCCCTGCAGGACGGCGAG
TTCATCTACAAGGTGAAGCTGCGCGGCACCAACTTCCCCTCCGACGGCCCCGTAATGCAGAAGAAGACCATGG
GCTGGGAGGCCTCCTCCGAGCGGATGTACCCCGAGGACGGCGCCCTGAAGGGCGAGATCAAGCAGAGGCTGAA
GCTGAAGGACGGCGGCCACTACGACGCTGAGGTCAAGACCACCTACAAGGCCAAGAAGCCCGTGCAGCTGCCC
GGCGCCTACAACGTCAACATCAAGTTGGACATCACCTCCCACAACGAGGACTACACCATCGTGGAACAGTACG
AACGCGCCGAGGGCCGCCACTCCACCGGCGGCATGGACGAGCTGTACAAGTAAAGCGGCCGCTCTAGAGTCGA
CGGGCCGCGGTAACAATTGTTAACTAACTTAAGCTAGCAACGGTTTCCCTCTAGCGGGATCAATTCCG*CCCCC*
*CCCCCCTAACGTTACTGGCCGAAGCCGCTTGGAATAAGGCCGGTGTGCGTTTGTCTATATGTTATTTTCCACC*
*ATATTGCCGTCTTTTGGCAATGTGAGGGCCCGGAAACCTGGCCCTGTCTTCTTGACGAGCATTCCTAGGGGTC*
*TTTCCCCTCTCGCCAAAGGAATGCAAGGTCTGTTGAATGTCGTGAAGGAAGCAGTTCCTCTGGAAGCTTCTTG*
*AAGACAAACAACGTCTGTAGCGACCCTTTGCAGGCAGCGGAACCCCCCACCTGGCGACAGGTGCCTCTGCGGC*
*CAAAAGCCACGTGTATAAGATACACCTGCAAAGGCGGCACAACCCCAGTGCCACGTTGTGAGTTGGATAGTTG*
*TGGAAAGAGTCAAATGGCTCTCCTCAAGCGTATTCAACAAGGGGCTGAAGGATGCCCAGAAGGTACCCCATTG*
*TATGGGATCTGATCTGGGGCCTCGGTGCACATGCTTTACATGTGTTTAGTCGAGGTTAAAAAACGTCTAGGCC*
*CCCCGAACCACGGGGACGTGGTTTTCCTTTGAAAAACACGAT*AATACCATGGTCATGAAAACATTTAACATTT
CTCAACAAGATCTAGAATTAGTAGAAGTAGCGACAGAGAAGATTACAATGCTTTATGAGGATAATAAACATCA
TGTGGGAGCGGCAATTCGTACGAAAACAGGAGAAATCATTTCGGCAGTACATATTGAAGCGTATATAGGACGA
GTAACTGTTTGTGCAGAAGCCATTGCGATTGGTAGTGCAGTTTCGAATGGACAAAAGGATTTTGACACGATTG
TAGCTGTTAGACACCCTTATTCTGACGAAGTAGATAGAAGTATTCGAGTGGTAAGTCCTTGTGGTATGTGTAG
GGAGTTGATTTCAGACTATGCACCAGATTGTTTTGTGTTAATAGAAATGAATGGCAAGTTAGTCAAAACTACG
ATTGAAGAACTCATTCCACTCAAATATACCCGAAATTAA

Lifeact [the F-actin targeting motif of *Saccharomyces cerevisiae* Abp140 (residues 1–17)][S11] mCherry *IRES* Bsr$^{R}$

**pPBpuro-eDHFR(69K6)-EGFP**

```
>Amino acid sequence
MISLIAALAVDRVIGMENAMPWNLPADLAWFKRNTLNKPVIMGRHTWESIGRPLPGRKNIILSSQPGTDKKKK
KKDRVTWVKSVDEAIAACGDVPEIMVIGGGRVYEQFLPKAQKLYLTHIDAEVEGDTHFPDYEPDDWESVFSEF
HDADAQNSHSYCFEILERRSGSGDPPVATMVSKGEELFTGVVPILVELDGDVNGHKFSVSGEGEGDATYGKLT
LKFICTTGKLPVPWPTLVTTLTYGVQCFSRYPDHMKQHDFFKSAMPEGYVQERTIFFKDDGNYKTRAEVKFEG
DTLVNRIELKGIDFKEDGNILGHKLEYNYNSHNVYIMADKQKNGIKVNFKIRHNIEDGSVQLADHYQQNTPIG
DGPVLLPDNHYLSTQSALSKDPNEKRDHMVLLEFVTAAGITLGMDELYK*-[IRES]-MTEYKPTVRLATRDD
VPRAVRTLAAAFADYPATRHTVDPDRHIERVTELQELFLTRVGLDIGKVWVADDGAAVAVWTTPESVEAGAVF
AEIGPRMAELSGSRLAAQQQMEGLLAPHRPKEPAWFLATVGVSPDHQGKGLGSAVVLPGVEAAERAGVPAFLE
TSAPRNLPFYERLGFTVTADVECPKDRATWCMTRKPGA*
```

```
>DNA sequence
ATGATCAGTCTGATTGCGGCGTTAGCGGTAGATCGCGTTATCGGCATGGAAAACGCCATGCCGTGGAACCTGC
CTGCCGATCTCGCCTGGTTTAAACGCAACACCTTAAATAAACCCGTGATTATGGGCCGCCATACCTGGGAATC
AATCGGTCGTCCGTTGCCAGGACGCAAAAATATTATCCTCAGCAGTCAACCGGGTACGGACAAAAAAAAGAAA
AAGAAAGATCGCGTAACGTGGGTGAAGTCGGTGGATGAAGCCATCGCGGCGTGTGGTGACGTACCAGAAATCA
TGGTGATTGGCGGCGGTCGCGTTTATGAACAGTTCTTGCCAAAAGCGCAAAAACTGTATCTGACGCATATCGA
CGCAGAAGTGGAAGGCGACACCCATTTCCCGGATTACGAGCCGGATGACTGGGAATCGGTATTCAGCGAATTC
CACGATGCTGATGCGCAGAACTCTCACAGCTATTGCTTTGAGATTCTGGAGCGGCGGAGCGGCTCCGGGGATC
CACCGGTCGCCACCATGGTGAGCAAGGGCGAGGAGCTGTTCACCGGGGTGGTGCCCATCCTGGTCGAGCTGGA
CGGCGACGTAAACGGCCACAAGTTCAGCGTGTCCGGCGAGGGCGAGGGCGATGCCACCTACGGCAAGCTGACC
CTGAAGTTCATCTGCACCACCGGCAAGCTGCCCGTGCCCTGGCCCACCCTCGTGACCACCCTGACCTACGGCG
TGCAGTGCTTCAGCCGCTACCCCGACCACATGAAGCAGCACGACTTCTTCAAGTCCGCCATGCCCGAAGGCTA
CGTCCAGGAGCGCACCATCTTCTTCAAGGACGACGGCAACTACAAGACCCGCGCCGAGGTGAAGTTCGAGGGC
GACACCCTGGTGAACCGCATCGAGCTGAAGGGCATCGACTTCAAGGAGGACGGCAACATCCTGGGGCACAAGC
TGGAGTACAACTACAACAGCCACAACGTCTATATCATGGCCGACAAGCAGAAGAACGGCATCAAGGTGAACTT
CAAGATCCGCCACAACATCGAGGACGGCAGCGTGCAGCTCGCCGACCACTACCAGCAGAACACCCCCATCGGC
GACGGCCCCGTGCTGCTGCCCGACAACCACTACCTGAGCACCCAGTCCGCCCTGAGCAAAGACCCCAACGAGA
AGCGCGATCACATGGTCCTGCTGGAGTTCGTGACCGCCGCCGGGATCACTCTCGGCATGGACGAGCTGTACAA
GTAAAGCGGCCGCTCTAGAGTCAACTAACTTAAGCTAGCAACGGTTTCCCTCTAGCGGGATCAATTCCGCCCC
CCCCCCCTAACGTTACTGGCCGAAGCCGCTTGGAATAAGGCCGGTGTGCGTTTGTCTATATGTTATTTTCCAC
CATATTGCCGTCTTTTGGCAATGTGAGGGCCCGGAAACCTGGCCCTGTCTTCTTGACGAGCATTCCTAGGGGT
CTTTCCCCTCTCGCCAAAGGAATGCAAGGTCTGTTGAATGTCGTGAAGGAAGCAGTTCCTCTGGAAGCTTCTT
GAAGACAAACAACGTCTGTAGCGACCCTTTGCAGGCAGCGGAACCCCCCACCTGGCGACAGGTGCCTCTGCGG
CCAAAAGCCACGTGTATAAGATACACCTGCAAAGGCGGCACAACCCCAGTGCCACGTTGTGAGTTGGATAGTT
GTGGAAAGAGTCAAATGGCTCTCCTCAAGCGTATTCAACAAGGGGCTGAAGGATGCCCAGAAGGTACCCATT
GTATGGGATCTGATCTGGGGCCTCGGTGCACATGCTTTACATGTGTTTAGTCGAGGTTAAAAAACGTCTAGGC
CCCCCGAACCACGGGGACGTGGTTTTCCTTTGAAAAACACGATAATACCATGACCGAGTACAAGCCCACGGTG
CGCCTCGCCACCCGCGACGACGTCCCCAGGGCCGTACGCACCCTCGCCGCCGCGTTCGCCGACTACCCCGCCA
CGCGCCACACCGTCGATCCGGACCGCCACATCGAGCGGGTCACCGAGCTGCAAGAACTCTTCCTCACGCGCGT
CGGGCTCGACATCGGCAAGGTGTGGGTCGCGGACGACGGCGCCGCGGTGGCGGTCTGGACCACGCCGGAGAGC
GTCGAAGCGGGGGCGGTGTTCGCCGAGATCGGCCCGCGCATGGCCGAGTTGAGCGGTTCCCGGCTGGCCGCGC
AGCAACAGATGGAAGGCCTCCTGGCGCCGCACCGGCCCAAGGAGCCCGCGTGGTTCCTGGCCACCGTCGGCGT
CTCGCCCGACCACCAGGGCAAGGGTCTGGGCAGCGCCGTCGTGCTCCCCGGAGTGGAGGCGGCCGAGCGCGCC
GGGGTGCCCGCCTTCCTGGAGACCTCCGCGCCCCGCAACCTCCCCTTCTACGAGCGGCTCGGCTTCACCGTCA
CCGCCGACGTCGAGTGCCCGAAGGACCGCGACCTGGTGCATGACCCGCAAGCCCGGTGCCTGA
```

eDHFR K6-tag(69K6) EGFP *IRES* Puro$^{R}$

**pPBpuro-cRaf-mNG-eDHFR(69K6)**

```
>Amino acid sequence
MEHIQGAWKTISNGFGFKDAVFDGSSCISPTIVQQFGYQRRASDDGKLTDPSKTSNTIRVFLPNKQRTVVNVR
NGMSLHDCLMKALKVRGLQPECCAVFRLLHEHKGKKARLDWNTDAASLIGEELQVDFLDHVPLTTHNFARKTF
LKLAFCDICQKFLLNGFRCQTCGYKFHEHCSTKVPTMCVDWSNIRQLLLFPNSTIGDSGVPALPSLTMRRMRE
SVSRMPVSSQHRYSTPHAFTFNTSSPSSEGSLSQRQRSTSTPNVHMVSTTLPVDSRMIEDAIRSHSESASPSA
LSSSPNNLSPTGWSQPKTPVPAQRERAPVSGTQEKNKIRPRGQRDSSYYWEIEASEVMLSTRIGSGSFGTVYK
GKWHGDVAVKILKVVDPTPEQFQAFRNEVAVLRKTRHVNILLFMGYMTKDNLAIVTQWCEGSSLYKHLHVQET
KFQMFQLIDIARQTAQGMDYLHAKNIIHRDMKSNNIFLHEGLTVKIGDFGLATVKSRWSGSQQVEQPTGSVLW
MAPEVIRMQDNNPFSFQSDVYSYGIVLYELMTGELPYSHINNRDQIIFMVGRGYASPDLSKLYKNCPKAMKRL
VADCVKKVKEERPLFPQILSSIELLQHSLPKINRSASEPSLHRAAHTEDINACTLTTSPRLPVFDPPVATMVS
KGEEDNMASLPATHELHIFGSINGVDFDMVGQGTGNPNDGYEELNLKSTKGDLQFSPWILVPHIGYGFHQYLP
YPDGMSPFQAAMVDGSGYQVHRTMQFEDGASLTVNYRYTYEGSHIKGEAQVKGTGFPADGPVMTNSLTAADWC
RSKKTYPNDKTIISTFKWSYTTGNGKRYRSTARTTYTFAKPMAANYLKNQPMYVFRKTELKHSKTELNFKEWQ
KAFTDVMGMDELYKSGSAASSAGSMISLIAALAVDRVIGMENAMPWNLPADLAWFKRNTLNKPVIMGRHTWES
IGRPLPGRKNIILSSQPGTDKKKKKKDRVTWVKSVDEAIAACGDVPEIMVIGGGRVYEQFLPKAQKLYLTHID
AEVEGDTHFPDYEPDDWESVFSEFHDADAQNSHSYCFEILERR*-[IRES]-MTEYKPTVRLATRDDVPRAVR
TLAAAFADYPATRHTVDPDRHIERVTELQELFLTRVGLDIGKVWVADDGAAVAVWTTPESVEAGAVFAEIGPR
MAELSGSRLAAQQQMEGLLAPHRPKEPAWFLATVGVSPDHQGKGLGSAVVLPGVEAAERAGVPAFLETSAPRN
LPFYERLGFTVTADVECPKDRATWCMTRKPGA*

>DNA sequence
ATGGAGCACATACAGGGAGCTTGGAAGACGATCAGCAATGGTTTTGGATTCAAAGATGCCGTGTTTGATGGCT
CCAGCTGCATCTCTCCTACAATAGTTCAGCAGTTTGGCTATCAGCGCCGGGCATCAGATGATGGCAAACTCAC
AGATCCTTCTAAGACAAGCAACACTATCCGTGTTTTCTTGCCGAACAAGCAAAGAACAGTGGTCAATGTGCGA
AATGGAATGAGCTTGCATGACTGCCTTATGAAAGCACTCAAGGTGAGGGGCCTGCAACCAGAGTGCTGTGCAG
TGTTCAGACTTCTCCACGAACACAAAGGTAAAAAAGCACGCTTAGATTGGAATACTGATGCTGCGTCTTTGAT
TGGAGAAGAACTTCAAGTAGATTTCCTGGATCATGTTCCCCTCACAACACACAACTTTGCTCGGAAGACGTTC
CTGAAGCTTGCCTTCTGTGACATCTGTCAGAAATTCCTGCTCAATGGATTTCGATGTCAGACTTGTGGCTACA
AATTTCATGAGCACTGTAGCACCAAAGTACCTACTATGTGTGTGGACTGGAGTAACATCAGACAACTCTTATT
GTTTCCAAATTCCACTATTGGTGATAGTGGAGTCCCAGCACTACCTTCTTTGACTATGCGTCGTATGCGAGAG
TCTGTTTCCAGGATGCCTGTTAGTTCTCAGCACAGATATTCTACACCTCACGCCTTCACCTTTAACACCTCCA
GTCCCTCATCTGAAGGTTCCCTCTCCCAGAGGCAGAGGTCGACATCCACACCTAATGTCCACATGGTCAGCAC
CACCCTGCCTGTGGACAGCAGGATGATTGAGGATGCAATTCGAAGTCACAGCGAATCAGCCTCACCTTCAGCC
CTGTCCAGTAGCCCCAACAATCTGAGCCCAACAGGCTGGTCACAGCCGAAAACCCCCGTGCCAGCACAAAGAG
AGCGGGCACCAGTATCTGGGACCCAGGAGAAAAACAAAATTAGGCCTCGTGGACAGAGAGATTCAAGCTATTA
TTGGGAAATAGAAGCCAGTGAAGTGATGCTGTCCACTCGGATTGGGTCAGGCTCTTTTGGAACTGTTTATAAG
GGTAAATGGCACGGAGATGTTGCAGTAAAGATCCTAAAGGTTGTCGACCCAACCCCAGAGCAATTCCAGGCCT
TCAGGAATGAGGTGGCTGTTCTGCGCAAAACACGGCATGTGAACATTCTGCTTTTCATGGGGTACATGACAAA
GGACAACCTGGCAATTGTGACCCAGTGGTGCGAGGGCAGCAGCCTCTACAAACACCTGCATGTCCAGGAGACC
AAGTTTCAGATGTTCCAGCTAATTGACATTGCCCGGCAGACGGCTCAGGGAATGGACTATTTGCATGCAAAGA
ACATCATCCATAGAGACATGAAATCCAACAATATATTTCTCCATGAAGGCTTAACAGTGAAAATTGGAGATTT
TGGTTTGGCAACAGTAAAGTCACGCTGGAGTGGTTCTCAGCAGGTTGAACAACCTACTGGCTCTGTCCTCTGG
ATGGCCCCAGAGGTGATCCGAATGCAGGATAACAACCCATTCAGTTTCCAGTCGGATGTCTACTCCTATGGCA
TCGTATTGTATGAACTGATGACGGGGGAGCTTCCTTATTCTCACATCAACAACCGAGATCAGATCATCTTCAT
GGTGGGCCGAGGATATGCCTCCCCAGATCTTAGTAAGCTATATAAGAACTGCCCCAAAGCAATGAAGAGGCTG
GTAGCTGACTGTGTGAAGAAAGTAAAGGAAGAGAGGCCTCTTTTTCCCCAGATCCTGTCTTCCATTGAGCTGC
TCCAACACTCTCTACCGAAGATCAACCGGAGCGCTTCCGAGCCATCCTTGCATCGGGCAGCCCACACTGAGGA
TATCAATGCTTGCACGCTGACCACGTCCCCGAGGCTGCCTGTCTTCGATCCACCGGTCGCCACCATGGTGAGC
AAGGGCGAGGAGGATAACATGGCCTCTCTCCCAGCGACACATGAGTTACACATCTTTGGCTCCATCAACGGTG
TGGACTTTGACATGGTGGGTCAGGGCACCGGCAATCCAAATGATGGTTATGAGGAGTTAAACCTGAAGTCCAC
CAAGGGTGACCTCCAGTTCTCCCCCTGGATTCTGGTCCCTCATATCGGGTATGGCTTCCATCAGTACCTGCCC
TACCCTGACGGGATGTCGCCTTTCCAGGCCGCCATGGTAGATGGCTCCGGCTACCAAGTCCATCGCACAATGC
AGTTTGAAGATGGTGCCTCCCTTACTGTTAACTACCGCTACACCTACGAGGGAAGCCACATCAAAGGAGAGGC
CCAGGTGAAGGGGACTGGTTTCCCTGCTGACGGTCCTGTGATGACCAACTCGCTGACCGCTGCGGACTGGTGC
AGGTCGAAGAAGACTTACCCCAACGACAAAACCATCATCAGTACCTTTAAGTGGAGTTACACCACTGGAAATG
```

GCAAGCGCTACCGGAGCACTGCGCGGACCACCTACACCTTTGCCAAGCCAATGGCGGCTAACTATCTGAAGAA
CCAGCCGATGTACGTGTTCCGTAAGACGGAGCTCAAGCACTCCAAGACCGAGCTCAACTTCAAGGAGTGGCAA
AAGGCCTTTACCGATGTGATGGGCATGGACGAGCTGTACAAGTCTGGGTCCGCAGCCAGTAGCGCCGGAAGTA
TGATCAGTCTGATTGCGGCGTTAGCGGTAGATCGCGTTATCGGCATGGAAAACGCCATGCCGTGGAACCTGCC
TGCCGATCTCGCCTGGTTTAAACGCAACACCTTAAATAAACCCGTGATTATGGGCCGCCATACCTGGGAATCA
ATCGGTCGTCCGTTGCCAGGACGCAAAAATATTATCCTCAGCAGTCAACCGGGTACGGACAAAAAAAAGAAAA
AGAAAGATCGCGTAACGTGGGTGAAGTCGGTGGATGAAGCCATCGCGGCGTGTGGTGACGTACCAGAAATCAT
GGTGATTGGCGGCGGTCGCGTTTATGAACAGTTCTTGCCAAAAGCGCAAAAACTGTATCTGACGCATATCGAC
GCAGAAGTGGAAGGCGACACCCATTTCCCGGATTACGAGCCGGATGACTGGGAATCGGTATTCAGCGAATTCC
ACGATGCTGATGCGCAGAACTCTCACAGCTATTGCTTTGAGATTCTGGAGCGGCGGTAACCCGGGATAAGTCA
ACTAACTTAAGCTAGCAACGGTTTCCCTCTAGCGGGATCAATTCCG*CCCCCCCCCCTAACGTTACTGGCCGA*
*AGCCGCTTGGAATAAGGCCGGTGTGCGTTTGTCTATATGTTATTTTCCACCATATTGCCGTCTTTTGGCAATG*
*TGAGGGCCCGGAAACCTGGCCCTGTCTTCTTGACGAGCATTCCTAGGGGTCTTTCCCCTCTCGCCAAAGGAAT*
*GCAAGGTCTGTTGAATGTCGTGAAGGAAGCAGTTCCTCTGGAAGCTTCTTGAAGACAAACAACGTCTGTAGCG*
*ACCCTTTGCAGGCAGCGGAACCCCCCACCTGGCGACAGGTGCCTCTGCGGCCAAAAGCCACGTGTATAAGATA*
*CACCTGCAAAGGCGGCACAACCCCAGTGCCACGTTGTGAGTTGGATAGTTGTGGAAAGAGTCAAATGGCTCTC*
*CTCAAGCGTATTCAACAAGGGGCTGAAGGATGCCCAGAAGGTACCCCATTGTATGGGATCTGATCTGGGGCCT*
*CGGTGCACATGCTTTACATGTGTTTAGTCGAGGTTAAAAAACGTCTAGGCCCCCCGAACCACGGGGACGTGGT*
*TTTCCTTTGAAAAACACGAT*AATACCATGACCGAGTACAAGCCCACGGTGCGCCTCGCCACCCGCGACGACGT
CCCCAGGGCCGTACGCACCCTCGCCGCCGCGTTCGCCGACTACCCCGCCACGCGCCACACCGTCGATCCGGAC
CGCCACATCGAGCGGGTCACCGAGCTGCAAGAACTCTTCCTCACGCGCGTCGGGCTCGACATCGGCAAGGTGT
GGGTCGCGGACGACGGCGCCGCGGTGGCGGTCTGGACCACGCCGGAGAGCGTCGAAGCGGGGGCGGTGTTCGC
CGAGATCGGCCCGCGCATGGCCGAGTTGAGCGGTTCCCGGCTGGCCGCGCAGCAACAGATGGAAGGCCTCCTG
GCGCCGCACCGGCCCAAGGAGCCCGCGTGGTTCCTGGCCACCGTCGGCGTCTCGCCCGACCACCAGGGCAAGG
GTCTGGGCAGCGCCGTCGTGCTCCCCGGAGTGGAGGCGGCCGAGCGCGCCGGGGTGCCCGCCTTCCTGGAGAC
CTCCGCGCCCCGCAACCTCCCCTTCTACGAGCGGCTCGGCTTCACCGTCACCGCCGACGTCGAGTGCCCGAAG
GACCGCGCGACCTGGTGCATGACCCGCAAGCCCGGTGCCTGA

cRaf(human) mNeonGreen eDHFR K6-tag(69K6) *IRES* Puro$^{R}$

**pT7-eDHFR/pET41**

>Amino acid sequence
MISLIAALAVDRVIGMENAMPWNLPADLAWFKRNTLNKPVIMGRHTWESIGRPLPGRKNIILSSQPGTDDRVT
WVKSVDEAIAACGDVPEIMVIGGGRVYEQFLPKAQKLYLTHIDAEVEGDTHFPDYEPDDWESVFSEFHDADAQ
NSHSYCFEILERRVDKLAAALEHHHHHHHH*

>DNA sequence
ATGATCAGTCTGATTGCGGCGTTAGCGGTAGATCGCGTTATCGGCATGGAAAACGCCATGCCGTGGAACCTGC
CTGCCGATCTCGCCTGGTTTAAACGCAACACCTTAAATAAACCCGTGATTATGGGCCGCCATACCTGGGAATC
AATCGGTCGTCCGTTGCCAGGACGCAAAAATATTATCCTCAGCAGTCAACCGGGTACGGACGATCGCGTAACG
TGGGTGAAGTCGGTGGATGAAGCCATCGCGGCGTGTGGTGACGTACCAGAAATCATGGTGATTGGCGGCGGTC
GCGTTTATGAACAGTTCTTGCCAAAAGCGCAAAAACTGTATCTGACGCATATCGACGCAGAAGTGGAAGGCGA
CACCCATTTCCCGGATTACGAGCCGGATGACTGGGAATCGGTATTCAGCGAATTCCACGATGCTGATGCGCAG
AACTCTCACAGCTATTGCTTTGAGATTCTGGAGCGGCGGGTCGACAAGCTTGCGGCCGCACTCGAGCACCACC
ACCACCACCACCACCACTAA

eDHFR His-tag

**pT7-eDHFR(69K6)/pET41**

>Amino acid sequence
MISLIAALAVDRVIGMENAMPWNLPADLAWFKRNTLNKPVIMGRHTWESIGRPLPGRKNIILSSQPGTDKKKK
KKDRVTWVKSVDEAIAACGDVPEIMVIGGGRVYEQFLPKAQKLYLTHIDAEVEGDTHFPDYEPDDWESVFSEF
HDADAQNSHSYCFEILERRVDKLAAALEHHHHHHHH*

>DNA sequence
ATGATCAGTCTGATTGCGGCGTTAGCGGTAGATCGCGTTATCGGCATGGAAAACGCCATGCCGTGGAACCTGC
CTGCCGATCTCGCCTGGTTTAAACGCAACACCTTAAATAAACCCGTGATTATGGGCCGCCATACCTGGGAATC
AATCGGTCGTCCGTTGCCAGGACGCAAAAATATTATCCTCAGCAGTCAACCGGGTACGGACAAAAAAAAGAAA
AAGAAAGATCGCGTAACGTGGGTGAAGTCGGTGGATGAAGCCATCGCGGCGTGTGGTGACGTACCAGAAATCA
TGGTGATTGGCGGCGGTCGCGTTTATGAACAGTTCTTGCCAAAAGCGCAAAAACTGTATCTGACGCATATCGA
CGCAGAAGTGGAAGGCGACACCCATTTCCCGGATTACGAGCCGGATGACTGGGAATCGGTATTCAGCGAATTC
CACGATGCTGATGCGCAGAACTCTCACAGCTATTGCTTTGAGATTCTGGAGCGGCGGGTCGACAAGCTTGCGG
CCGCACTCGAGCACCACCACCACCACCACCACCACTAA

eDHFR K6-tag(69K6) His-tag

**pT7-eDHFR(145K6)/pET41**

>Amino acid sequence
MISLIAALAVDRVIGMENAMPWNLPADLAWFKRNTLNKPVIMGRHTWESIGRPLPGRKNIILSSQPGTDDRVT
WVKSVDEAIAACGDVPEIMVIGGGRVYEQFLPKAQKLYLTHIDAEVEGDTHFPDYEPDDWESVFSEFHDADAK
KKKKKQNSHSYCFEILERRVDKLAAALEHHHHHHHH*

>DNA sequence
ATGATCAGTCTGATTGCGGCGTTAGCGGTAGATCGCGTTATCGGCATGGAAAACGCCATGCCGTGGAACCTGC
CTGCCGATCTCGCCTGGTTTAAACGCAACACCTTAAATAAACCCGTGATTATGGGCCGCCATACCTGGGAATC
AATCGGTCGTCCGTTGCCAGGACGCAAAAATATTATCCTCAGCAGTCAACCGGGTACGGACGATCGCGTAACG
TGGGTGAAGTCGGTGGATGAAGCCATCGCGGCGTGTGGTGACGTACCAGAAATCATGGTGATTGGCGGCGGTC
GCGTTTATGAACAGTTCTTGCCAAAAGCGCAAAAACTGTATCTGACGCATATCGACGCAGAAGTGGAAGGCGA
CACCCATTTCCCGGATTACGAGCCGGATGACTGGGAATCGGTATTCAGCGAATTCCACGATGCTGATGCGAAA
AAAAAGAAAAAGAAACAGAACTCTCACAGCTATTGCTTTGAGATTCTGGAGCGGCGGGTCGACAAGCTTGCGG
CCGCACTCGAGCACCACCACCACCACCACCACCACTAA

eDHFR K6-tag(69K6) His-tag

**pT7-K6-eDHFR/pET41**

>Amino acid sequence
MKKKKKKGSGASAGGGSGAGSGAISLIAALAVDRVIGMENAMPWNLPADLAWFKRNTLNKPVIMGRHTWESIG
RPLPGRKNIILSSQPGTDDRVTWVKSVDEAIAACGDVPEIMVIGGGRVYEQFLPKAQKLYLTHIDAEVEGDTH
FPDYEPDDWESVFSEFHDADAQNSHSYCFEILERRVDKLAAALEHHHHHHHH*

>DNA sequence
ATGAAAAAAAAGAAAAAGAAAGGCTCCGGTGCCAGTGCTGGTGGTGGCAGCGGTGCTGGTTCCGGCGCTATCA
GTCTGATTGCGGCGTTAGCGGTAGATCGCGTTATCGGCATGGAAAACGCCATGCCGTGGAACCTGCCTGCCGA
TCTCGCCTGGTTTAAACGCAACACCTTAAATAAACCCGTGATTATGGGCCGCCATACCTGGGAATCAATCGGT
CGTCCGTTGCCAGGACGCAAAAATATTATCCTCAGCAGTCAACCGGGTACGGACGATCGCGTAACGTGGGTGA
AGTCGGTGGATGAAGCCATCGCGGCGTGTGGTGACGTACCAGAAATCATGGTGATTGGCGGCGGTCGCGTTTA
TGAACAGTTCTTGCCAAAAGCGCAAAAACTGTATCTGACGCATATCGACGCAGAAGTGGAAGGCGACACCCAT
TTCCCGGATTACGAGCCGGATGACTGGGAATCGGTATTCAGCGAATTCCACGATGCTGATGCGCAGAACTCTC
ACAGCTATTGCTTTGAGATTCTGGAGCGGCGGGTCGACAAGCTTGCGGCCGCACTCGAGCACCACCACCACCA
CCACCACCACTAA

eDHFR K6-tag(69K6) His-tag
